# A telomere-to-telomere genome assembly of Zhonghuang 13, a widely-grown soybean variety from the original center of Glycine max

**DOI:** 10.1101/2023.09.27.559666

**Authors:** Anqi Zhang, Tangchao Kong, Baiquan Sun, Shizheng Qiu, Jiahe Guo, Shuyong Ruan, Yu Guo, Jirui Guo, Zhishuai Zhang, Yue Liu, Zheng Hu, Tao Jiang, Yadong Liu, Shuqi Cao, Shi Sun, Tingting Wu, Huilong Hong, Bingjun Jiang, Maoxiang Yang, Xiangyu Yao, Yang Hu, Bo Liu, Tianfu Han, Yadong Wang

## Abstract

Soybean (*Glycine max*) stands as a globally significant agricultural crop, and the comprehensive assembly of its genome is of paramount importance for unraveling its biological characteristics and evolutionary history. Nevertheless, previous soybean genome assemblies have harbored gaps and incompleteness, which have constrained in-depth investigations into soybean. Here, we present the first Telomere-to-Telomere (T2T) assembly of the Chinese soybean cultivar “Zhonghuang 13” (ZH13) genome, termed ZH13-T2T, utilizing PacBio Hifi and ONT ultralong reads. We employed a multi-assembler approach, integrating Hifiasm, NextDenovo, and Canu, to minimize biases and enhance assembly accuracy. The assembly spans 1,015,024,879 bp, effectively resolving all 393 gaps that previously plagued the reference genome. Our annotation efforts identified 50,564 high-confidence protein-coding genes, 707 of which are novel. ZH13-T2T revealed longer chromosomes, 421 not-aligned regions (NARs), 112 structure variations (SVs), and a substantial expansion of repetitive element compared to earlier assemblies. Specifically, we identified 25.67 Mb of tandem repeats, an enrichment of 5S and 48S rDNAs, and characterized their genotypic diversity. In summary, we deliver the first complete Chinese soybean cultivar T2T genome. The comprehensive annotation, along with precise centromere and telomere characterization, as well as insights into structural variations, further enhance our understanding of soybean genetics and evolution.

## Introduction

Soybeans (*Glycine max* [L.] Merr.), originating in China, hold a paramount position as one of the most crucial oil and protein crops. They contribute to more than a quarter of the protein utilized in both food and animal feed ^1, 2, 3, 4, 5^. It is widely acknowledged that the cultivated soybean emerged through the domestication of its wild annual progenitor, *Glycine soja*, around 5,000 years ago from the Yellow River Basin in temperate regions of China. This specific geographical range represents the greatest allelic diversity of soybean ^6, 7^. Subsequently, its distribution expanded northward to encompass high-latitude cold zones and southward to encompass low-latitude tropical regions. Therefore, the exploration of genetic resources within the origin region bears immense significance in advancing the global frontiers of soybean breeding.

“Zhonghuang 13”, a soybean cultivar meticulously developed and released by Chinese breeders in 2001, occupied the largest planting area in the first two decades of 21st century in China, and stood as a testament to advanced agronomic traits and remarkable adaptability to wide regions including Yellow River Basin, southern Northeast, and some parts of Northwest and South China ^8, 9^. In comparison to the widely recognized Williams 82 cultivar, “Zhonghuang 13” boasts heightened genetic diversity and ecological type of origin reign ^10^. Futhermore, “Zhonghuang 13” is an ideal variety in the breeding strategy called “Potalaization”, which allows breeding of novel widely adapted soybean varieties through the use of multiple molecular tools in existing elite widely adapted varieties ^7^.

Whole-genome sequencing of “Zhonghuang 13” has been previously conducted ^8, 9^. This approach enables the identification of crucial genes and genetic variants linked to favorable traits, thereby enhancing our comprehension of soybean breeding ^5, 11, 12, 13, 14, 15, 16^. Nonetheless, limitations inherent in second-generation sequencing, including inadequate coverage of the genome and challenges in precisely assembling and annotating repetitive genomic regions, such as telomeres and centromeres, have resulted in the persistence of over 1,000 gaps within the most recent soybean reference genome ^17, 18, 19, 20, 21^.

Telomere-to-Telomere (T2T) sequencing is a state-of-the-art genomic sequencing method that employs long-read sequencing platforms such as Pacific Biosciences (PacBio) or Oxford Nanopore Technologies (ONT) to obtain the comprehensive sequence from one telomere to another, encompassing highly repetitive regions, centromeres, and telomeres ^22, 23, 24^. This approach yields a complete and contiguous assembly of the entire genome ^23, 25^. T2T sequencing effectively overcomes the limitations associated with sequence gaps and assembly errors that are frequently encountered in whole-genome sequencing ^26^. Moreover, it offers enhanced resolution for the detection and characterization of large-scale SVs ^27, 28^. Consequently, T2T sequencing exhibits immense potential in expanding our knowledge of intricate genomes, such as soybeans, and driving advancements in breeding programs.

In this study, we utilized T2T sequencing to conduct *de novo* assembly of “Zhonghuang 13” genomes. By employing this innovative genomic assembly approach, we aim to deliver a fully covered soybean sequence encompassing 100% of the genome. This significant advancement will enhance our comprehension of the structure and functional significance of the soybean genome, and also provide a reference genome sequence for elite cultivar improvement.

## Results

### T2T assembly of the soybean ZH13 genome

Four types of sequencing data were initially produced for a single ZH13 sample, including PacBio Hifi reads (96.89 Gbp), ONT ultralong reads (96.63 Gbp), Illumina whole genome sequencing (WGS) (55.40Gbp) and Illumina high-throughput chromosome conformation capture (Hi-C) reads (106.4 Gbp). We only used the long reads (PacBio Hifi and ONT ultralong) to implement T2T assembly, and the short WGS and Hi-C reads were employed to assess assembly quality. The assembly was implemented by a pipeline based on multiple assemblers and in-house tools in three phases as following (a flowchart is in **Fig. 1**).

1) Draft assembly. At first, three set of contigs were independently produced by various assemblers, i.e., Hifiasm ^29, 30^, NextDenovo^31^ and Canu ^32^. Both of Hifiasm and NextDenovo used all the PacBio Hifi and ONT ultralong reads, and Canu used PacBio reads only. The 23 >1Mbp contigs produced by Hifiasm were employed as primary contigs and aligned to the current version of ZH13 reference ^33^ (termed as ZH13-ref-2019) by minimap2 ^34^. 22 of them can be colinearly aligned and 1 contig were aligned to two different chromosomes. We manually checked this split alignment and confirmed that it was a mis-assembly caused by Hifiasm. The contig was then divided into two. A 24-contigs draft assembly was then generated, which 17 of the 20 ZH13 chromosomes were covered by a single contig, 2 and 1 chromosomes have 2 and 3 contigs, respectively. The PacBio reads, ONT reads and the contigs produced by NextDenovo and Canu were aligned to the draft assembly for further refinement.

2) Assembly refinement. There were four remaining gaps in the draft assembly. Moreover, we also detected high- and low coverage regions (HCRs and LCRs) by an in-house script as they could be also mis-assembled regions. 43 HCRs and 2 LCRs were found. We searched the sequences of the HCRs by BLAST and the results indicated that all of them can be aligned to mitochondria, chloroplast, mRNAs or mobile elements. Thus, we realized that they could be not mis-assembly. However, plenty of read clippings were observed around the two LCRs, which indicated mis-assembles. Thus, the four gaps (gap1: CM010418.2: 18,024,780-18,025,280, gap2: CM010419.2:27,778,852-27,779,352, gap3: CM010421.2: 3,326,824-3,327,324 and gap4: CM010421.2: 40,056,171-40,056,671) and the two LCRs (LCR1: CM010409.2: 15,403,000-15,404,000 and LCR2: CM010427.2: 15,777,563-16,073,378) were refined with spanning NextDenovo and Canu contigs (refer to Methods for more detailed information). Gap2, gap4 and LCR1 were successfully reconstructed by two NextDenovo contigs and one Canu contig (**Supplementary Figs. 1-3**), respectively. However, Gap1, gap3 and LCR2 were turned to be HCRs (**Fig. 2-3, Supplementary Fig. 4**), indicating that the spanning contigs also cannot well-handle them. Highly repetitive sequences were found there, i.e., gap1 is full of LTR retrotransposons, while gap3 and LCR2 are rDNA arrays. Further, an in-house local assembly tool was employed to iteratively collect and tile the reads anchored to the corresponding regions to refine the assembly. The solved Gap1 is 467.3 Kbp long which mostly consists of gypsy and copia. Gap3 and LCR2 are about 4.15 Mbp and 414 Kbp long, respectively, having 545 48S rDNA copies and 1269 5S rDNA copies. We manually checked the read re-alignments to the three regions with IGV ^35^ and normal coverages were observed (**Fig. 2-3, Supplementary Fig. 4**). The genomic structures of all gaps were presented in the **Supplementary Fig. 5**.

**Fig. 1.**
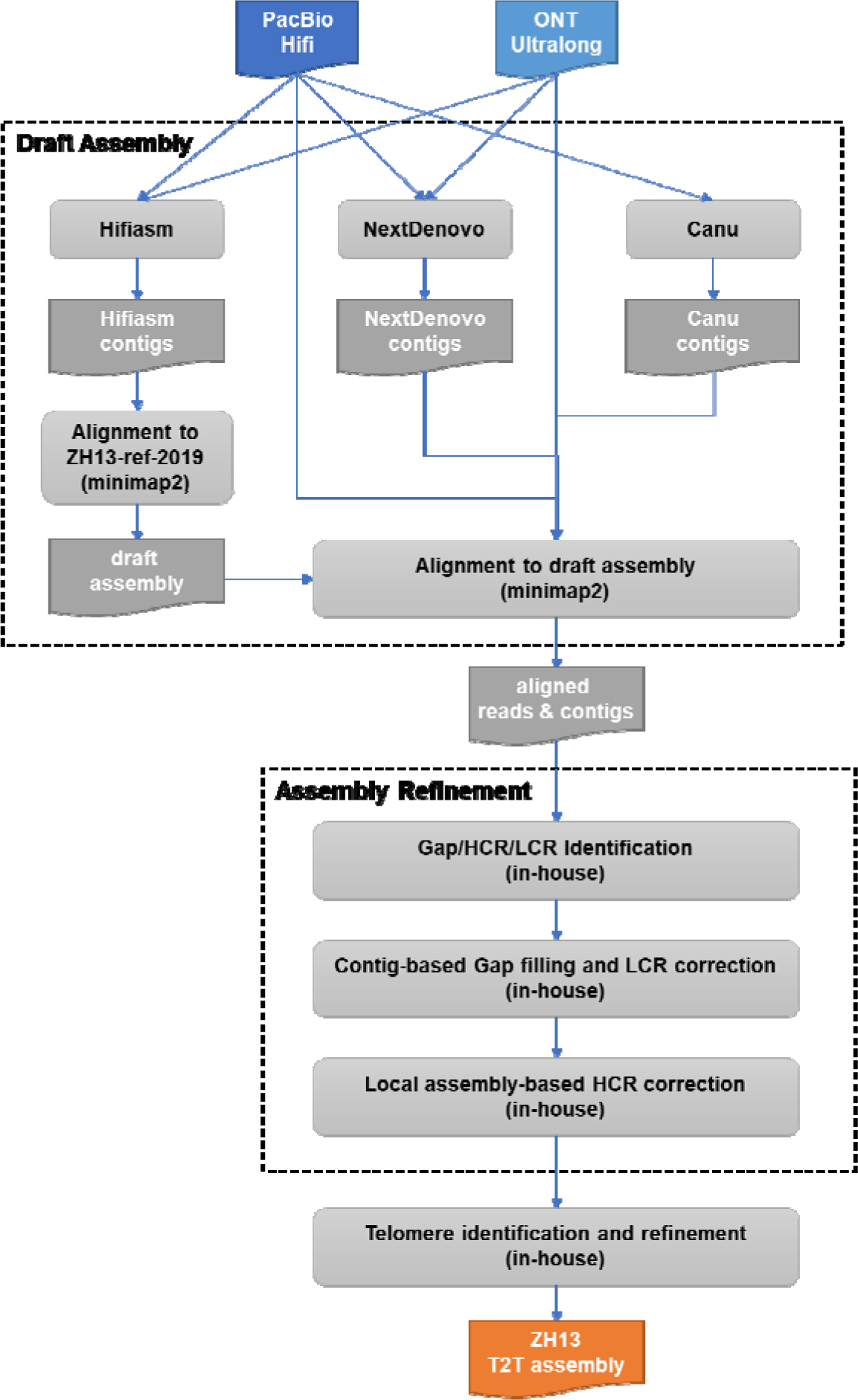
ZH13-T2T assembly pipeline.

**Fig. 2.**
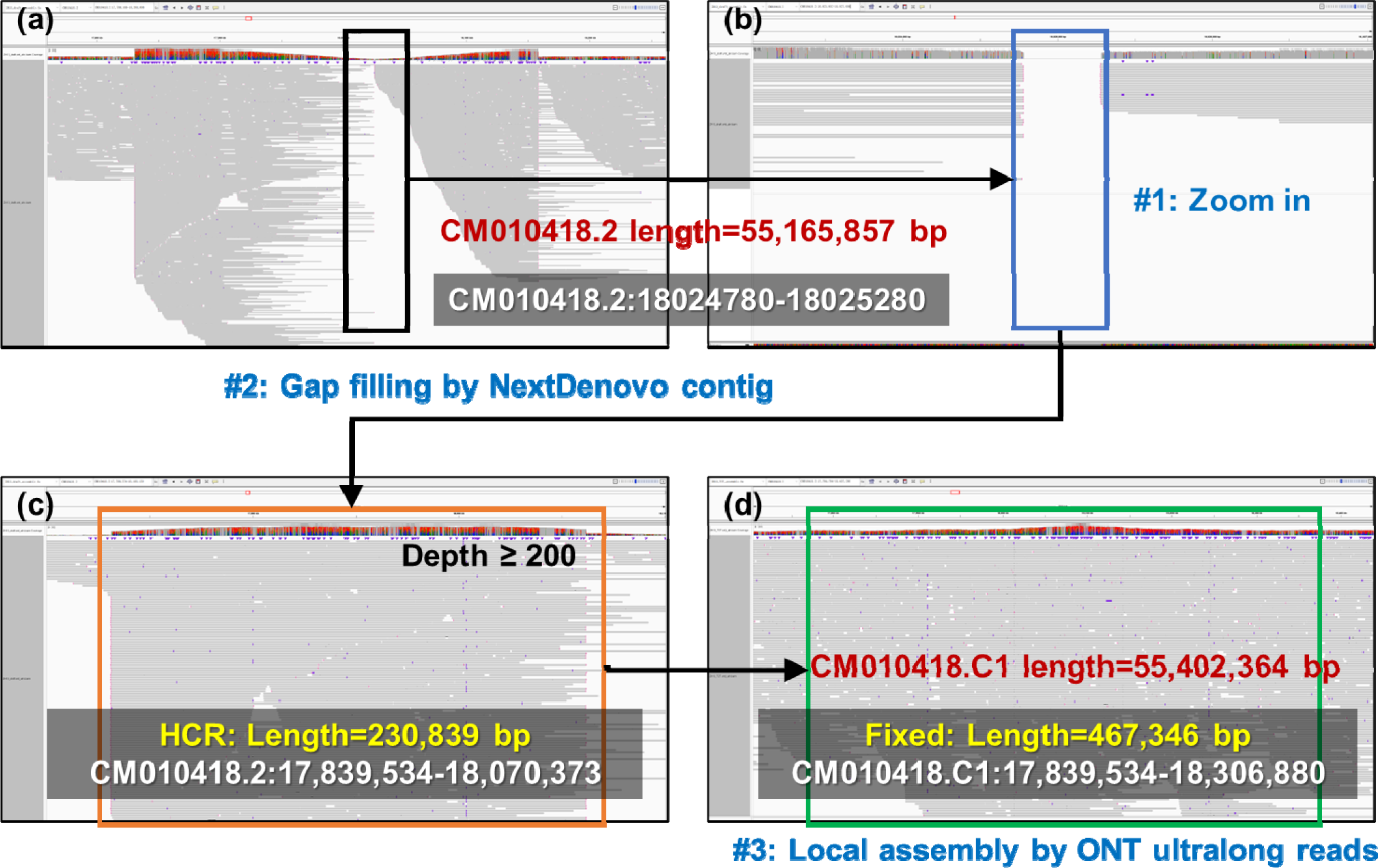
The filling of gap1. a) the IGV snapshot of ONT ultralong read alignment in the surrounding region of gap1 in ZH13-ref-2019; b) a zoom-in view of gap1; c) the IGV snapshot read alignment after NextDenovo contig correction (HCR was observed); d) the IGV snapshot of ONT ultralong read alignment after local assemply-based correction with anchored ONT ultralong reads (consecutive alignments and normal coverage were observed).

**Fig. 3.**
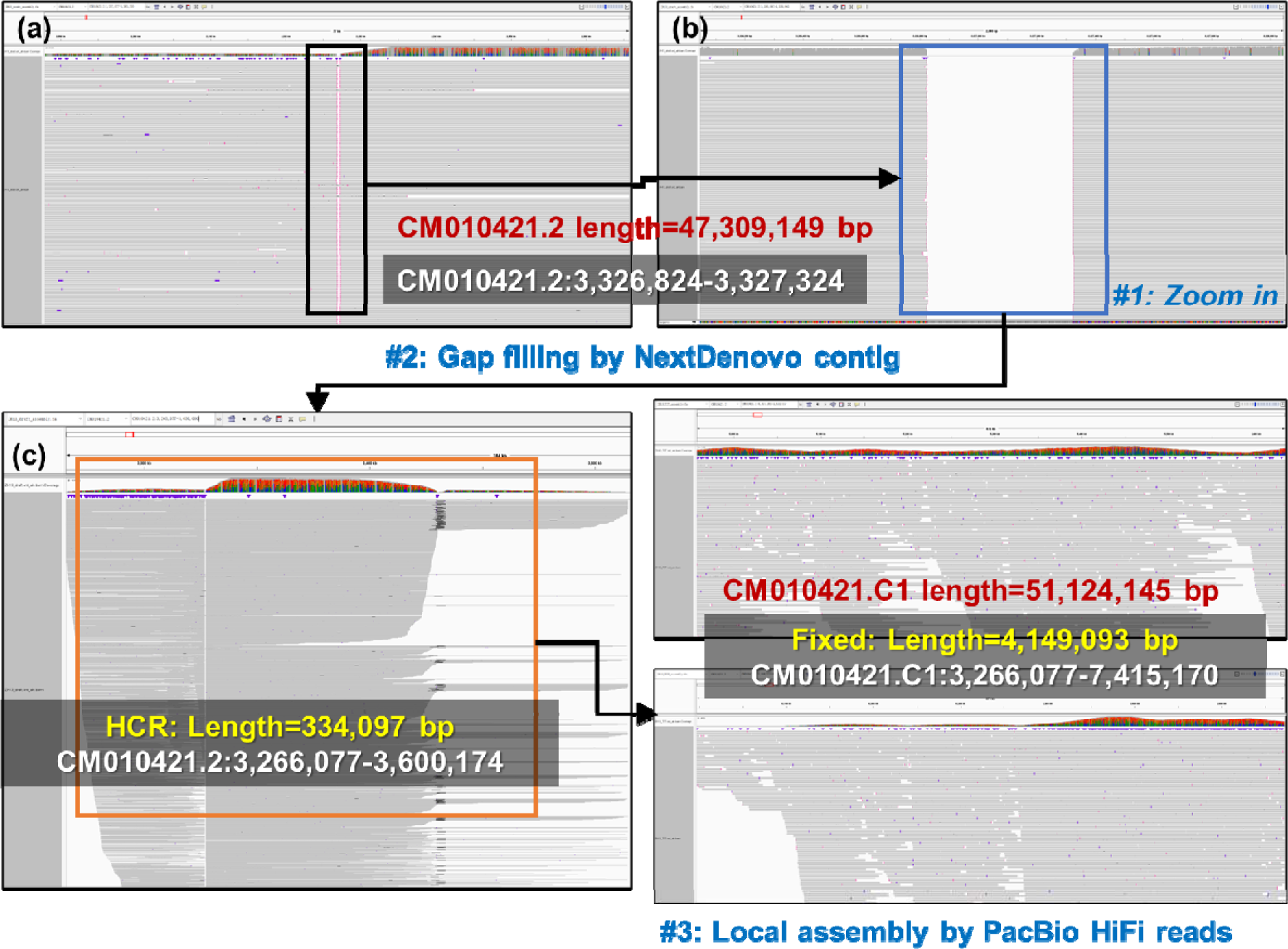
The filing of gap3. a) the IGV snapshot of ONT ultralong read alignment in the surrounding region of gap3 in ZH13-ref-2019; b) the IGV snapshot of ONT ultralong r NextDenovo contig correction (HCR was observed and 48S rDNA array was identified); c) the IGV snapshot of ONT ultralong read align embly-based correction with anchored reads, consecutive read alignments were observed. Moreover, by manually checking the alignment de many reads having very low MAPQ, i.e., each of the reads had multiple candidate mapping positions and cannot be confidently aligned. Ov ge was proved by considering all the primary and secondary alignments of the reads.

3) Telomere identification and refinement. 37 telomeres were identified from 17 chromosomes of the draft assembly. We further checked the contigs produced by NextDenovo, Canu and Hifiasm (using Hifi reads only), and reconstructed the three missing telomeres, i.e., CM010417.C1 (downstream, 3831bp), CM010418. C1 (upstream, 5889 bp) and CM010423.C1 (downstream, 9764 bp). Thus, all the 40 telomeres were recovered with an 8449 bp median length.

Finally, a complete genome of ZH13 (termed as ZH13-T2T, **Fig. 4**) was generated whose total length is 1,015,024,879 bp (no gap, N50: 52,033,905 bp). The quality of the assembly was evaluated by various metrics and four issues are observed as following. Firstly, the complete BUSCO metric ^36^ (99.8%, lineage dataset: embryophyta_odb10) suggests its high completeness. More importantly, all the 393 gaps of ZH13-ref-2019 have been filled. Secondly, Illumina WGS read-based Merqury’s Qv metric ^37^ reaches 46.441, suggesting that it also achieves high base-level accuracy. Thirdly, it is observed from the Hi-C map (**Supplementary Fig. 6**, generated by Juicerbox ^38^) that strong interactions are concentrated along the diagonal, indicating that no obvious mis-assembly can be discovered from the view of Hi-C data. Fourthly, with careful detection and correction of HCRs and LCRs, the coverages of PacBio Hifi and ONT Ultralong reads are nearly uniform along the whole ZH13-T2T genome, also suggesting that the assembly could be free of mis-assembly.

**Fig. 4.**
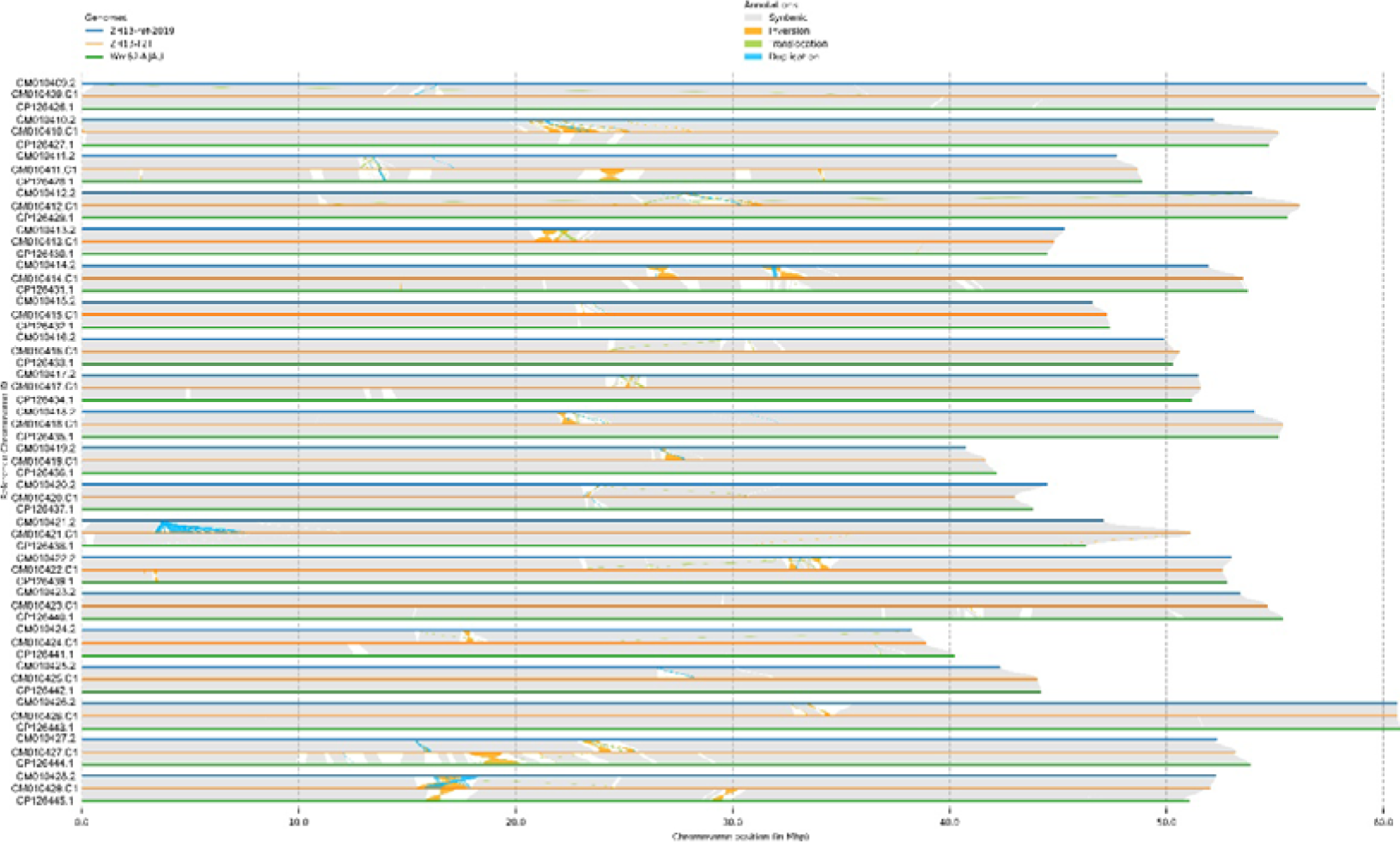
The ZH13-T2T assembly and its comparison to ZH13-ref-2019 and Wm82-NJAU. The orange, blue and green lines indicate ZH13-T2T, ZH13-ref-2019 and Wm82-NJAU, respectively. Gray and blank blocks between various genomes indicate syntenic regions and NARs. Inversions, translocations and duplications are marked by filled orange, green and blue curves.

### Genome-wide comparison to ZH13-ref-2019 and the T2T assembly of Wm82

We compared ZH13-T2T with ZH13-ref-2019 by SyRI ^39^(**Fig. 4**). Most of the ZH13-T2T chromosomes of are longer, mainly due to the filled gaps. There are also 421 >5K bp not-aligned regions (NARs, 16.3 Mbp in total), indicating that the corresponding local sequences are quite different. Meanwhile, SyRI also identified 112 structure variations (SVs), i.e., 30 inversions, 15 translocations and 67 duplications. Most of them are in the NARs and highly complex, i.e., the combinations of multiple inversions and duplications.

We further aligned the ONT ultralong reads of ZH13-T2T to ZH13-ref-2019 (by minimap2). The local alignments in the NARs showed concentrated and extremely complex SV signatures, i.e., plenty of large clippings, split alignments and abnormal local coverages, especially for those SV-surrounding regions (an example is in **Supplementary Fig. 7**). On the contrary, colinear alignments with normal coverages were observed from the corresponding regions of ZH13-T2T, suggesting no obvious SV signature there. We also investigated the alignments on the NARs without SVs and similar results were observed (**Supplementary Fig. 8**). Under such circumstance, we realize that although the different donor samples potentially have divergences in genomic sequences, there could be also a number of mis-assemblies in ZH13-ref-2019, possibly due to the limitation of its sequencing data. Moreover, considering the consecutive read alignments on ZH13-T2T, the mis-assemblies should have been largely resolved.

A comparison between ZH13-T2T and the newly published T2T assembly of Wm82 (Wm82-NJAU) ^40^ was also conducted. Mainly, 167 NARs (23.02 Mbp in total) and 30 SVs (16 inversions, 7 translocations and 7 duplications) were identified. To investigate the NARs and SVs, we also downloaded the PacBio Hifi and ONT ultralong datasets of Wm82-NJAU and re-aligned them to the genome. In a large proportion of the investigated regions, both of ZH13-T2T and Wm82-NJAU have normal read alignments (**Supplementary Fig. 9**), suggesting the differences between the two soybean genomes. However, in some of the NARs, abnormal read alignments can still be found from Wm82-NJAU but not for ZH13-T2T (**Supplementary Fig. 10**). Moreover, we also checked the local read coverages along the whole Wm82-NJAU genome and found tens of HCRs and LCRs.

### Genome annotation and gene prediction

Complete T2T assembly of 20 chromosomes revealed that approximately 57.07% of the soybean genome consisted of annotated repeating elements. Among these elements, retrotransposons accounted for 38.16% (comprising 0.12% SINEs, 1.58% LINEs, and 36.47% LTR elements), while DNA transposons accounted for 6.72% (**Table 1, Fig. 5**). Furthermore, we detected 3.64 Mb of microsatellites, 11.58 Mb of minisatellites, 11.44 Mb of satellites, 0.41 Mb of 5S rDNAs, and 4.16 Mb of 48S rDNAs (**Table 2**). Collectively, these tandem repeats constitute 2.63% (26.65 Mb) of the soybean genome, which significantly surpasses the 1.03% (10.54Mb) observed in the Zhonghuang 13 reference sequence. Additionally, the intergenic spacer (IGS) region of 5S rDNA is approximately 220 bp in length, while the 5S region itself spans approximately 110 bp (**Fig. 6**). There is a partial overlap region of around 1533 bp between 5.8S rDNA and 28S rDNA. Based on the analysis of 1 Indels, the 48S rDNA has been categorized into 2 distinct genotypes. Furthermore, an examination of 13 SNPs and Indels has led to the classification of the 5S rDNA into 32 different genotypes.

**Fig. 5.**
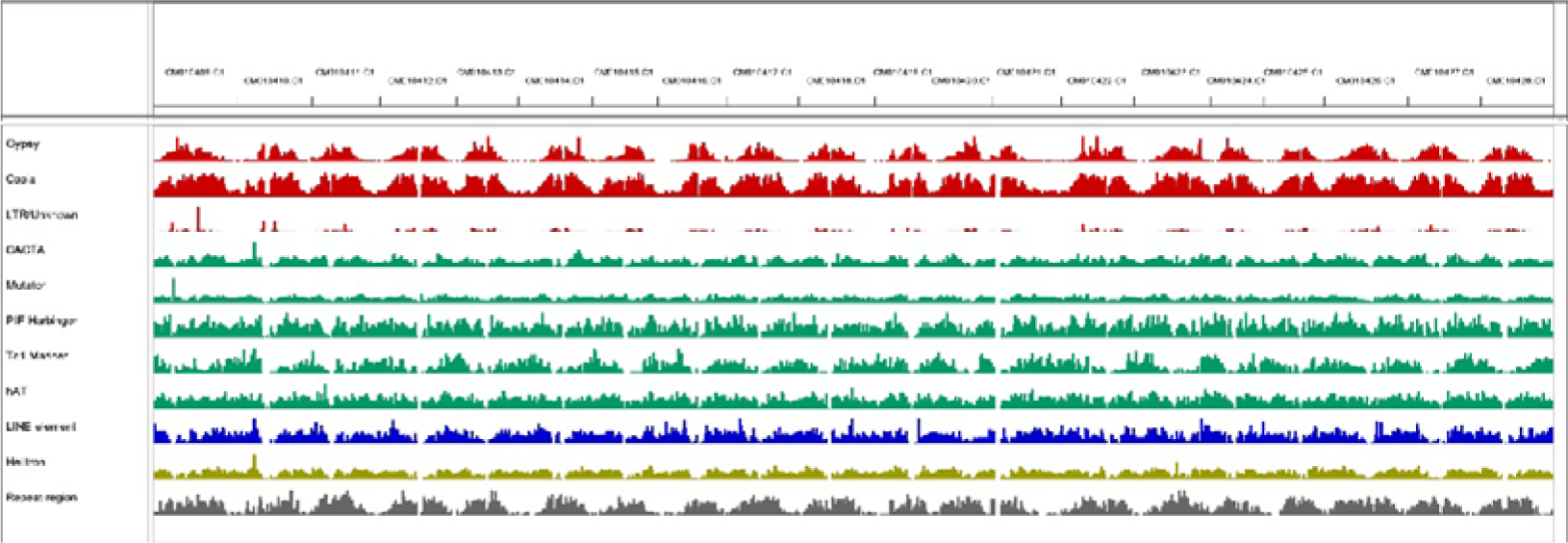
Distribution of TE in the ZH13-T2T genome.

**Fig. 6.**
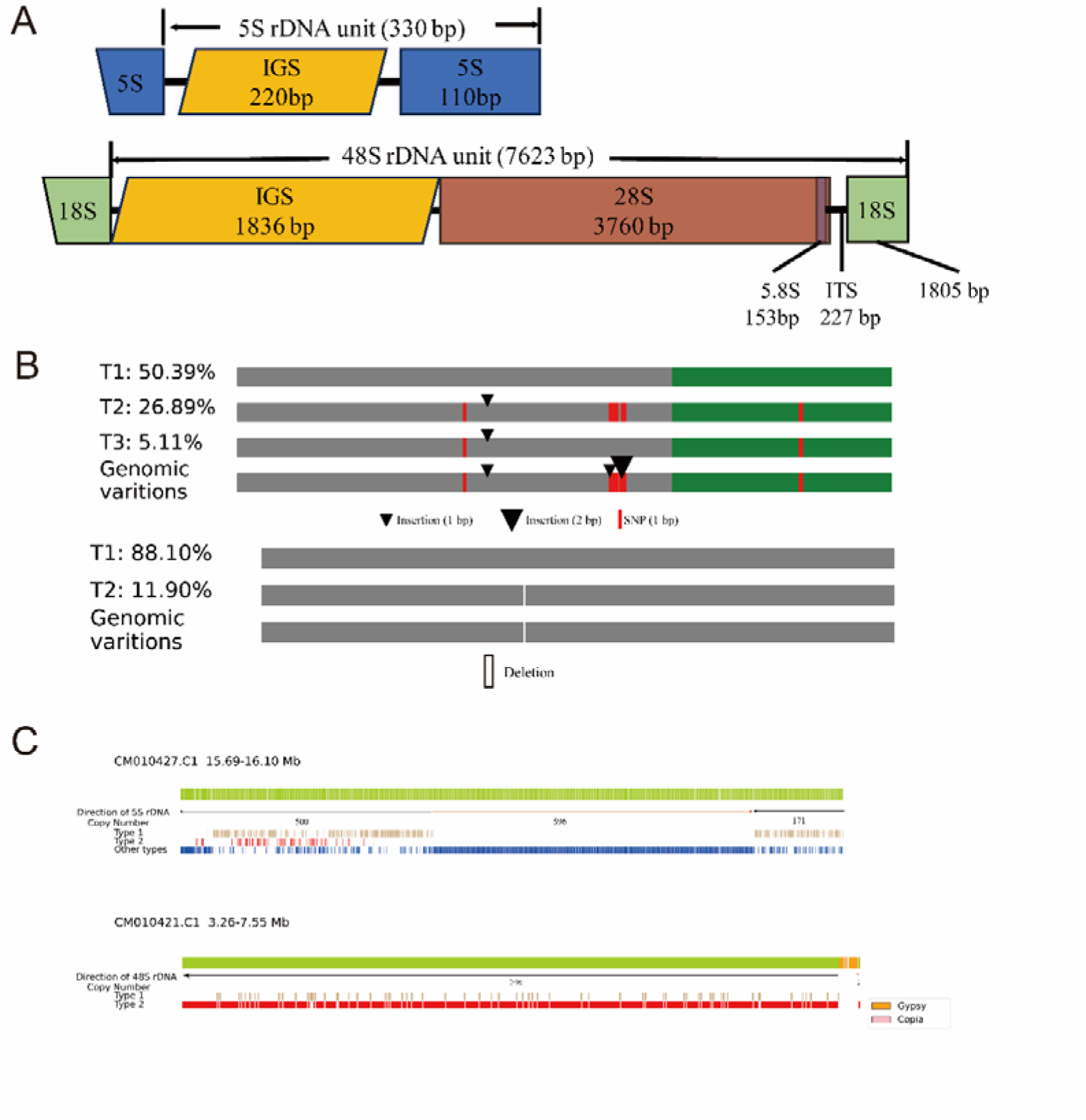
Genome structure of 5S and 48S rDNA arrays. (A). Sequence structure of a typical 5S (up) and 48S rDNAs (down) repeat unit. IGS, intergenic spacer region; ITS, internal transcribed spacer region. (B). Variations of the most abundant genotypes of 5S rDNAs (up) and 48S rDNAs (down). (C). Genome structure of rDNAs (up) and 48S rDNAs (down).

**Table 1.**
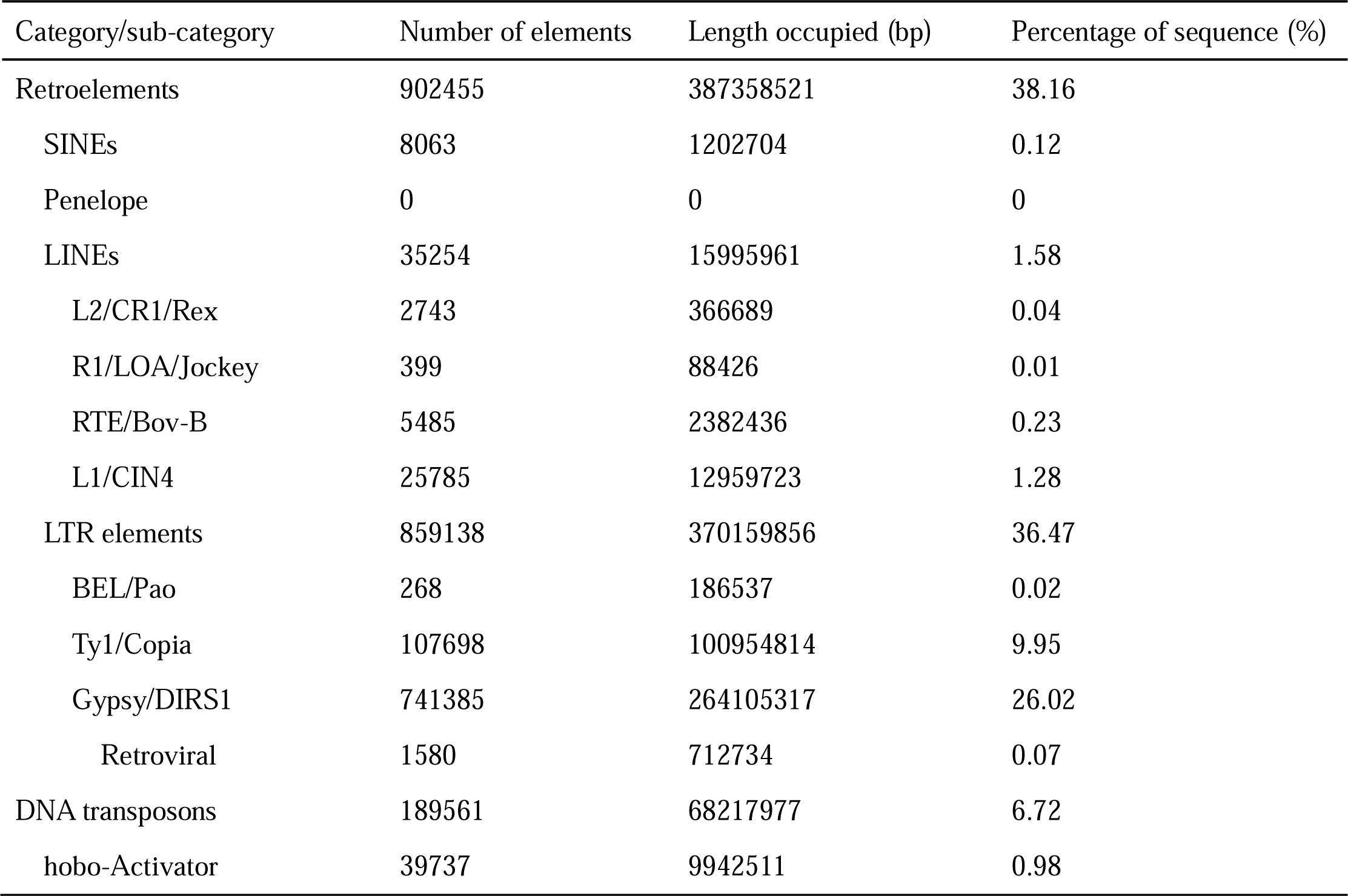
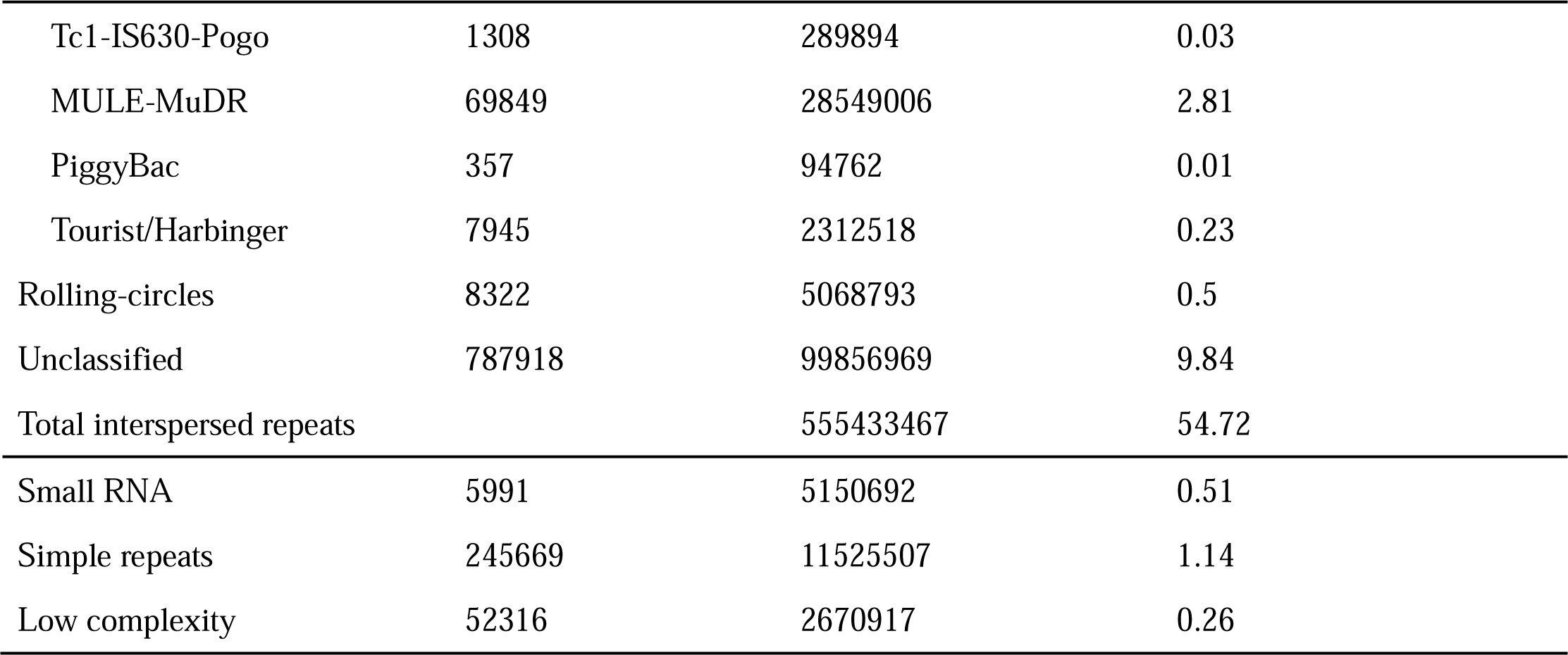
Statistics of repetitive elements in ZH13-T2T.

**Table 2.**
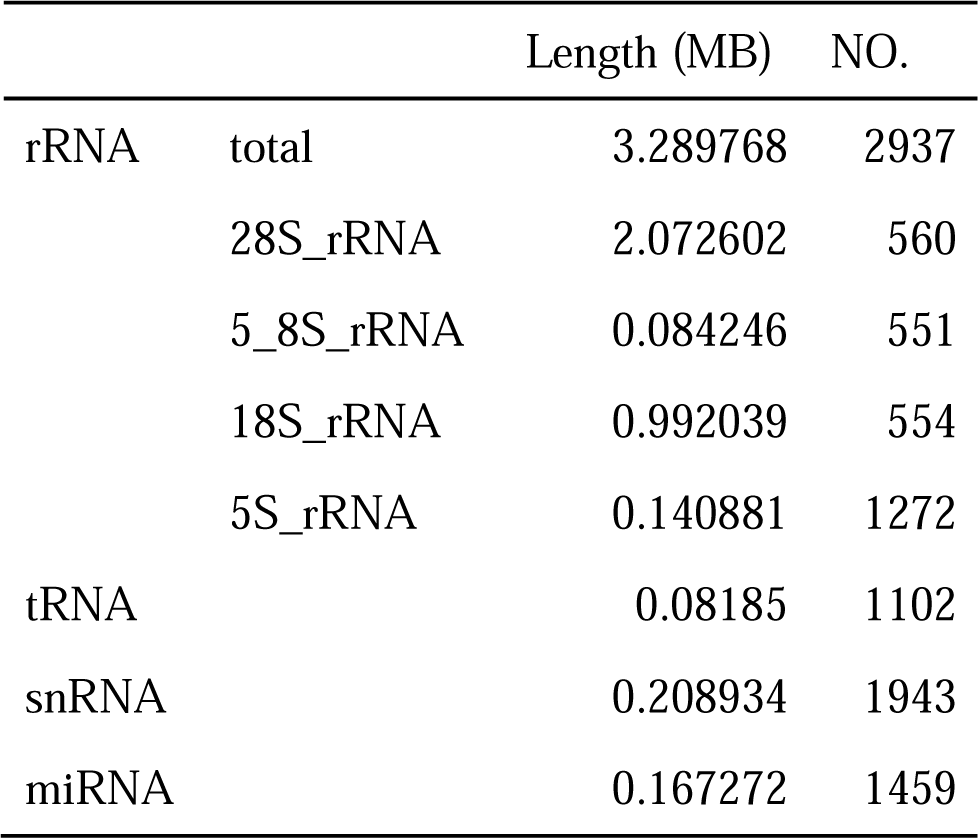
Summary of non-coding RNAs.

The annotation of the ZH13-T2T genome was performed using the Augustus software for ab initio annotation. After excluding transposon genes and applying a gene filtering process, a total of 50,564 high-confidence protein-coding genes were obtained. Notably, in comparison to ZH13-2019, we identified 707 novel genes within the gap regions. The results of the Gene Ontology (GO) function and Kyoto Encyclopedia of Genes and Genomes (KEGG) pathway enrichment analysis obtained through KOBAS (Knowledgebase for Ontology-based Functional Annotation and Analysis) reveal significant enrichments for newly discovered genes in various biological pathways. Specifically, the novel genes were involved in various biological processes, cellular components, and molecular functions, ranging from cell surface to nucleus, including negative regulation of transcription, DNA-templated (p=7.52E-05), protein autophosphorylation (p=0.0003), positive regulation of transcription by RNA polymerase II (p=0.0012), and mRNA export from nucleus (p=0.0094) (**Supplementary Fig. 11**). In the context of KEGG, these genes participated in phenylalanine, tyrosine and tryptophan biosynthesis (p=0.0004), fatty acid biosynthesis (p=0.0007), phosphatidylinositol signaling system (p=0.0046), fatty acid metabolism (p=0.0048), RNA transport (p=0.0105), and the MAPK signaling pathway - plant (p=0.0156) (**Supplementary Fig. 12**).

In the previous gap regions, we observed the highest number of newly discovered genes within the 14.84-17.73Mb region of chromosome CM010421.C1, totaling 135 new genes. Furthermore, our analysis identified 42,668 TEs, 300 GmCent-1 elements, and 586 GmCent-2 elements within these gap regions. Notably, the CM010420.C1 chromosome’s 30.54-32.38Mb region contained the highest count of TEs, with a total of 3,416 TEs. In the CM010413.C1 chromosome’s 20.40-24.24Mb region, we observed the highest number of GmCent-1 elements, amounting to 131, and in the CM010419.C1 chromosome’s 25.51-27.93Mb region, the highest count of GmCent-2 elements, totaling 64 (**Supplementary Fig. 5**).

### Detection of centromere

The centromere, a crucial component of chromosome structure, consists of highly repetitive heterochromatin and plays a vital role in ensuring accurate chromosome segregation. In plants, the centromere region is characterized by an abundance of retrotransposon and tandem repeats ^41^. Investigating the potential functions of the centromere in genome evolution and chromatin assembly holds significant importance. However, current genome sequencing approaches face challenges in fully assembling the repetitive sequences within the centromere region. Here, we employed the Tandem Repeat Finder (TRF) tool to identify repeat monomers within the ZH13-T2T genome that likely constitute the centromere. Consistent with previous studies, we found a large number of tandem repeats of 91 and 92bp in length, complementing the gaps in TE of centromere region (**Fig. 7A-D, Supplementary Fig. 13**) ^42^. The heat map shows high similarity of sequences in the centromere region, indicating that the centromere region is highly tandem repetitive (**Fig. 7E**).

**Fig. 7.**
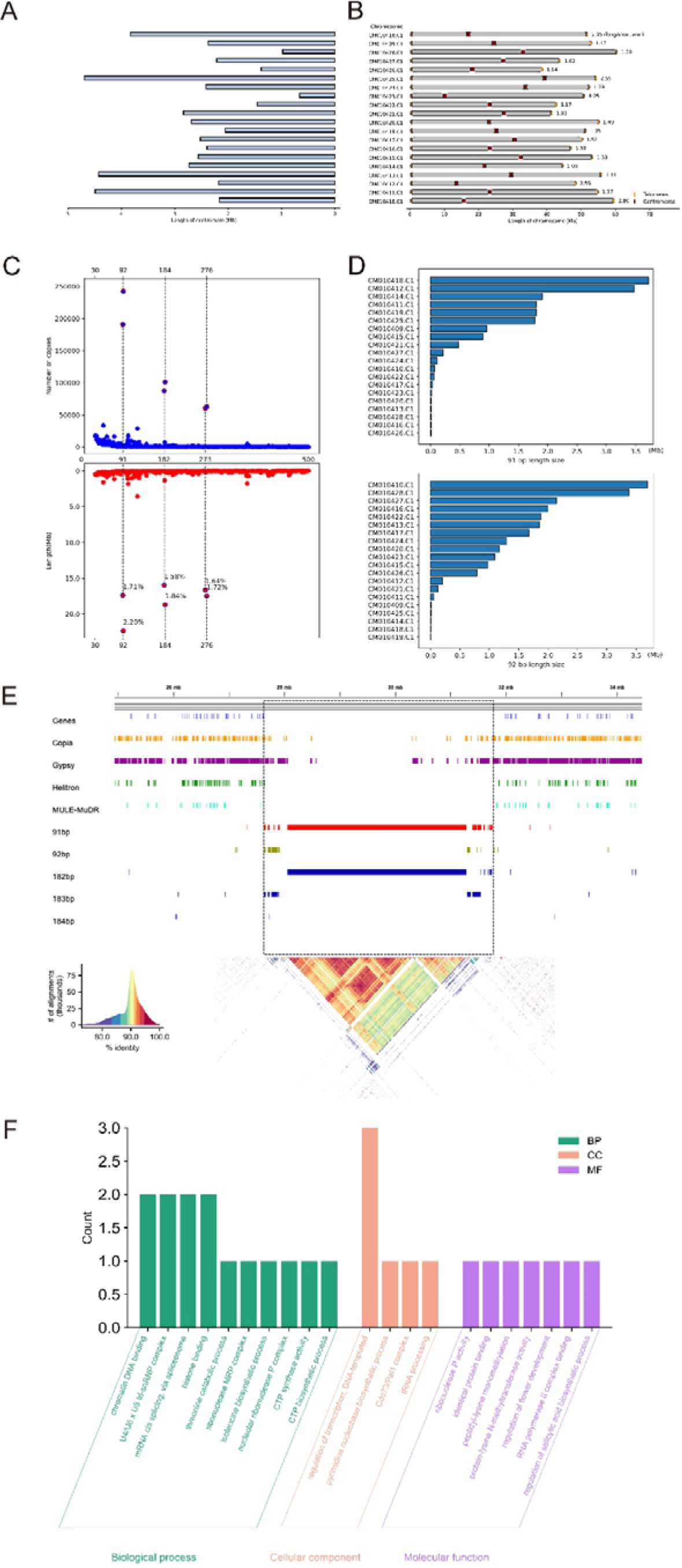
Genomic structure of telomere and centromere. (A) The length of the centromere. (B) The location of centromere in the genome. (C) The number of copies in the genome of tandem repeats with lengths of 91 and 92bp. (D) The total length of tandem repeats of 91bp and 92bp in each chromosome. (E) The distribution and degree of correlation of various sequences in the centromere region are represented by heat maps. (F) Enrichment of genes in centromere region.

Our findings indicate that the average length of the 20 centromeres analyzed is 2.40 Mb, with the longest centromere observed on CM010410.C1 (4.42 Mb) and the shortest on CM010421.C1 (0.66 Mb) (**Fig. 7A**). Notably, no significant correlation was observed between centromere length and chromosome size. Furthermore, the relative positions of centromeres varied among different chromosomes, with the minimum L/S ratio (long arm length/short arm length) recorded as 1.02 (CM010415.C1) and the maximum L/S ratio as 2.95 (CM010423.C1). A total of 8 genes were identified in the centromeric region of the soybean T2T genome. These genes were mainly enriched in chromatin DNA binding, mRNA cis splicing via spliceosome, histone binding, basal transcription factors, spliceosome, and pyrimidine metabolism (**Fig. 7F**).

On average, centromere sequences were composed of 96.0% centromere satellite DNA (CentC), centromere retrotransposons (CRM), and other non-CRM Gypsy retrotransposons. The proportions of these components varied significantly across different centromeres, ranging from 0.0% to 73.3% for GmCent-1, 0.0% to 90.4% for GmCent-2, 0.0% to 2.2% for CRM, and 7.3% to 68.2% for other non-CRM Gypsy retrotransposons (**Supplementary Fig. 14**). Almost all centromeres were rich in CentC, and there are no CentC-poor centromeres.

## Discussion

Cultivated soybeans originated in China, it has undergone strict genetic bottlenecks during domestication, resulting in accessions from origin region possibly exhibiting high genetic diversity. In this context, the cultivar “Zhonghuang 13,” developed in 2001 through meticulous breeding efforts by Chinese scientists, stands as a testament to the advancement of soybean agronomy and adaptability ^8, 9^. Derived from parent cultivars originating in the Yellow River Basin, “Zhonghuang 13” boasts not only heightened genetic diversity but also a distinct ecological origin compared to the widely recognized Williams 82 cultivar. While whole-genome sequencing efforts of “Zhonghuang 13” have been previously undertaken, limitations inherent to second-generation sequencing technologies, such as incomplete genome coverage and challenges in assembling and annotating repetitive genomic regions, have hindered the attainment of a comprehensive reference genome.

Herein, we generate ZH13-T2T, the first T2T genome assembly of Chinese soybeans. With its unprecedented completeness and high quality, the assembly provides a superior reference genome, as well as a new opportunity to comprehensively decode and deeply understand the complex repeats in soybean genomes, which is invaluable to the society for cutting-edge plant genomics studies. Moreover, as the most planted soybean cultivar in China, the ZH13-T2T genome is also a valuable resource to molecular inbreeding.

Although efforts have been made, it is still a non-trivial task to implement high quality T2T genome assembly, as the employed assembly tools could still have bias and lead to mis-assemblies, while the read length could be limited to solve those extremely long repeats. During the generation of ZH13-T2T, we used several tailored approaches guarantee assembly quality.

One is the use of multiple assemblers to take their advantages and reduce bias. More precisely, the three sets of contigs independently produced by the Hifiasm, NextDenovo and Canu played different roles. Overall, the Hifiasm contigs reconcile accuracy and continuity, which were used as primary contigs. Some of NextDenovo contigs have even higher ability to span long repetitive regions so that they were employed to fill the unsolved regions. The Canu contigs are usually shorter due to the limited length of Hifi reads, however, they are accurate and useful to reveal the elements of difficult repetitive regions such as retrotransposon- or rDNA-rich loci. Moreover, Canu also has good performance in telomere regions. Thus, they also played an important role to guide gap filling, LCR correction and telomere refinement.

Another one is the monitoring of read coverage, which is effective to prevent mis-assembly. Theoretically, a perfect assembly should have uniform read coverage along the whole genome, especially with the low GC-bias of long read sequencing. Thus, abnormal read coverage is a good indicator to mis-assembly. During ZH13-T2T assembly, we used local read coverages to conduct quality control all the way, which not only helps to detect mis-assemblies, but also guide to correctly reconstruct the sequences of gaps, LCRs and HCRs.

Long repeats are still difficult to solve in practice, even if ONT ultralong data is available. We used an in-house tool to carefully collect and align anchored reads to iterative infer those extremely long repetitive sequences, with the guidance of the inherent sequence divergences and read coverages in local regions. The tool is able to improve the assembly in long repetitive regions, especially with known elements. However, it is still an open problem to develop more effective and generic tools to solve long repeats with limited read length.

The meticulous analysis of the previous gap regions within the soybean genome holds significant implications for soybean breeding. It is well established that soybean is a quintessential short-day plant and, as such, inherently exhibits sensitivity to photothermal conditions, particularly with regard to photoperiod. The responses to these photothermal conditions play a pivotal role in determining the soybean’s capacity for growth, development, yield formation, and its ability to thrive across varying geographical latitudes ^4, 43^. Among these responses, flowering time and maturity stand out as the most influential factors dictating the geographical adaptability of soybean. Genetic investigations have identified no less than 11 loci, denoted as E1 through E4, E6 through E11, and J, which actively participate in the photoperiodic regulation of flowering time and maturity in soybean. To date, only two of these loci, E7 and E8, remain elusive in terms of cloning. E7 is mapped to chromosome 06 and is situated approximately 6 cM apart from E1 (Glyma.06G207800, Chr06:20207076-20207940), representing a distinct photoperiod-related gene distinct from E1 ^44^. Meanwhile, E8 is localized on chromosome 04 in close proximity to two homologous genes, E1La (Glyma.04G156400, Chr04:36758124-36758770) and E1Lb (Glyma.04G143300, Chr04:26120010-26120532). Remarkably, despite extensive efforts, the E7 and E8 loci have remained uncloned. Our T2T-ZH13 reference genome annotation on chromosomes 04 and 06 has unveiled a multitude of novel genes, offering promising prospects for the cloning of E7 and E8 by providing fresh molecular targets.

Additionally, within these previously unexplored gap regions, we have annotated 505 novel genes. Through KEGG and GO enrichment analysis, it has come to light that these genes are implicated in a diverse array of biological pathways, encompassing various aspects of biosynthesis, metabolism, and cellular signal transduction. Simultaneously, they play pivotal roles in several biological functions, including gene regulation and RNA processing. These newfound genes hold the promise of serving as novel molecular targets for subsequent Genome-wide association studies (GWAS) and for the validation of related gene functions.

Beyond assembly improvements, “ZH13-T2T” enhances the soybean genome’s annotation, particularly regarding repetitive elements, centromeres, and telomeres. Repetitive elements play a significant role in genome evolution and gene regulation ^45, 46, 47, 48^. We found that repeat elements constituted a significant portion of the genome, with retrotransposons, particularly LTR elements, being predominant. We also identified the presence of abundant satellites and rDNAs. The T2T genome exhibits a notable expansion in repetitive sequences compared to previous assemblies. Moreover, the identification and characterization of centromeres and telomeres within “ZH13-T2T” offer valuable insights into the organization and maintenance of chromosomal integrity. Soybean, a relic of ancient tetraploid plant evolution, has undergone two significant whole-genome duplication or polyploidization events ^49^. Within soybean genome, two distinct centromeric repeat classes exist, and their distribution is notably uneven, signifying the presence of two subgenomes in soybean. It is plausible that these subgenomes may have originated from the hybridization of two now-extinct plants with 2n=20 chromosomes, followed by a subsequent partial homogenization of one centromeric repeat class by the other. Research findings suggest that the CentGm-1 ancestor possessed a higher chromosome count compared to the CentGm-2 ancestor ^42^.

In conclusion, the ZH13-T2T genome represents a significant advancement in soybean genomics. The comprehensive genome annotation, identification of key genomic features, and insights into structural variations contribute to our understanding of soybean genetics and evolution. This high-quality reference genome will serve as a valuable resource for future studies in biology and practices in molecular breeding of soybean

## Methods

### Plant material preparation and genome sequencing

The soybean seeds of *Glycine max*, cv. Zhonghuang 13 (ZH13) were from the Institute of Crop Science, Chinese Academy of Agricultural Sciences. Four soybean seeds were planted in 10-litre pots on June 5, 2023, and grown outdoors under natural conditions in Beijing, China (39.95°N, 116.32°E). On Day 20 after emergence (VE), the fresh young leaf tissue was collected and frozen immediately in liquid nitrogen for DNA extraction. The High Molecular Weight (HMW) DNA extraction was performed using the modified cetyltrimethylammonium bromide (CTAB) method and lagre fragments (>100 kb) were separated using the SageHLS HMW library system. The standard libraries were constructed for subsequent sequencing. For PacBio HiFi sequencing, the library was constructed using SMRTbell Express Template Prep Kit. For ONT ultra-long sequencing, the library was created using SQK-ULK001 kit. For WGS sequencing, the library was created using NEBNext Ultra II DNA Library Prep Kit. For Hi-C sequencing, fresh leaves were fixed in 4% (vol/vol) formaldehyde after grinding with liquid nitrogen. Cell lysis, chromatin capture and digestion, and DNA quality chcek were performed according to the modified methods from ^50^. The library was created using NEBNext Ultra II DNA Library Prep Kit. A PacBio Revio sequencer was used to produce a 96.89 Gbp Hifi datasets (mean read length: 100.7 kbp) and a PromethION 48 sequencer was used to produce a 96.63 Gbp ultralong dataset (mean read length: 100.7 kbp). Moreover, the 150 bp pair-end WGS (55.40Gbp) and Hi-C (106.4 Gbp) datasets were produced by an Illumina Novaseq 6000 sequencer.

### Draft genome assembly by PacBio Hifi and ONT ultralong reads

The PacBio Hifi reads were input to Hifiasm ^29^ (version: 0.19.5-r590, default parameters) to generate a Hifi graph, and the ONT ultralong reads were then aligned to the graph to produce choromosome-level contigs ^30^. For NextDenovo ^31^ (version: 2.5.2, parameters: read_cutoff = 1k, genome_size = 1g), the ONT ultralong reads were input at first to produce initial contigs which were further polished by NextPolish ^51^ (version: 1.4.1, parameters: -x map-hifi -min_read_len 1k -max_depth 100) with input PacBio Hifi reads. For Canu ^32^ (version: 2.2, parameters: genomeSize=1g), only the PacBio Hifi reads were input to produce Hifi-only contigs.

The long (>1 Mbp) Hifiasm contigs were employed as primary contigs at first. Moreover, BLAST was employed to check the 851 <1 Mbp Hifiasm contigs. We found out that 759, 56 and 30 of the short contigs can be aligned to chloroplast, mitochondrion and rDNA, and others can also be aligned to repeats of soybean genomes as well. Thus, they were filtered out and the primary contig set did not update since we would like to build a concise draft assembly and solve potential gaps and mis-assemblies with the supplement of other assemblers. The primary contigs were then aligned to ZH13-ref-2019 by minimap2 ^34^ (version: 2.26-r1175, parameters: -x asm5 -f 0.02) to determine their orders and orientations. A draft ZH13 assembly was then generated and all the PacBio reads, ONT reads, NextDenovo contigs and Canu contigs were aligned to it by minimap2 (parameters: -x map-pb -r 1000; -x map-hifi -r 1000; -x map-ont -r 10000; -x asm5 -f 0.02) for further processing.

### Refinement of draft assembly

An in-house script was used to divide the draft assembly into 10 Kbp sliding windows and scan the read coverages to detect HCRs and LCRs. Herein, an HCR (LCR) is defined as a window whose coverage is >200 (<30). The local sequences of HCRs and LCRs were then searched by BLAST to check their homologies as an evidence of mis-assembly or not. Moreover, the numbers and positions of read clippings were also investigated as they are more important signatures to discover mis-assemblies. Since the HCR sequences can be aligned to mitochondria, chloroplast, mRNAs or mobile elements, their high coverages could be not due to mis-assembly, but plausibly the affection of the reads from those elements as well as the aligner’s own strategy to handle repetitive reads. So, they were not considered for correction.

The two LCRs and four remaining gaps in the draft assembly were then reconstructed in two steps. Firstly, we collected the NextDenovo and Canu contigs which can span those regions. The local sequences were then replaced by corresponding contigs with the guidance of nearby anchors. The reads were also re-aligned after the reconstruction to re-check local coverages. Moreover, we selected the contig leading to a local coverage closest to the mean coverage of the whole genome, if multiple candidates exist. It is worthnoting that this approach can either solve the LCRs/gaps or turn them to HCRs, since the employed contigs can at least reflect most of the elements existed in local regions, even if the copy numbers are incorrect and/or some of the local sequences are still absent.

For the still unsolved HCRs, we used an in-house tool to implement local assembly. Given an HCR, the tool collected the reads harbored or anchored to that region at first (termed as active reads) and iteratively assembles them. In each iteration, the tool separately tries each of the active reads to extend the local sequence from the region boundary, and aligns other reads to the extended sequence (by BLAST). If there are enough reads being aligned with high scores, the HCR is updated and a number (relative to the mean read coverage of the whole genome) of highest scored reads are removed. The procedure continues until the contig reaches the other boundary of the HCR, or no active read remains. The produced contig is then integrated into the genome with manual curation and read coverage checking.

### Telomere identification and refinement

The 7-mer repeats (CCCTAAA / TTTAGGG) were used to identify telomeres in the draft assembly. 37 telomeres from 2212 to 18154 bp in length were identified. Further, we used the 7-mer motif to search the contigs produced by NextDenovo, Canu and Hifiasm (using hifi reads only), and identified the three missing ones. Two (CM010418.C1 and CM010423.C1) were supplied by Canu and one (CM010417.C1) was supplied by Hifiasm. Moreover, it was found that the CM010410.C1 upstream telomere produced by Canu (8449 bp) was obviously longer than that of Hifiasm (3045 bp), so that we updated it. The precise locations of telomeres within the ZH13-T2T genome were ascertained by using seqtk (https://github.com/lh3/seqtk). The command used was ‘seqtk telo -s 1 -m CCCTAAA ref.fa’.

### Genome-wide comparisons and identification of SVs

We conducted a comparative analysis using publicly available ZH13 soybean genome data and the assembled T2T sequencing results. First, we aligned the ZH13-T2T genome data to the soybean reference genome using Minimap2 (Version 2.26-r1175) (https://github.com/lh3/minimap2) ^34, 52^. Minimap2 was utilized to map the long sequencing fragments, present in fastq format files of each sample, to the reference genome provided in fasta format ^34^. To enhance comparative efficiency, we utilized the “-ax asm5 --eqx” parameters for fragment alignment, set the software to work with a maximum of 96 threads using the “-t” parameter, filtered out regions with sequence differences greater than 5%, and stored the results of sequence matches or mismatches in SAM format. All other comparison parameters during the process were left at their default settings. We subjected the comparison results of the two genome versions to structural variation detection using the SyRI mutation detection tool (https://github.com/schneebergerlab/syri), configured with default parameters, and saved the mutation detection results as a “syri.out” file ^39^. Subsequently, we employed the Plotsr software to visualize the mutation detection results, using default parameters for the transformation process, and saved the generated images in PDF format ^53^. The visualization command used was “plotsr --sr syri.out --genomes genome.txt”.

### Identification of rDNA and non-coding RNA

5S rRNA is transcribed from the 5S DNA, while 48S rRNA is composed of 28S rRNA, 5.8S rRNA, and 18S rRNA. We identified tRNAs using tRNAscan-SE v2.0, rRNAs using Barrnap v0.9 (https://github.com/tseemann/barrnap), and miRNA and snRNA using INFERNAL (http://eddylab.org/infernal/) against the Rfam (release 12.0) database ^54, 55, 56^. Copy numbers of both 5S rDNA and 48S rDNA were determined using Barrnap. Complete copies of 5S rDNA and 48S rDNA were used as input for genotype identification. Subsequently, a multiple sequence alignment was performed on 5S rDNA and 48S rDNA using MAFFT with default parameters (https://mafft.cbrc.jp/alignment/software/) ^57, 58^. For the 5S rDNA and 48S rDNA, genotype analysis was conducted using single nucleotide polymorphisms (SNPs) and insertions/deletions (indels) with over 10% support from the 5S rDNA copies. It is important to note that all selected indices for 48S rRNAs were located within intergenic spacer regions.

### Repeat identification and gene annotation

We utilized the EDTA pipeline (https://github.com/oushujun/EDTA) for transposable element (TE) annotation ^59, 60^. The main steps involved in identifying repetitive sequences in the genome were as follows: First, we created an indexed database using the RMBlast engine. Then, we employed RepeatModeler (https://www.repeatmasker.org/RepeatModeler/) for de novo prediction, which involved five iterative rounds to obtain repetitive sequences and Stockholm format seed alignment files ^61^. Subsequently, we performed genome annotation using RepeatMasker ^62, 63, 64^. Low-complexity sequences and small RNA (pseudo) genes were not masked, and the search for insertions of missing sequences was disabled. Repeat-masked genome and repeat sequence library constructed by RepeatModeler and RepeatMasker were used for subsequent TE analysis.

For ab initio annotation, we utilized BUSCO (https://github.com/metashot/busco) to create a training dataset for Augustus (https://github.com/Gaius-Augustus/Augustus)^65^. Based on this training dataset, we further applied Augustus to predict the coding regions of genes on the masked genome. We further analyzed the component composition of the previous gap region and enriched the new annotated genes by GO and KEGG using KOBAS, with p<0.05 as the threshold ^66^.

### Centromere localization

We employed Tandem Repeat Finder (TRF, version 4.09.1) (https://github.com/Benson-Genomics-Lab/TRF) to discern and classify satellite, small satellite, and microsatellite sequences within the soybean T2T genome ^67^. The default parameters utilized for TRF were set to ‘2 7 7 80 10 50 500 -f -d -m’, and the results of TRF annotation were merged using TRF2GFF (https://github.com/Adamtaranto/TRF2GFF). We manually eliminated tandem repeats with fewer than five copies and redundant occurrences. Sequences characterized by lengths of less than 10 base pairs (bp), between 10 bp and 100 bp, and exceeding 100 bp were respectively categorized as microsatellites, minisatellites, and satellites. Building upon the results obtained from the previous EDTA pipeline and TEsorter (https://github.com/zhangrengang/TEsorter), we obtained TE annotation files and the total number of copies of different period sequences in various chromosomes ^59, 60, 68^. Utilizing two high-copy satellite repeat subfamilies, CentGm-1 and CentGm-2, which are exclusive to the centromeric region, we ascertained the approximate location of the centromere ^42^. Lastly, by employing the Integrative Genomics Viewer (IGV) browser, we observed an overlap between the regions with TE annotation loss and the region where the 91/92bp-long sequences were concentrated, thereby identifying the centromere region ^69, 70, 71, 72^.

## Data availability

The genome assembly data generated in this study can be achieved from NCBI with BioProject ID: PRJNA1015379 and BioSample accession: SAMN37355196.

