## Supplementary Figure 1-14 for "A telomere-to-telomere genome assembly of Zhonghuang 13, a widely-grown soybean variety from the original center of Glycine max"

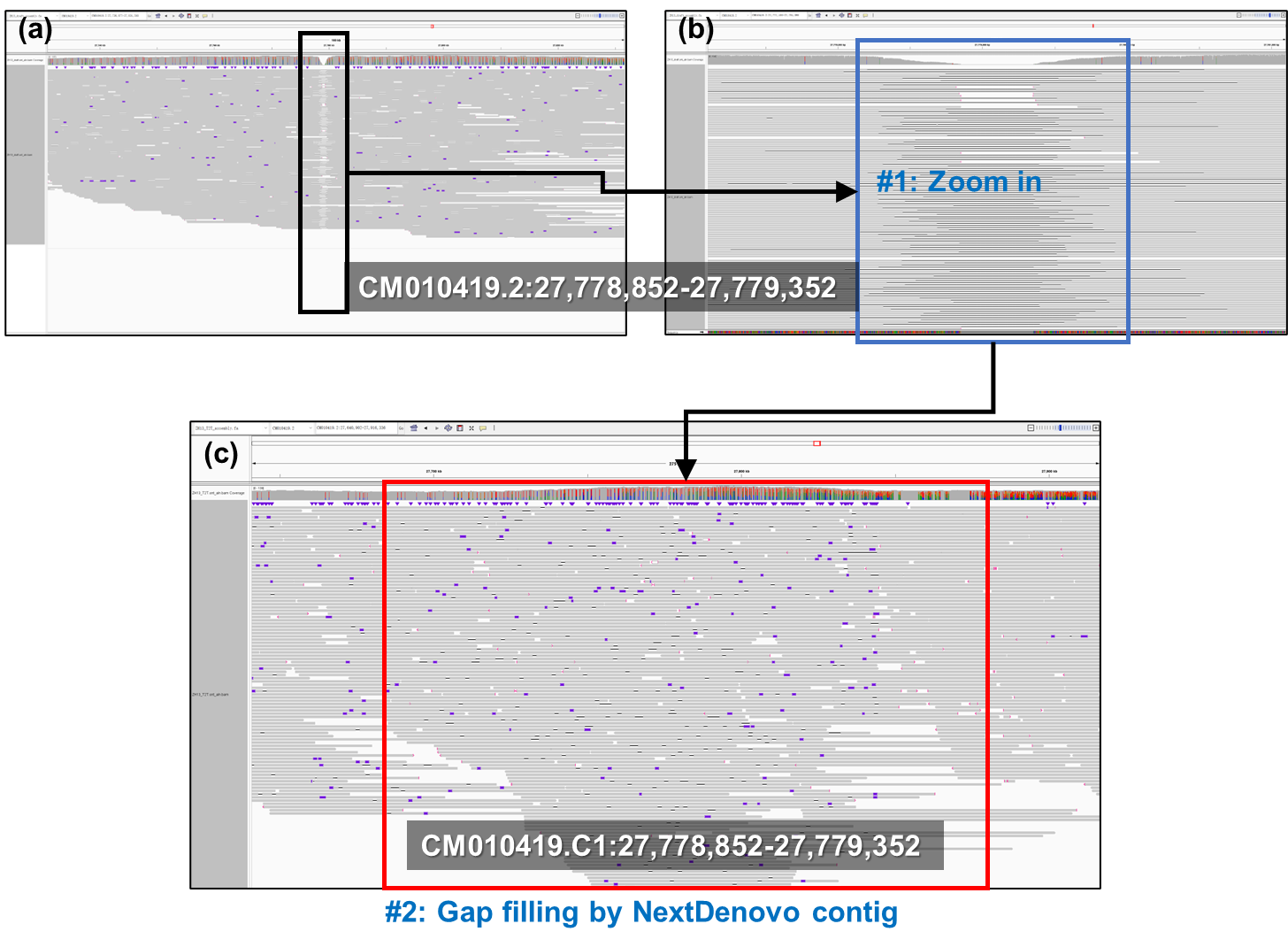

**Supplementary Fig. 1. The filling of gap2.**

a) the IGV snapshot of ONT ultralong read alignment in the surrounding region of gap2 in ZH13-ref-2019; b) a zoom-in view of gap2; c) the IGV snapshot of ONT ultralong read alignment against the gap filled by a NextDenovo contig (consecutive alignments and normal coverage were observed).

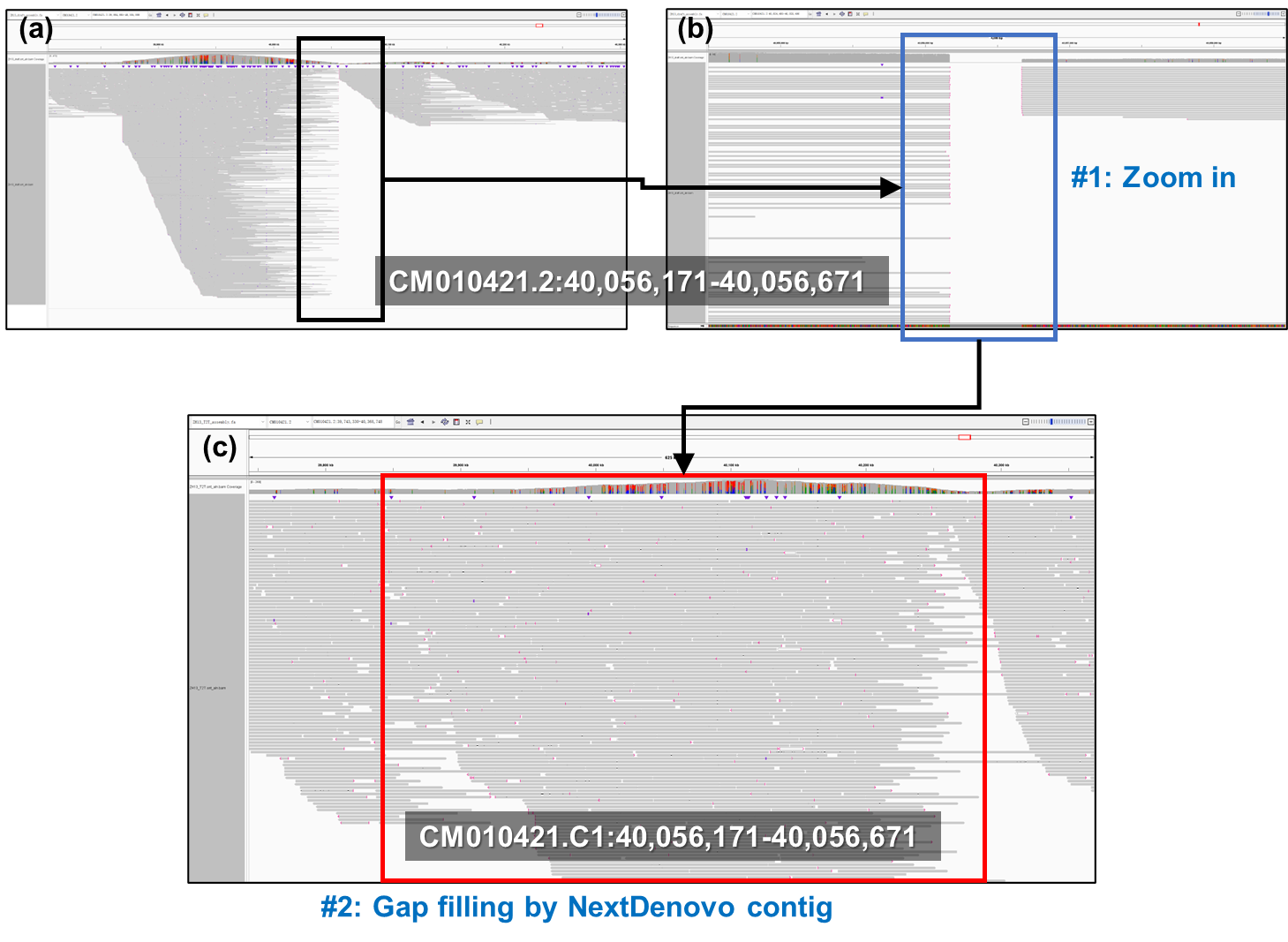

**Supplementary Fig. 2. The filling of gap4**

a) the IGV snapshot of ONT ultralong read alignment in the surrounding region of gap4 in ZH13-ref-2019; b) a zoom-in view of gap4; c) the IGV snapshot of ONT ultralong read alignment against the gap filled by a NextDenovo contig (consecutive alignments and normal coverage were observed).

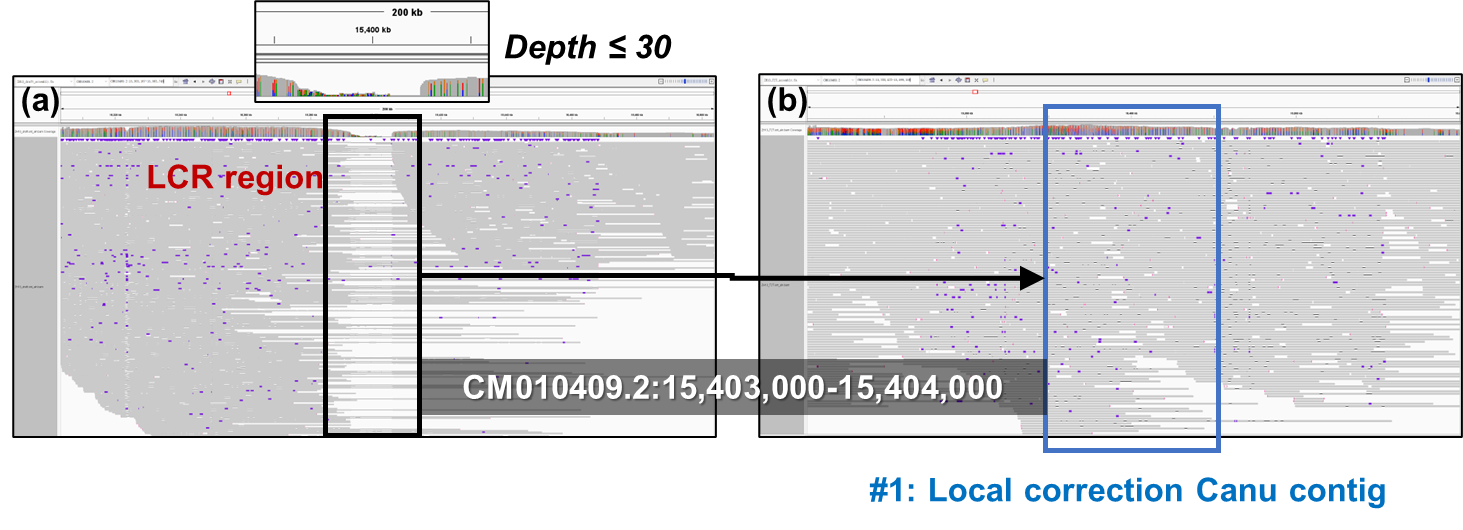

**Supplementary Fig. 3. The correction of LCR1.**

a) the IGV snapshot of ONT ultralong read alignment around LCR1; b) the IGV snapshot of ONT ultralong read alignment after Canu contig correction (consecutive alignments and normal coverage were observed).

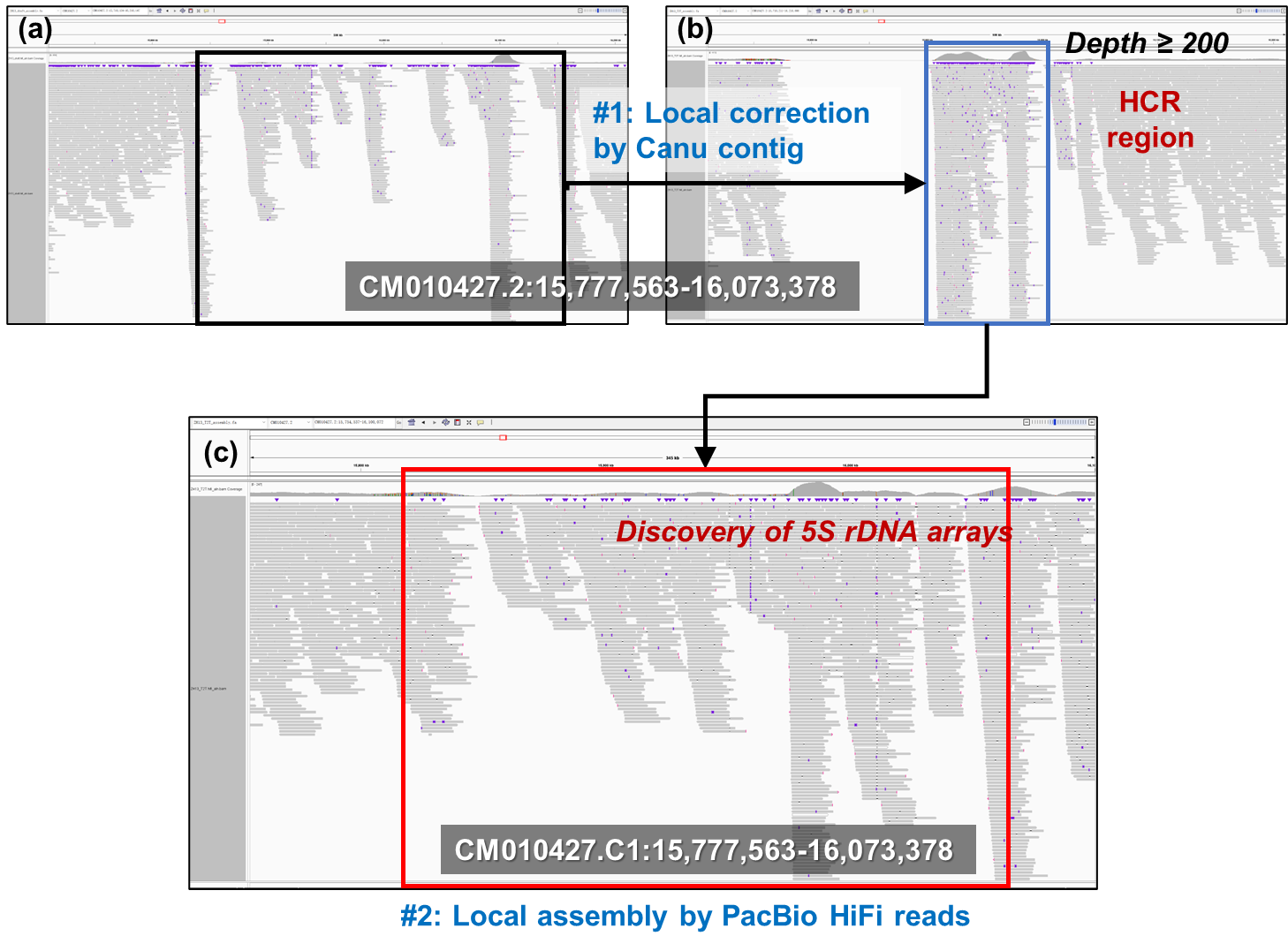

**Supplementary Fig. 4. The correction of LCR2.**

a) the IGV snapshot of PacBio Hifi read alignment around LCR2; b) the IGV snapshot of PacBio Hifi read alignment after Canu contig correction (HCR was observed and 5S rDNA array was identified); c) the IGV snapshot of PacBio Hifi read alignment after local assembly-based correction with anchored reads, consecutive read alignments were observed. Moreover, by manually checking the alignment details, we also found many reads having very low MAPQ, i.e., each of the reads had multiple candidate mapping positions and cannot be confidently aligned. Overall, normal coverage was proved by considering all the primary and secondary alignments of the reads.

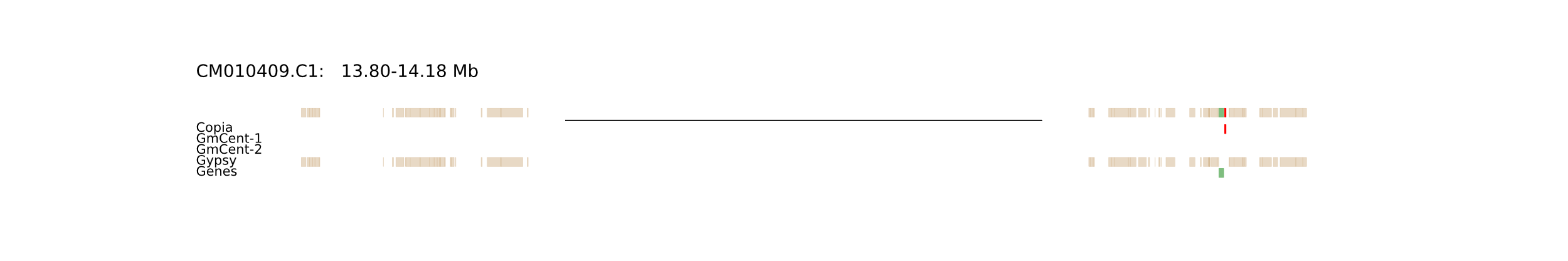

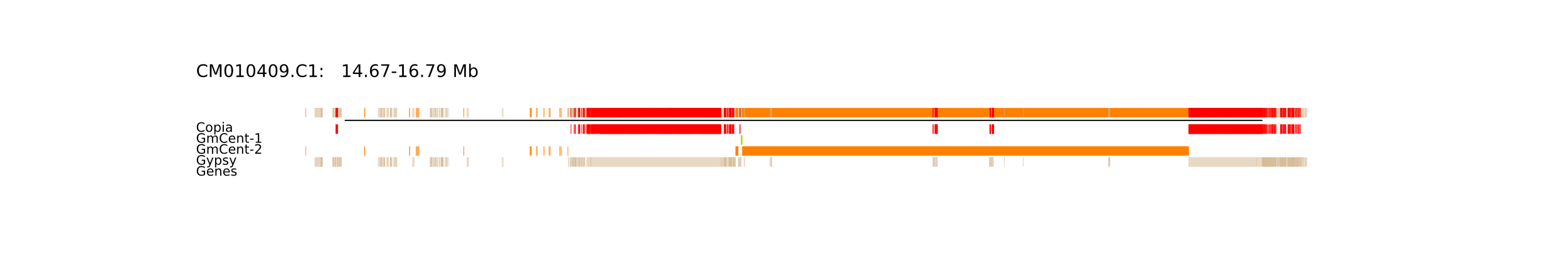

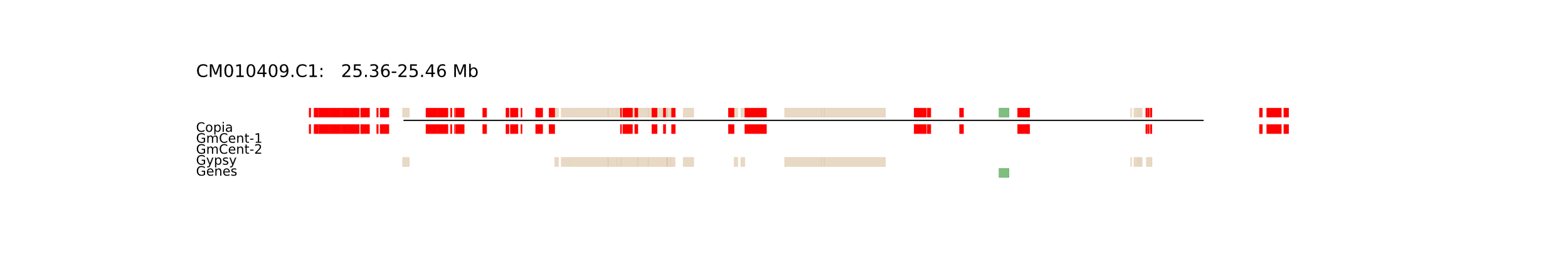

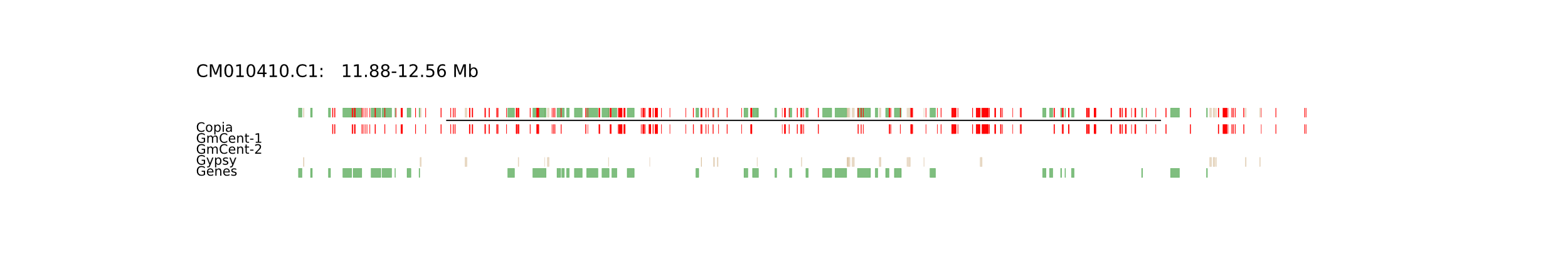

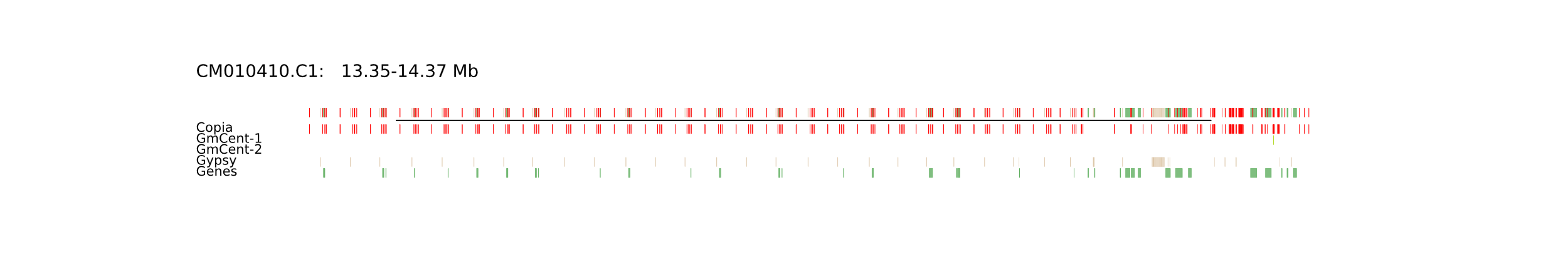

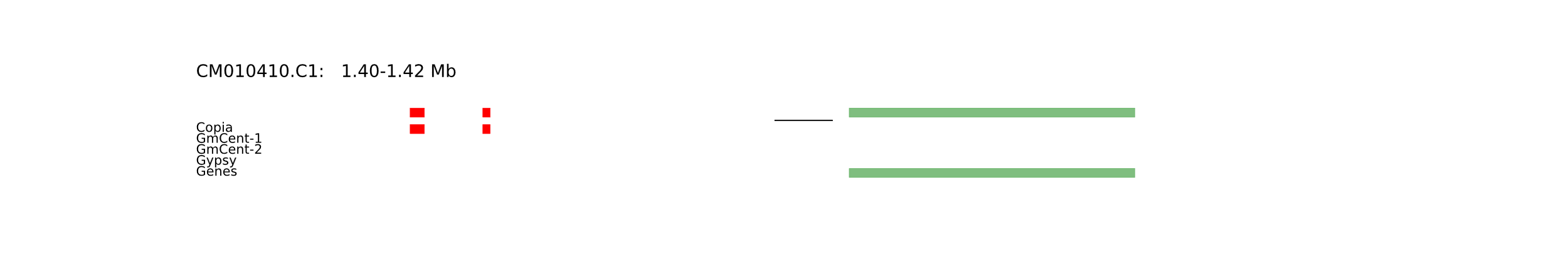

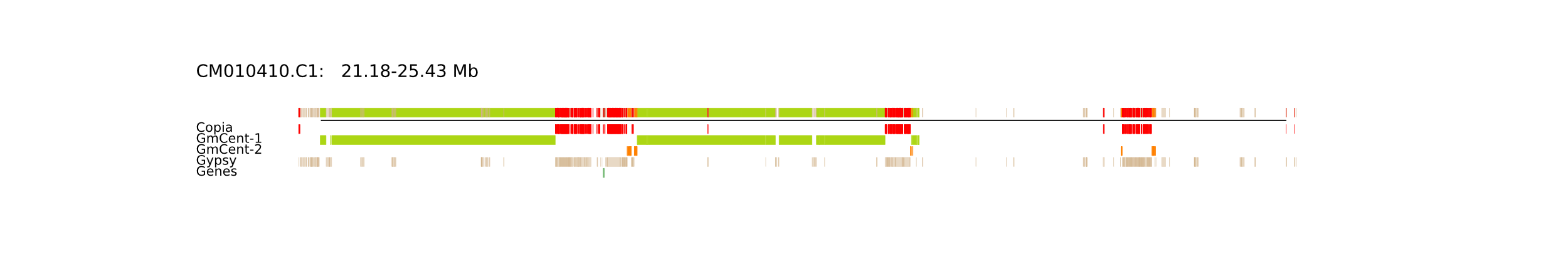

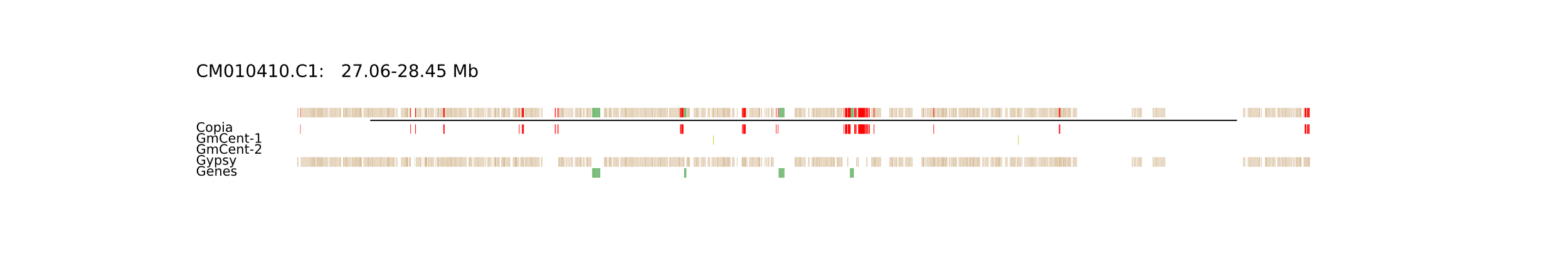

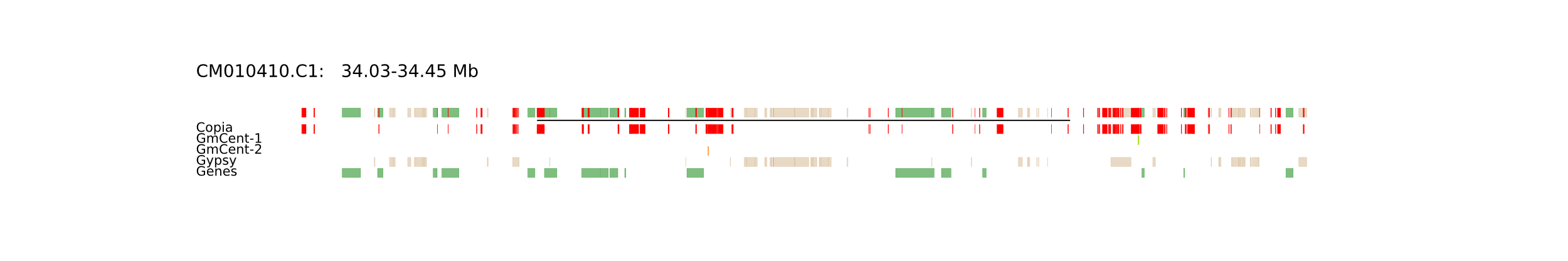

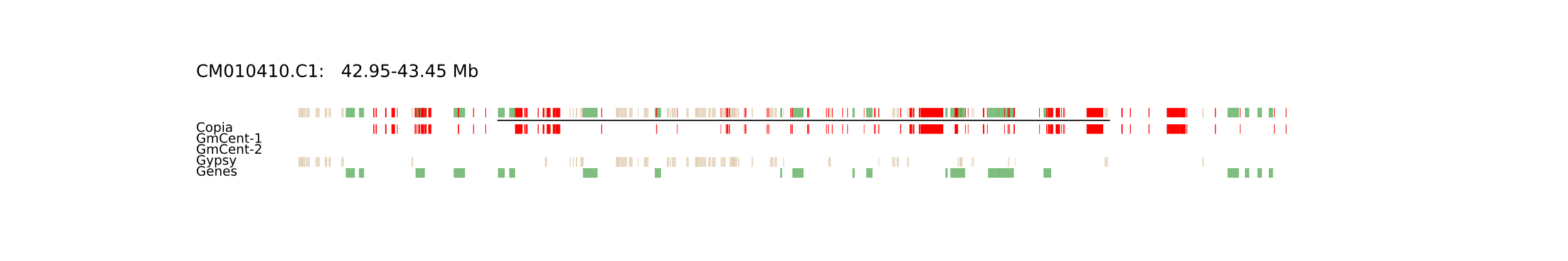

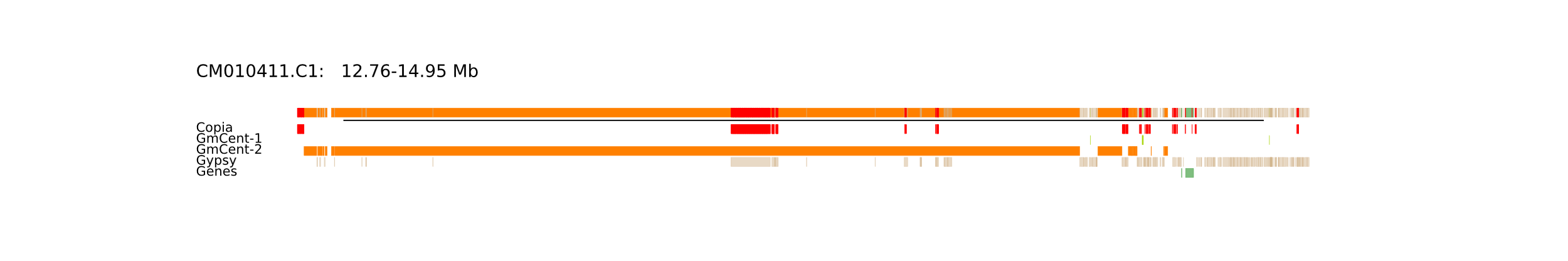

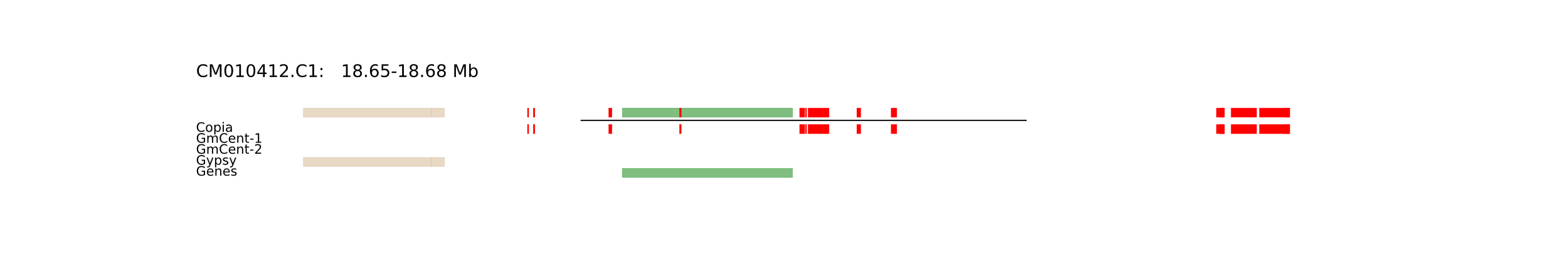

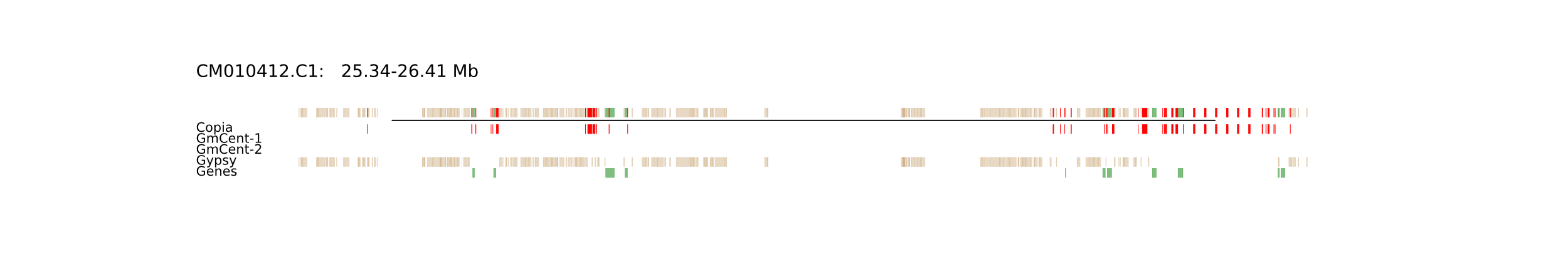

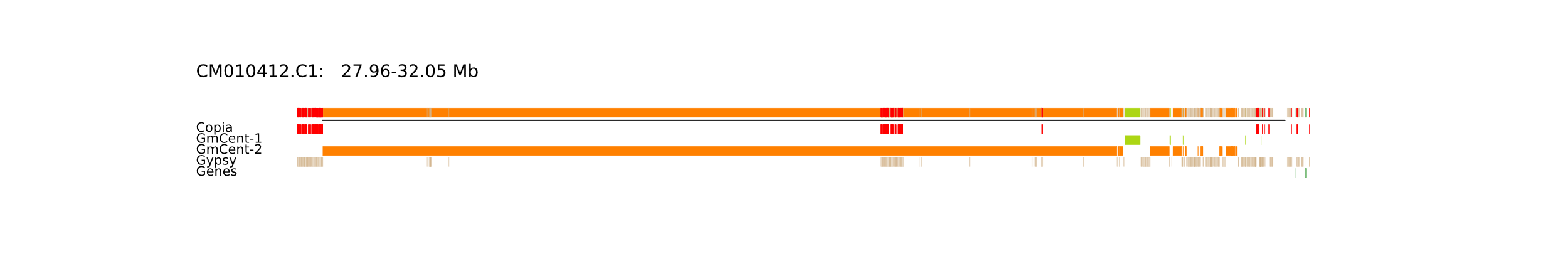

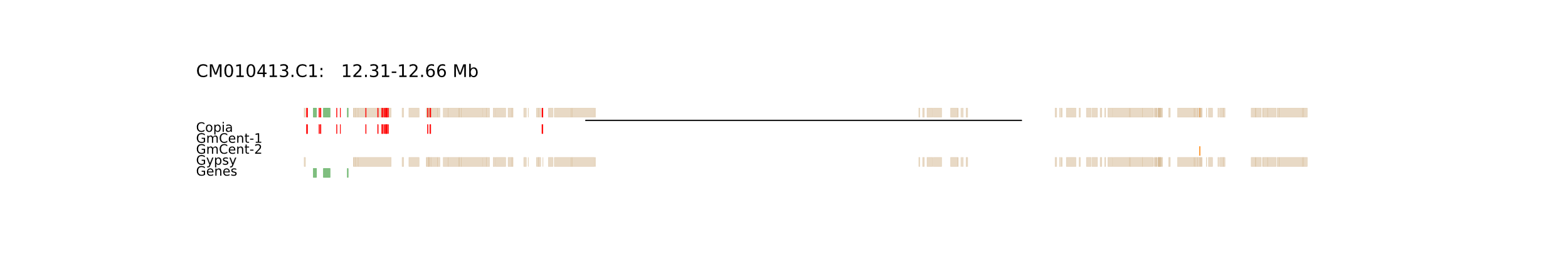

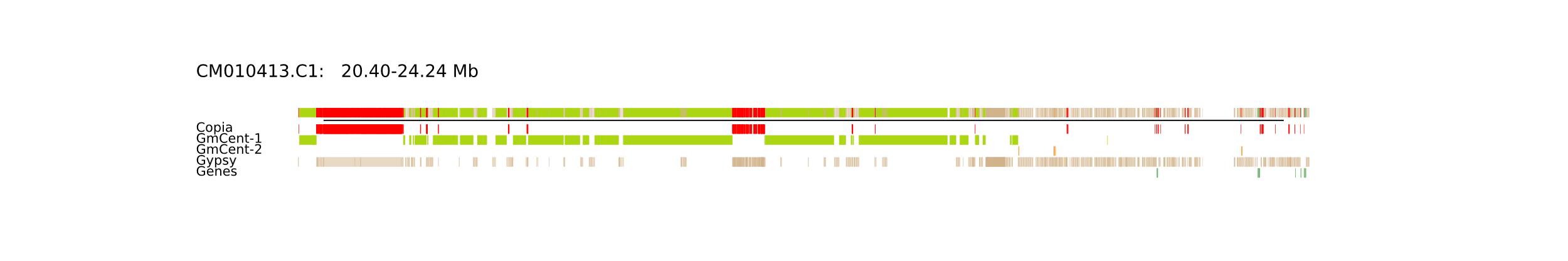

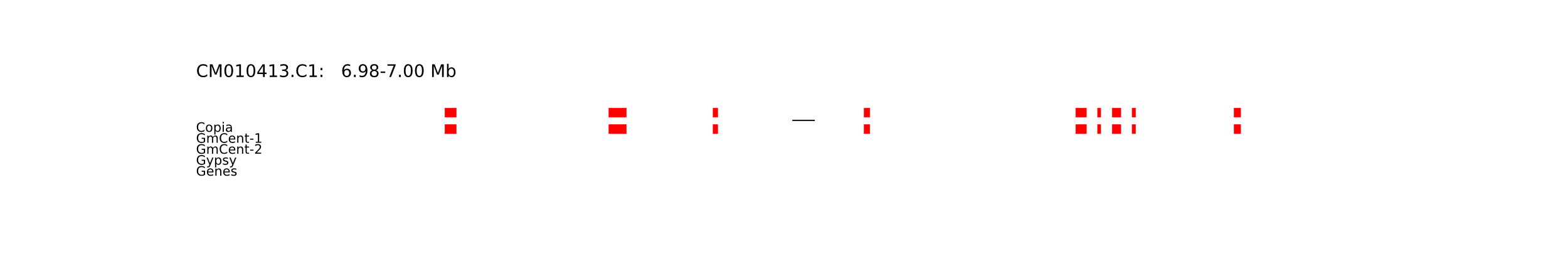

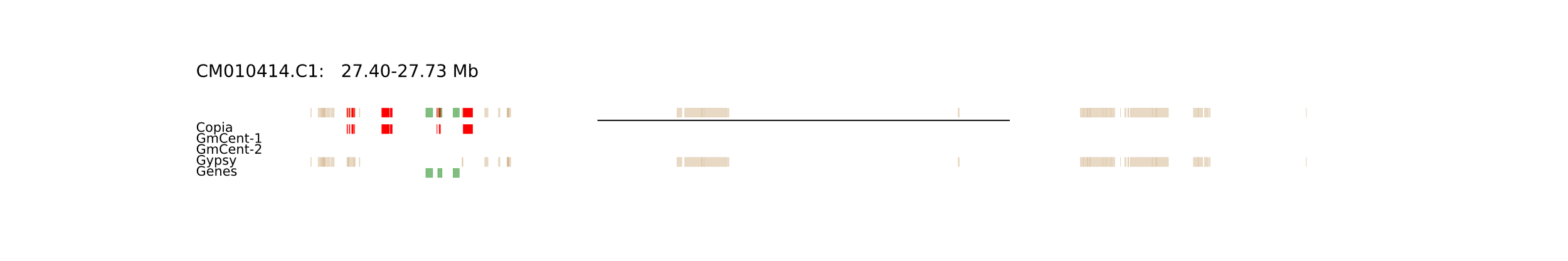

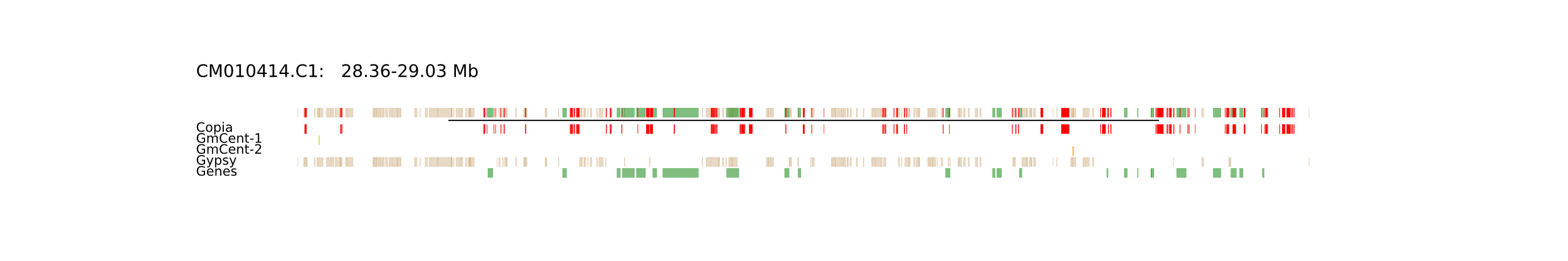

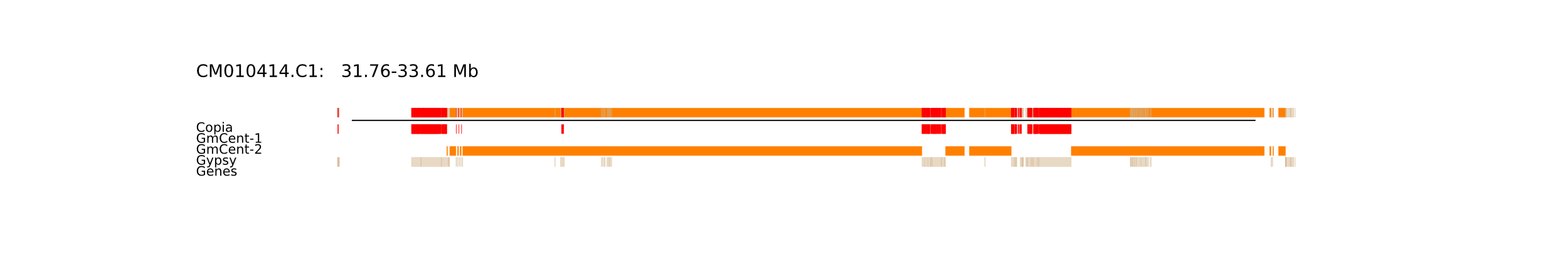

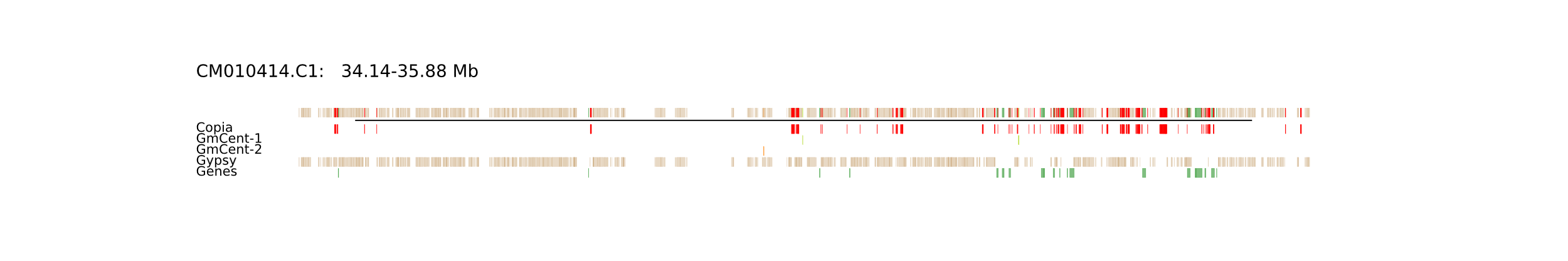

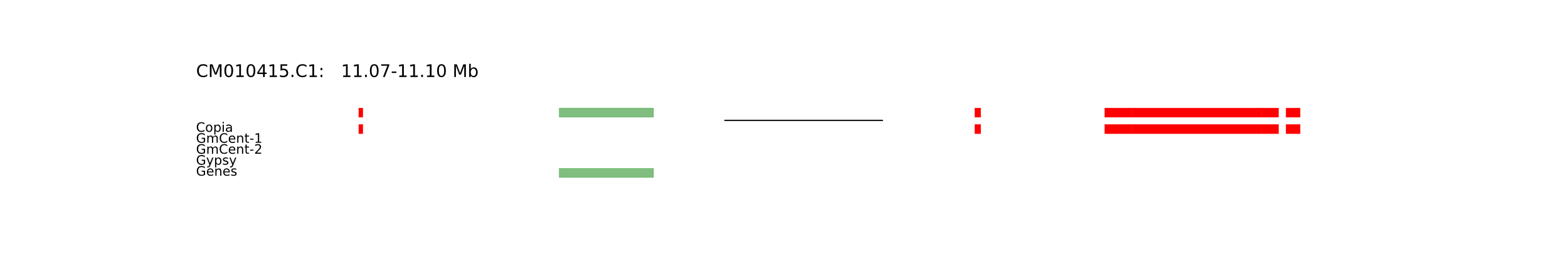

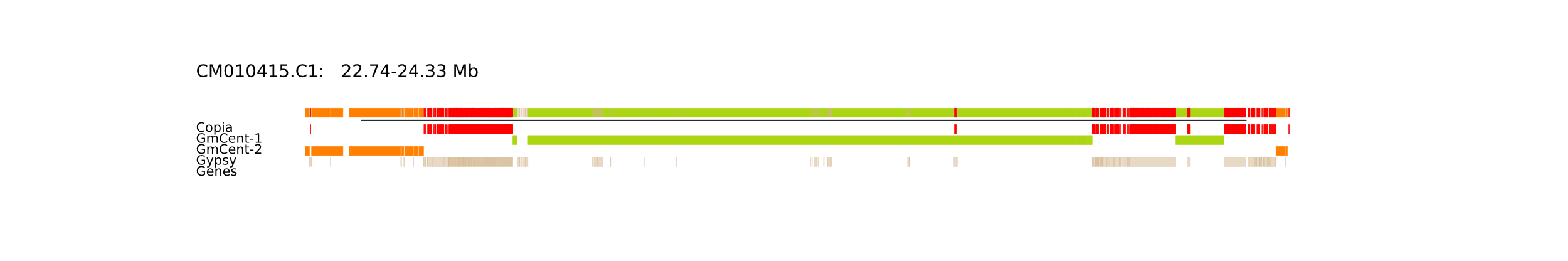

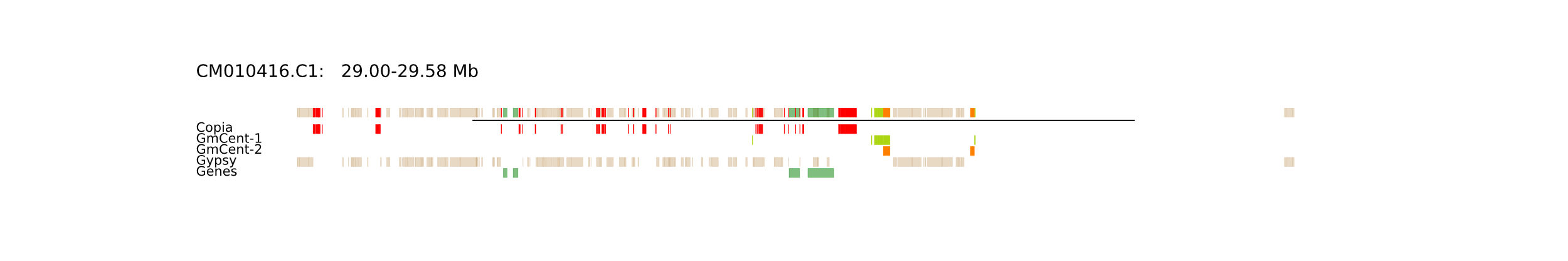

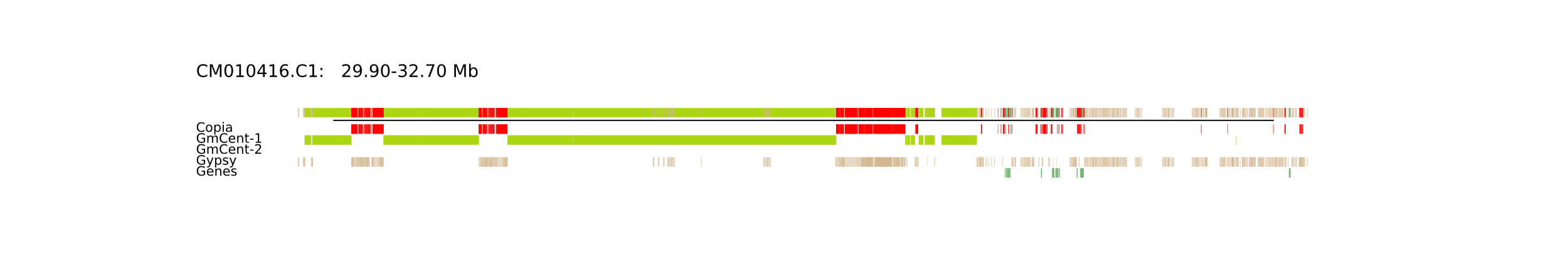

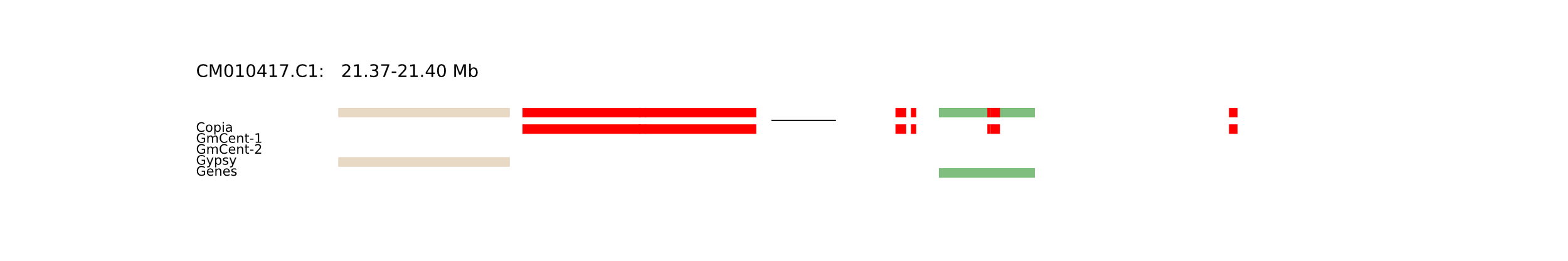

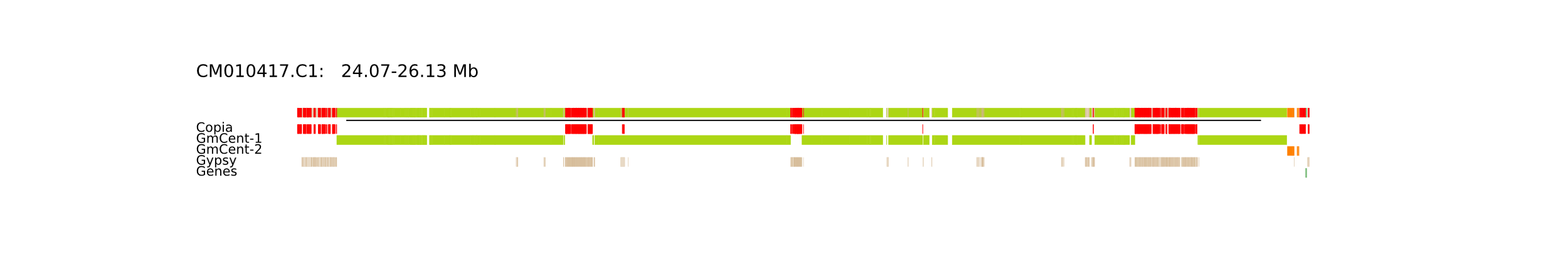

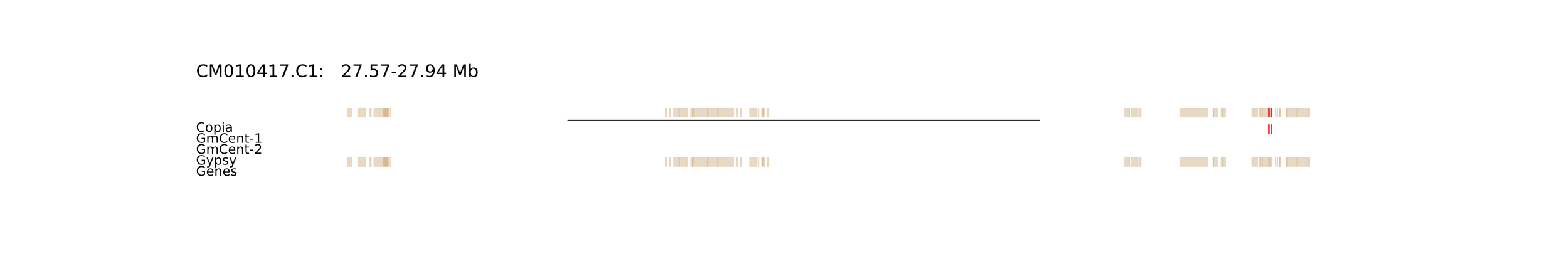

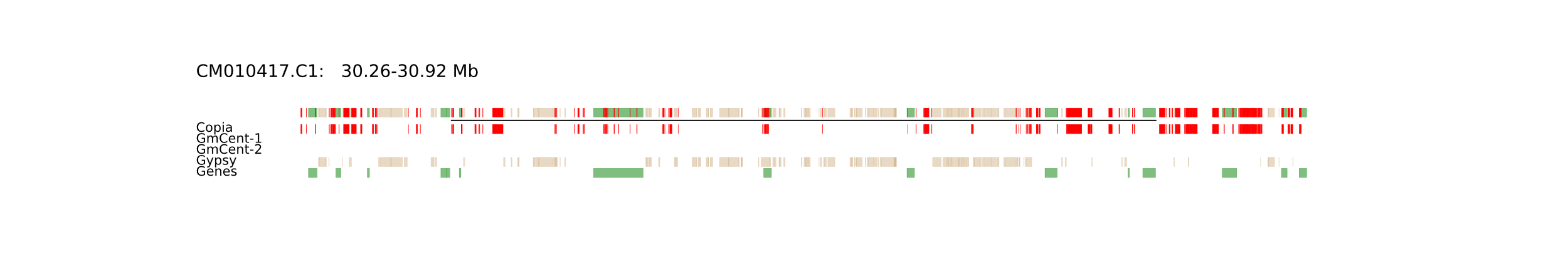

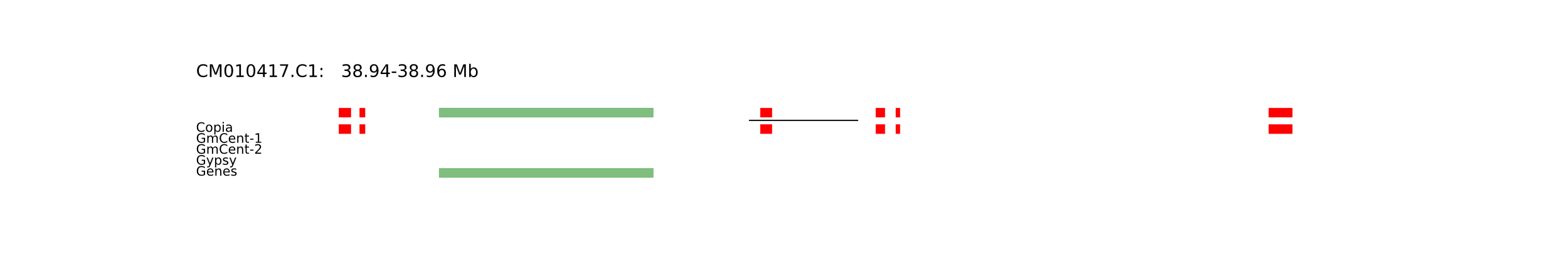

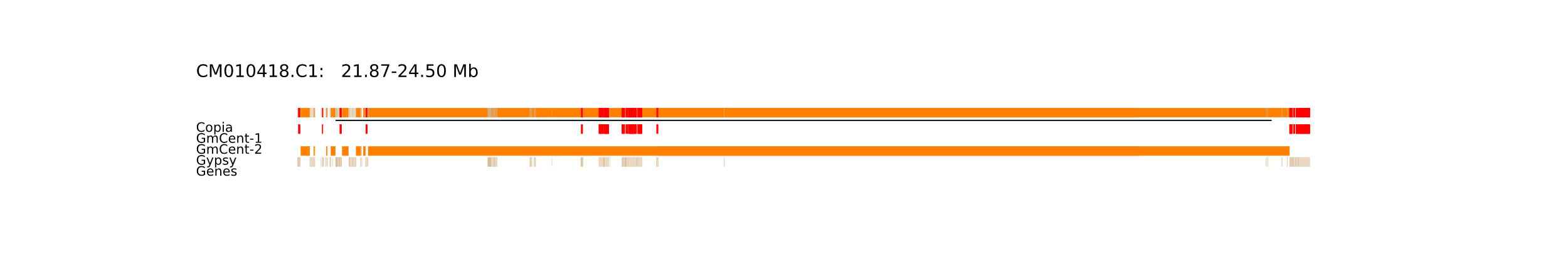

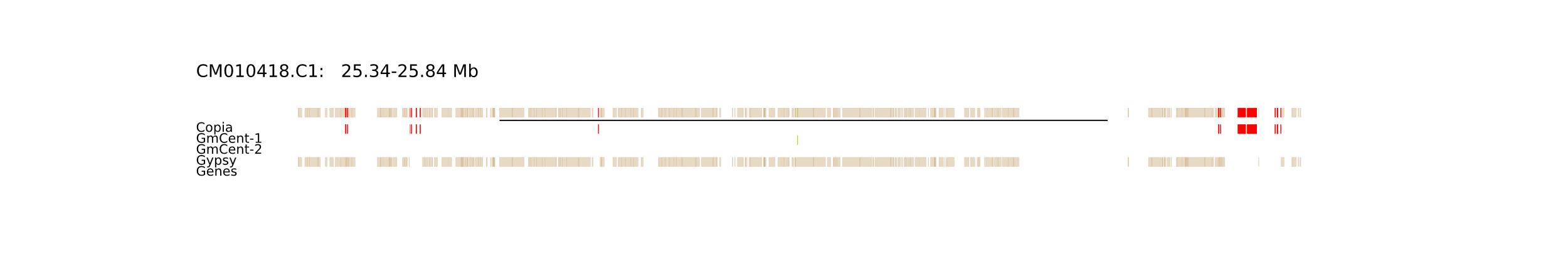

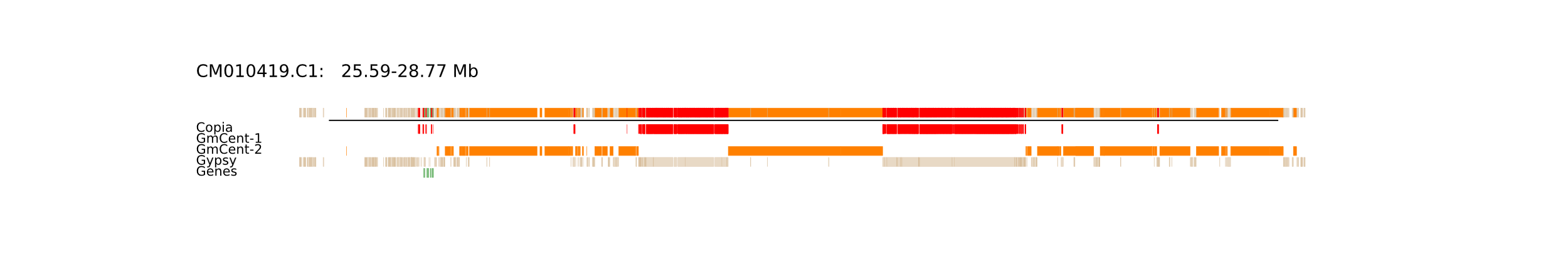

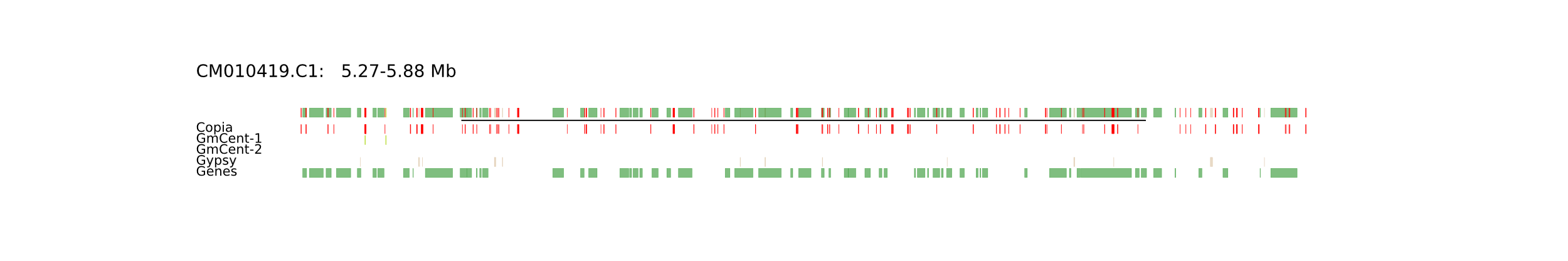

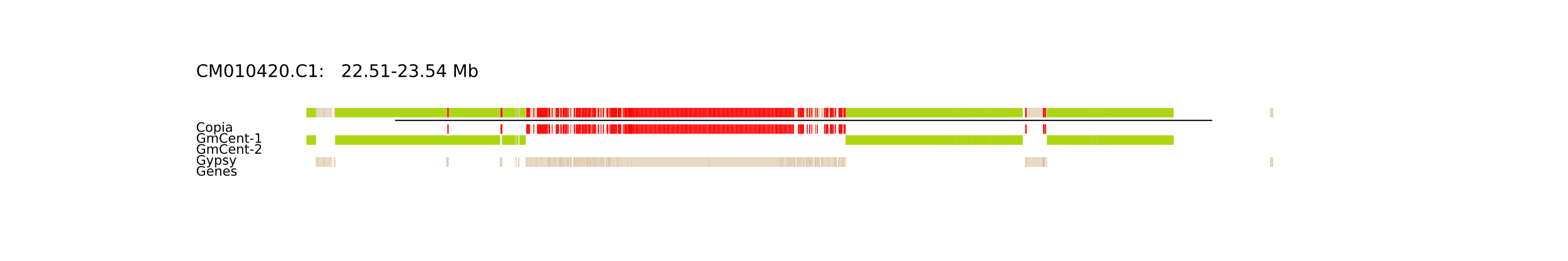

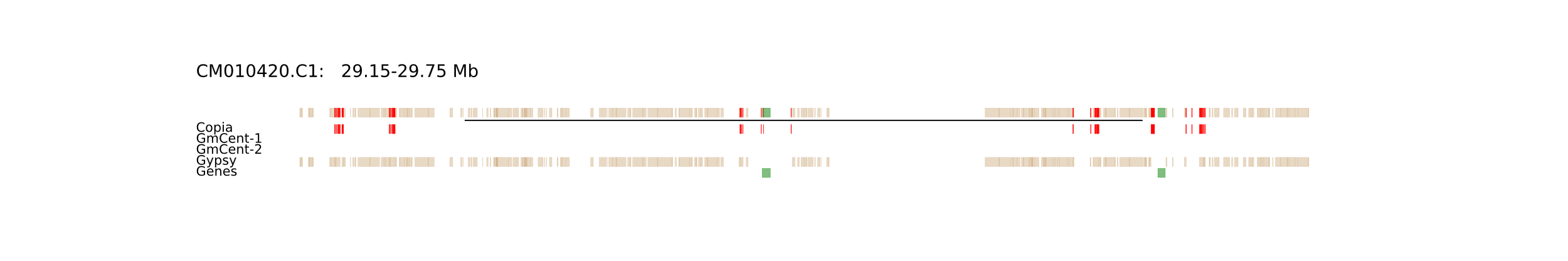

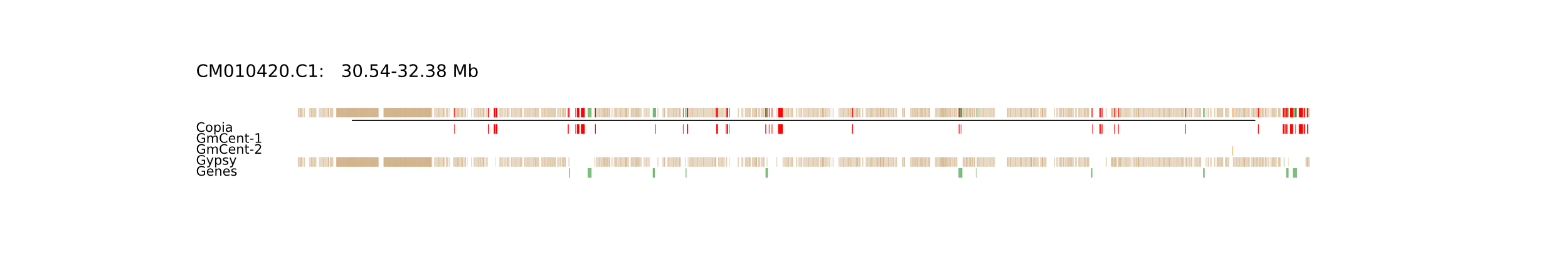

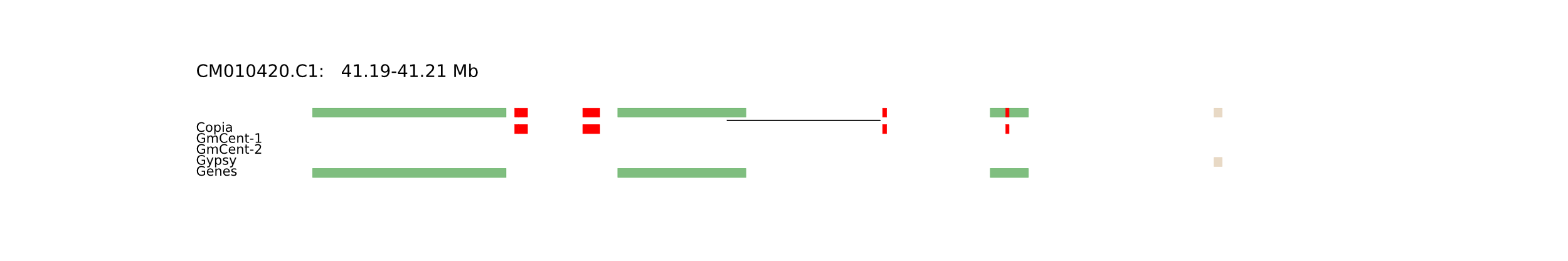

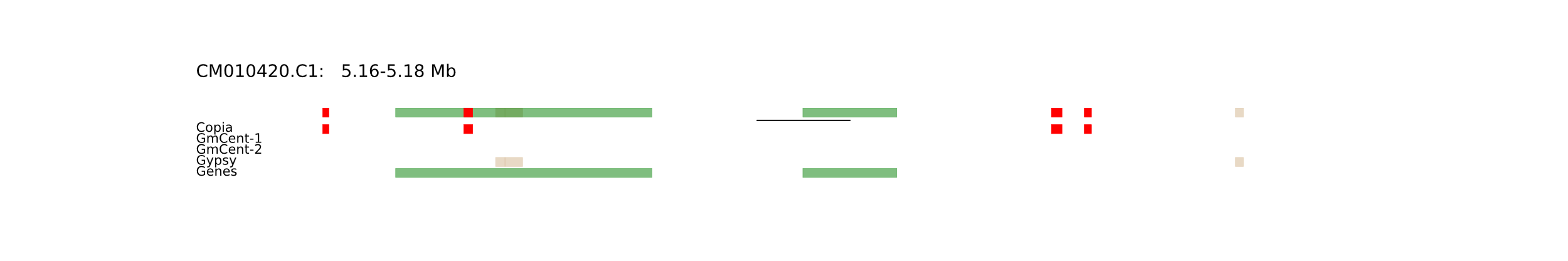

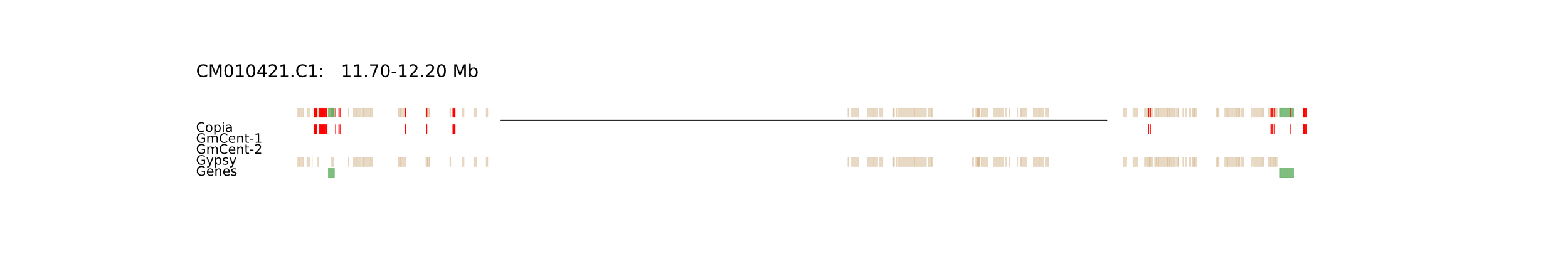

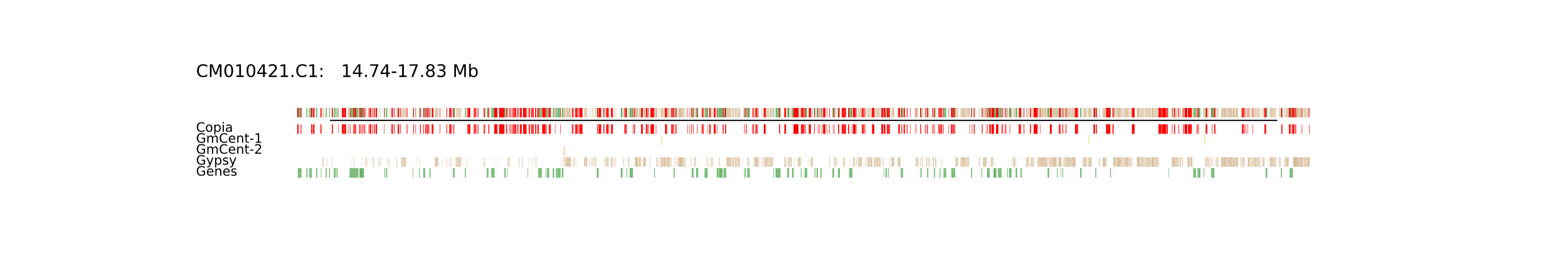

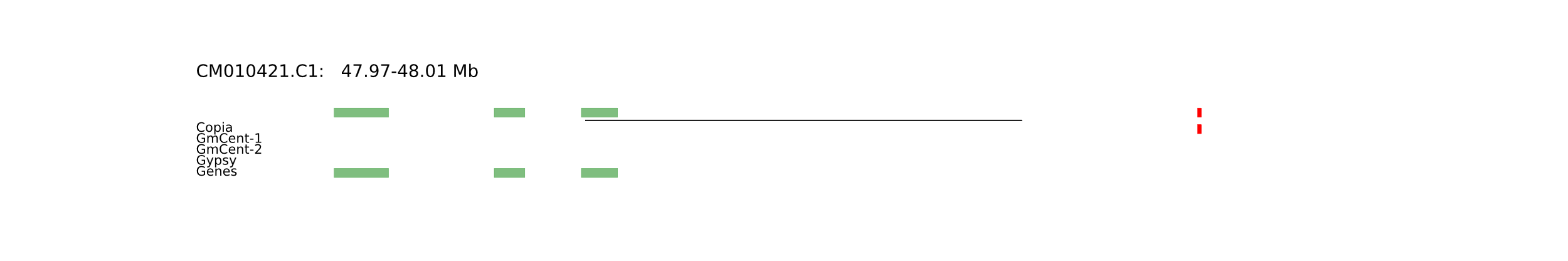

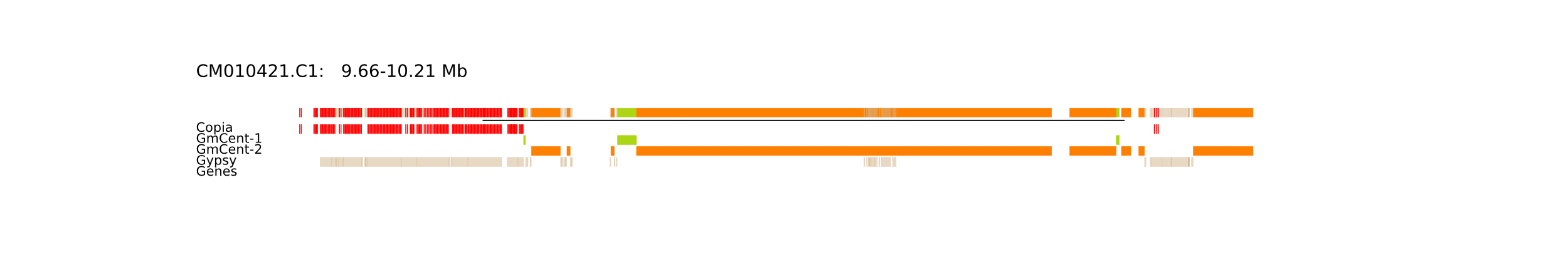

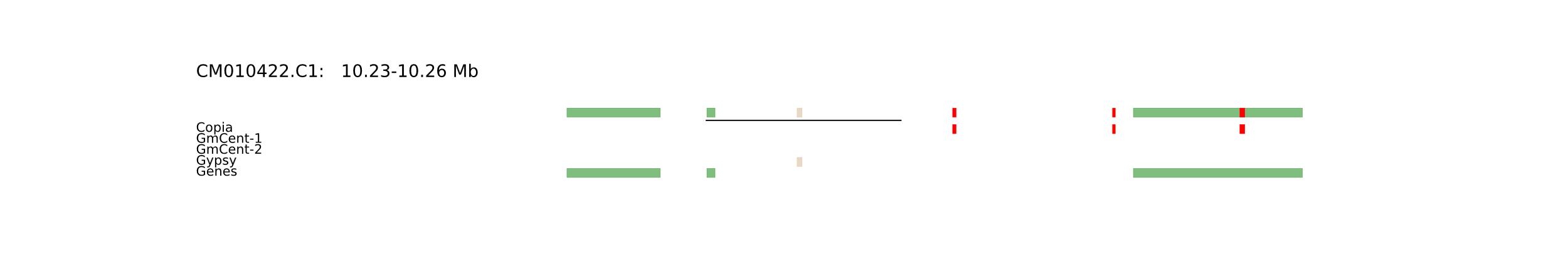

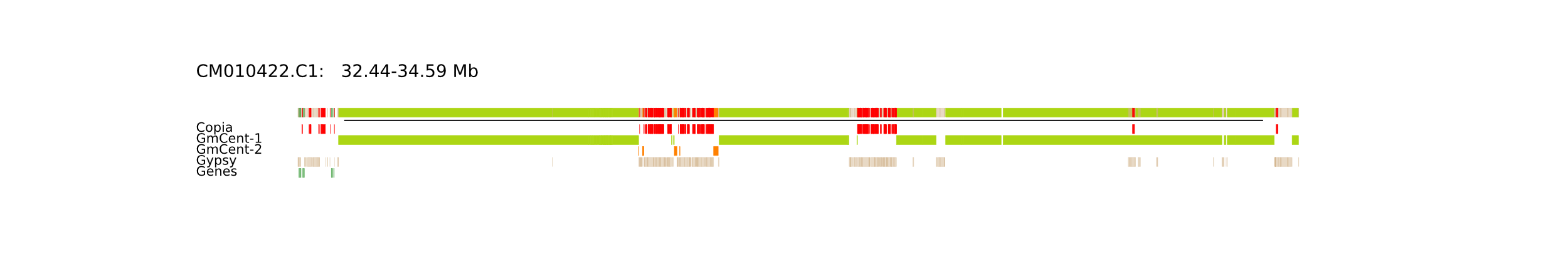

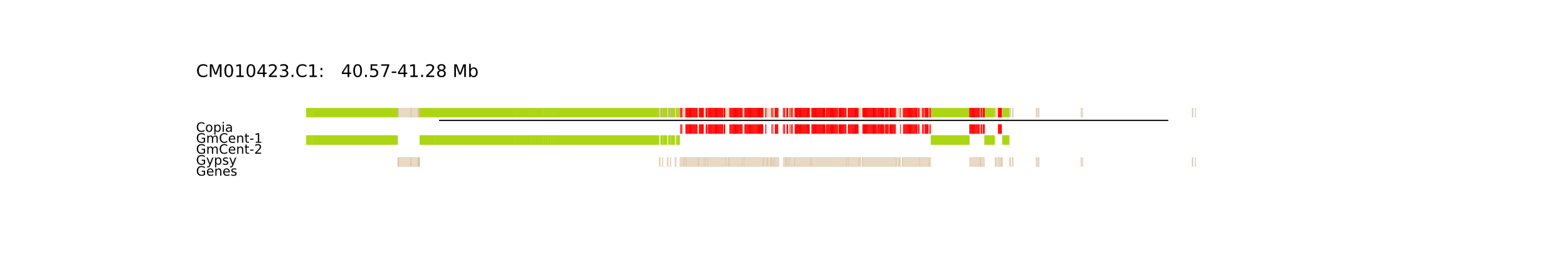

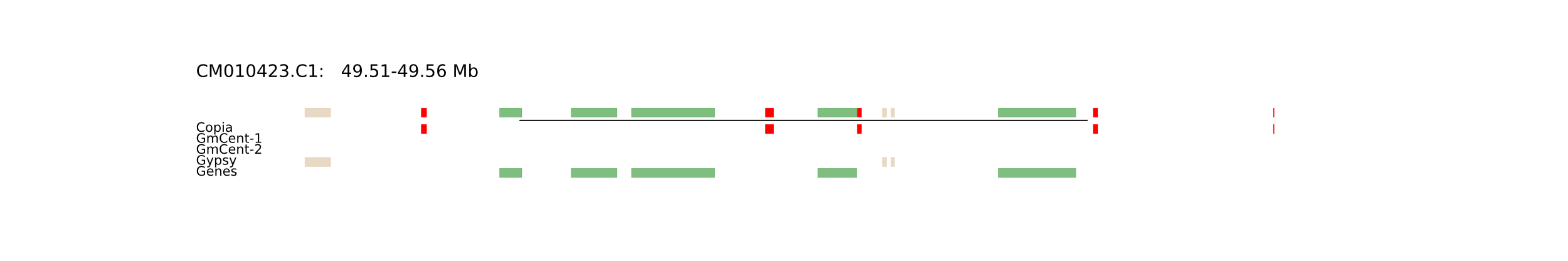

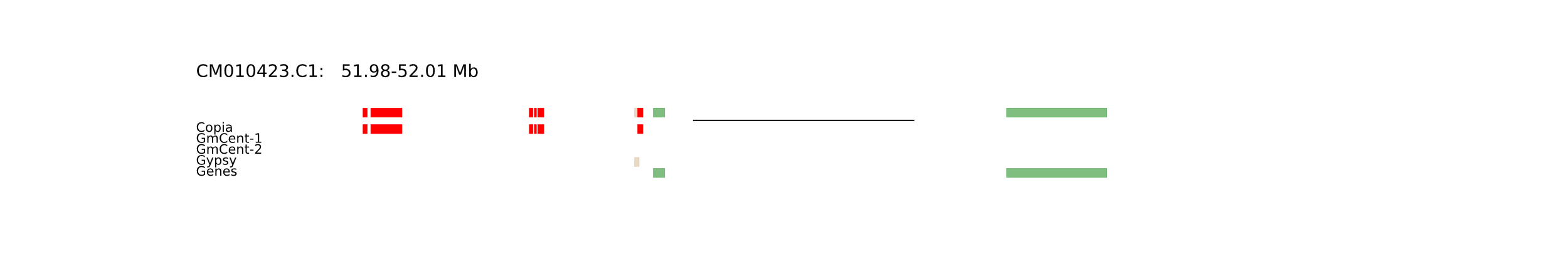

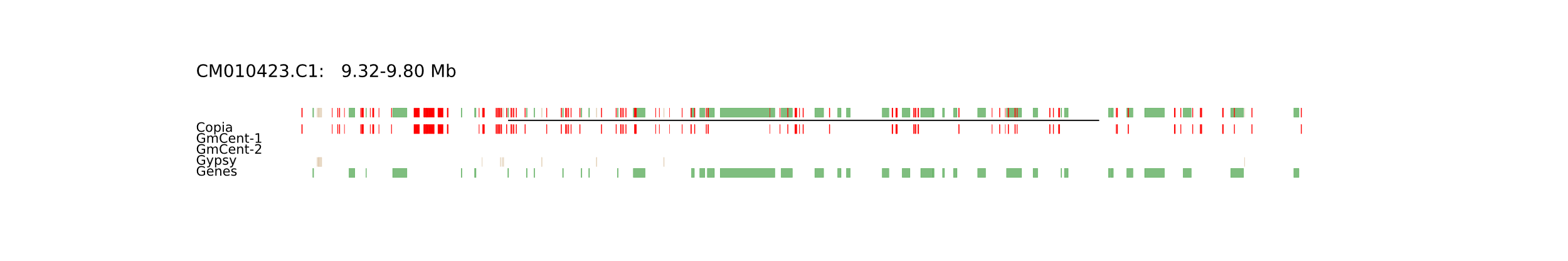

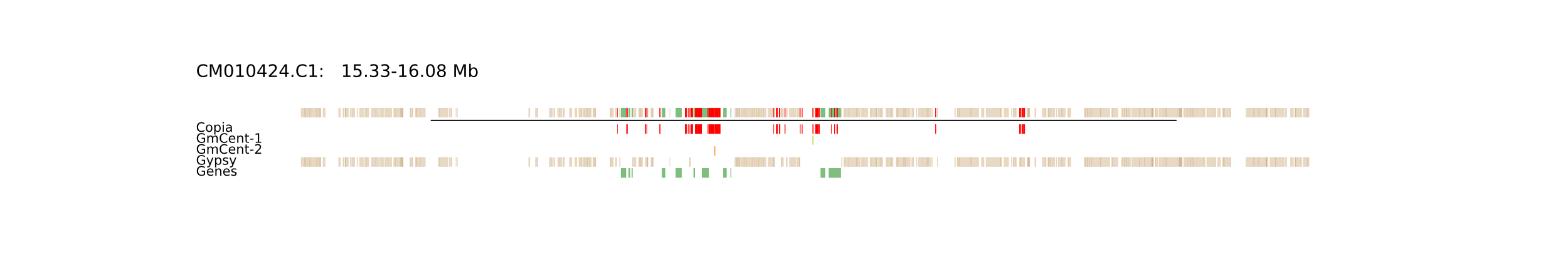

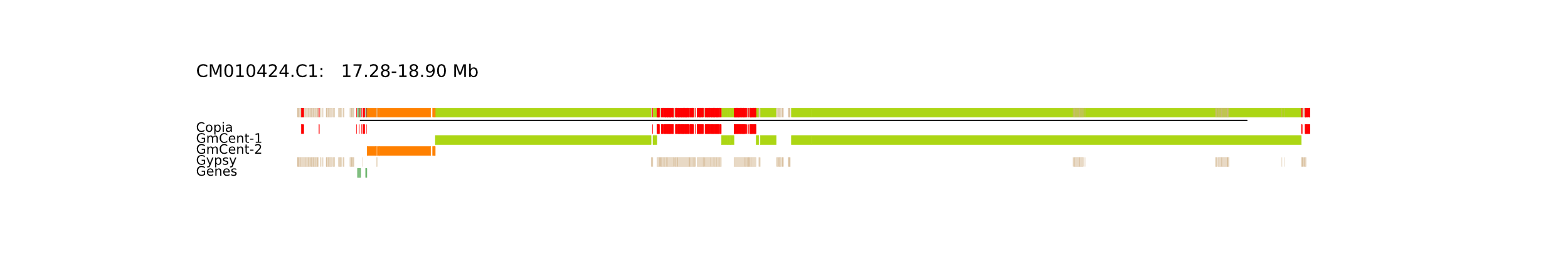

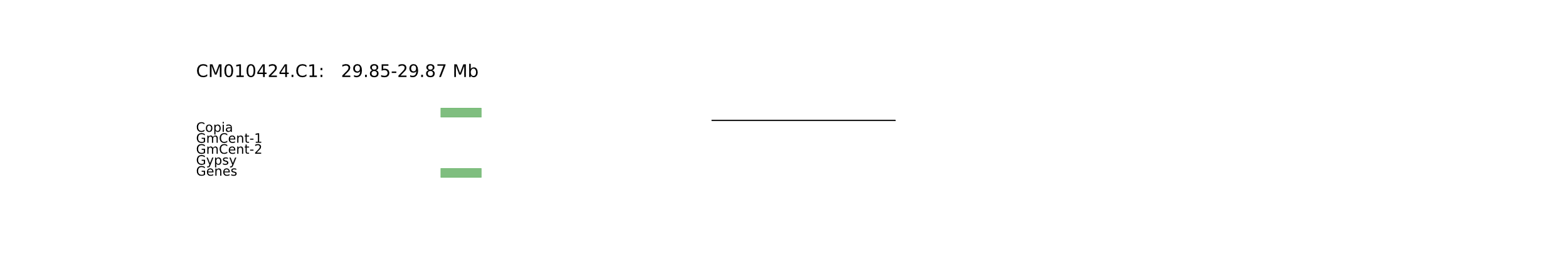

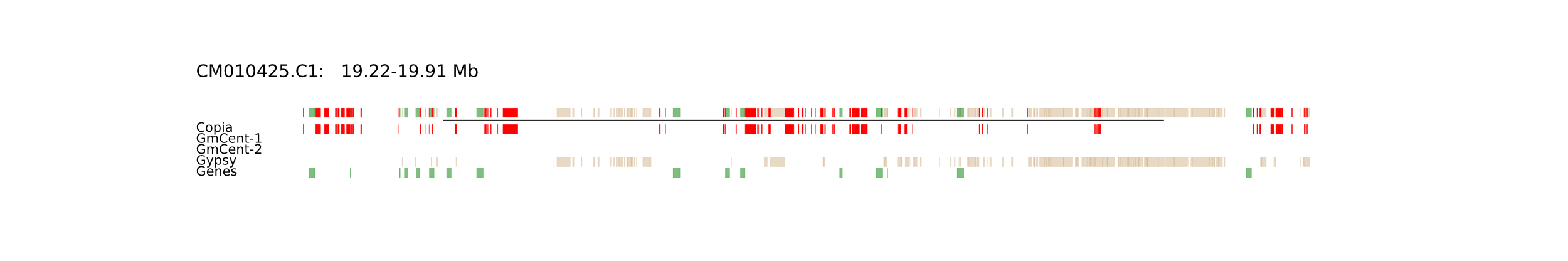

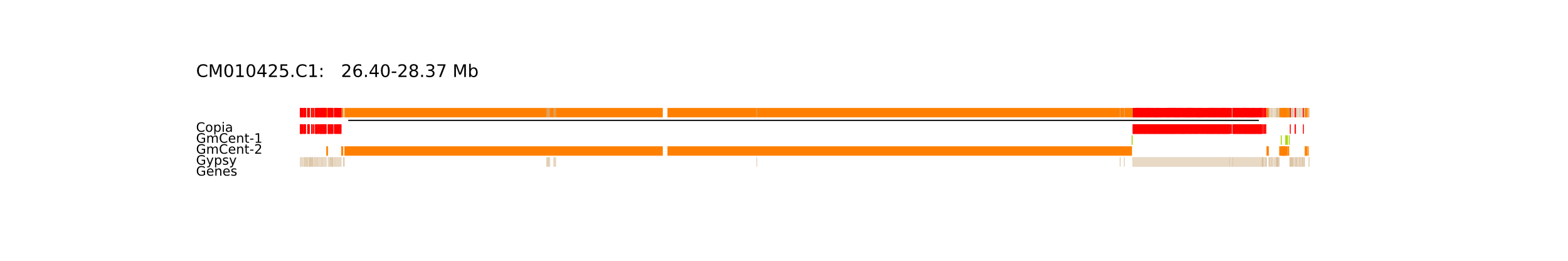

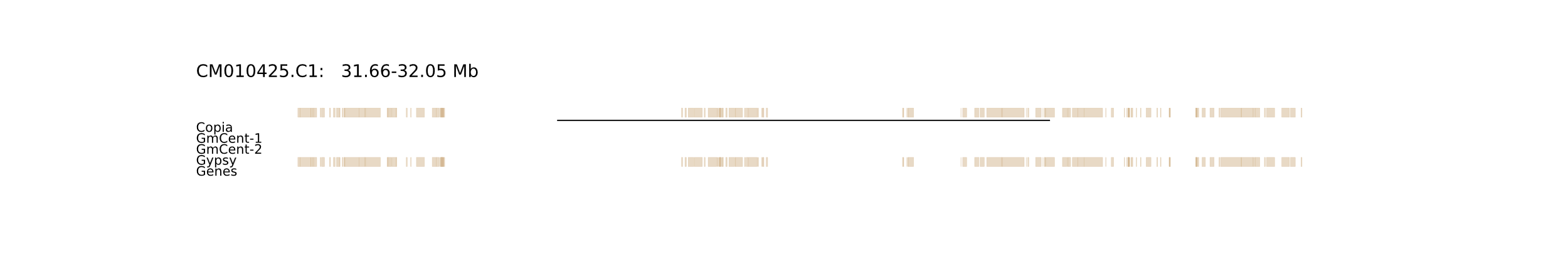

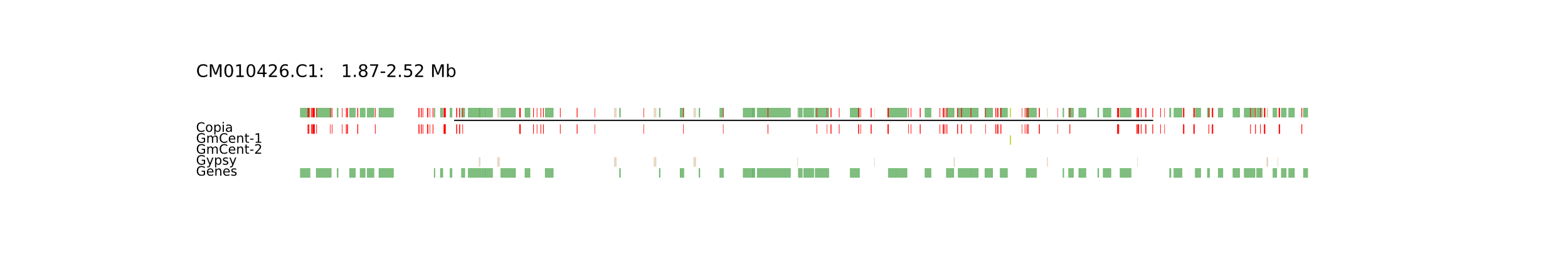

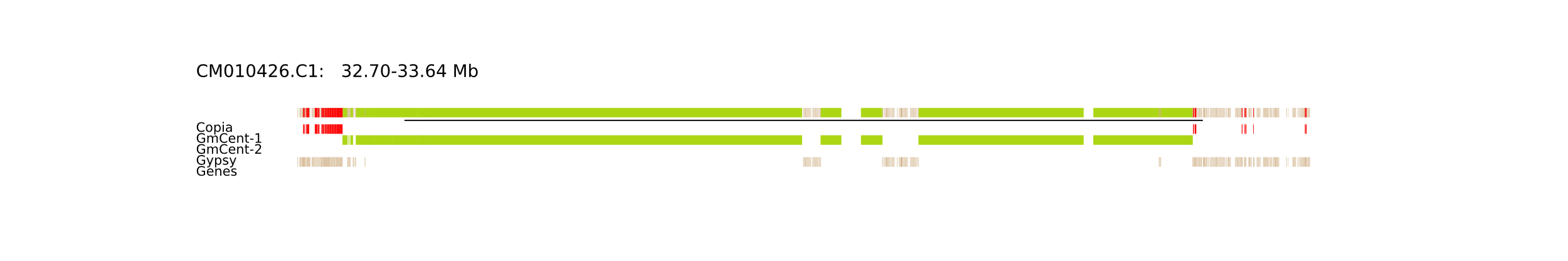

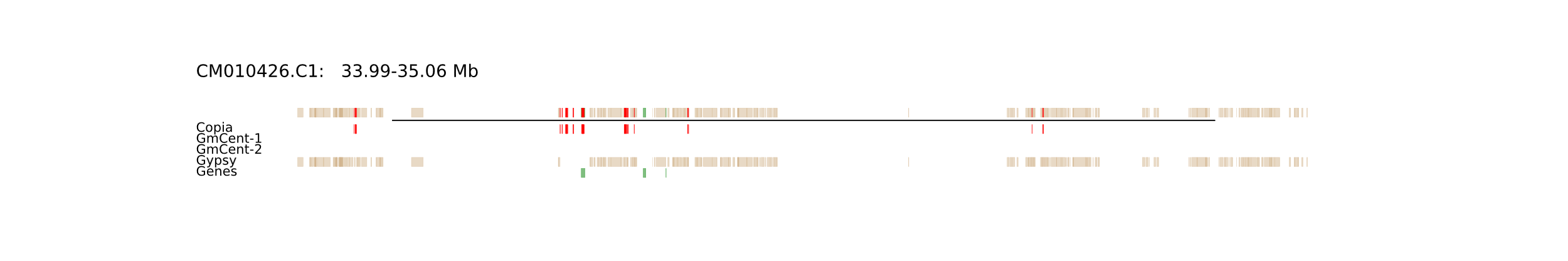

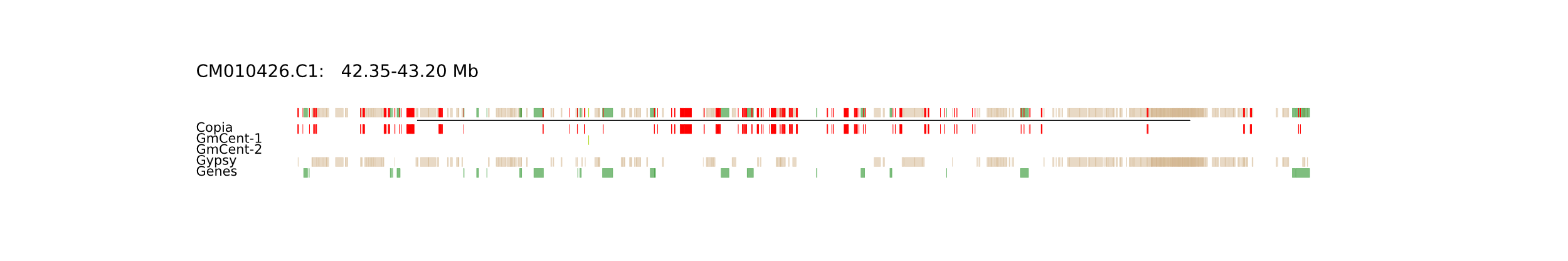

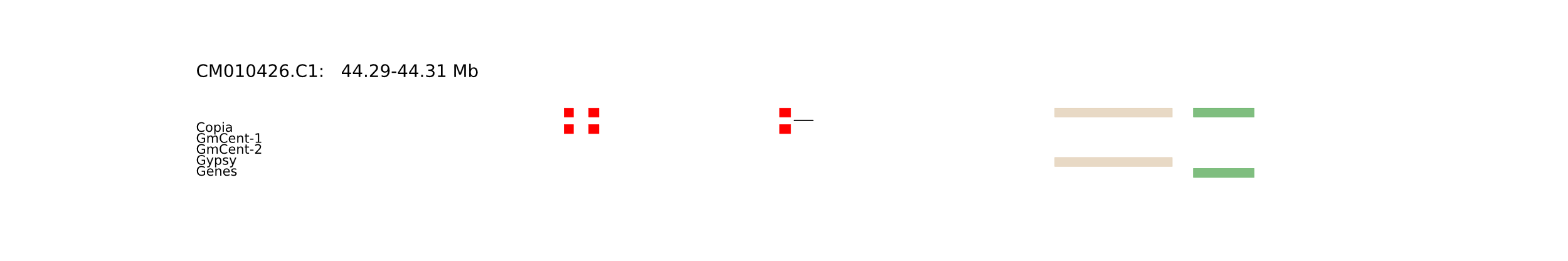

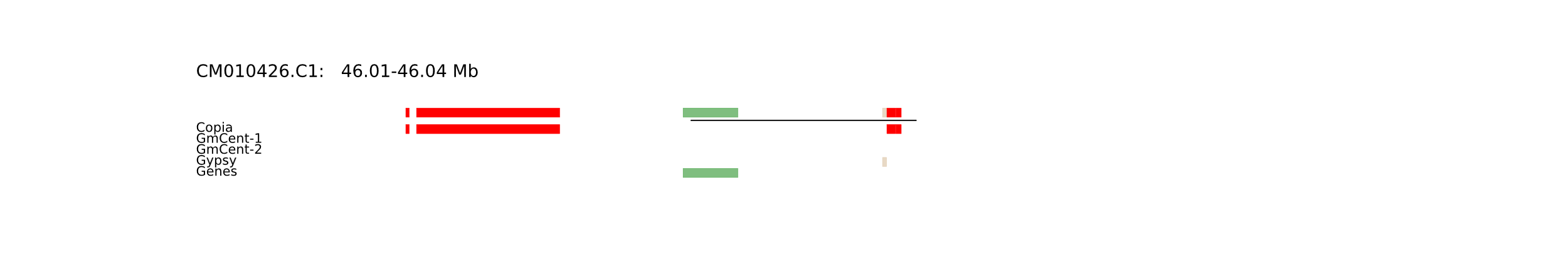

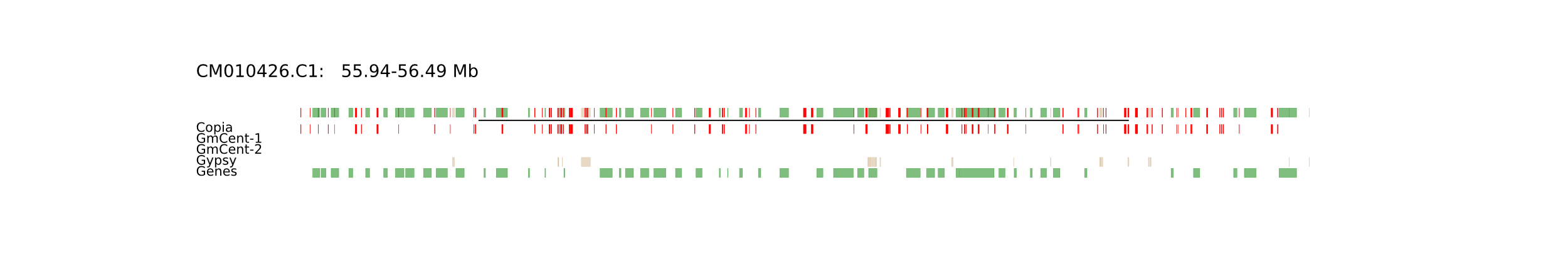

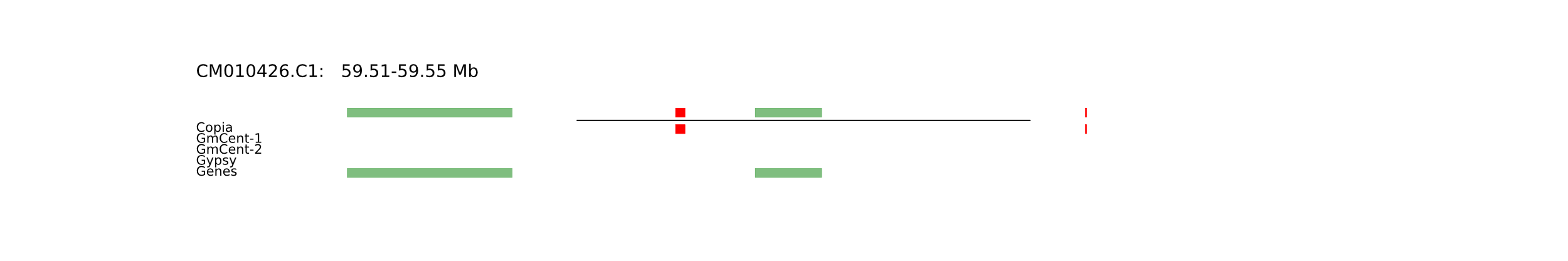

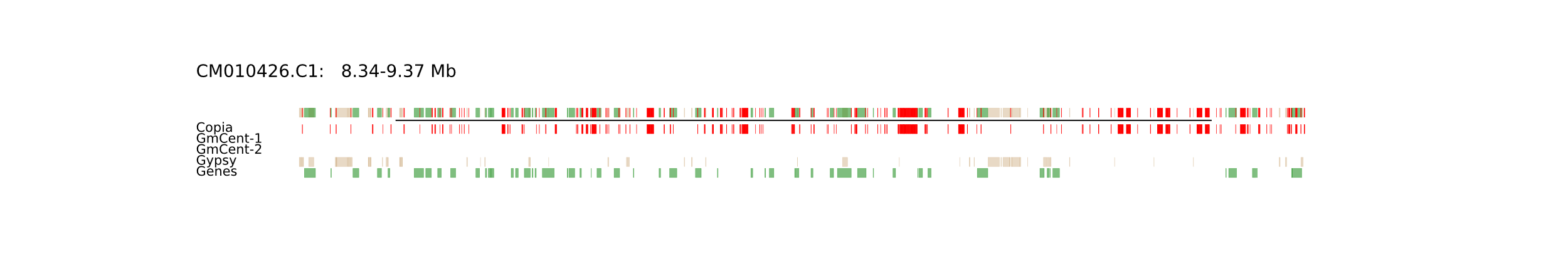

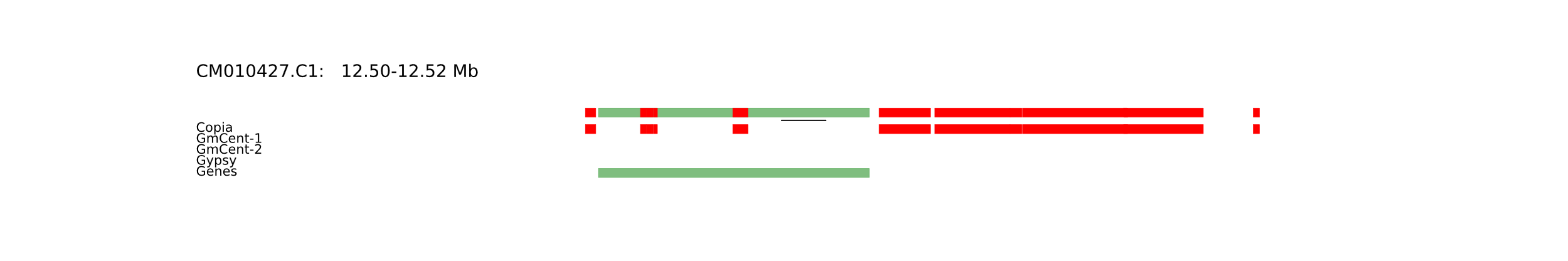

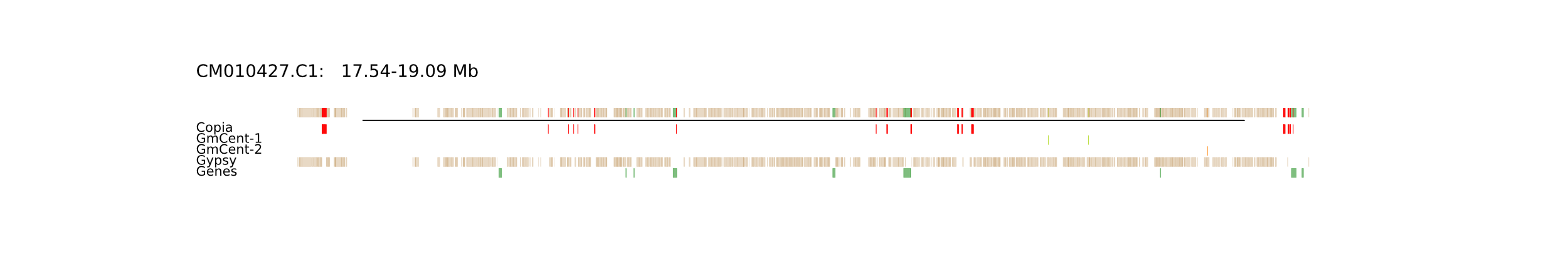

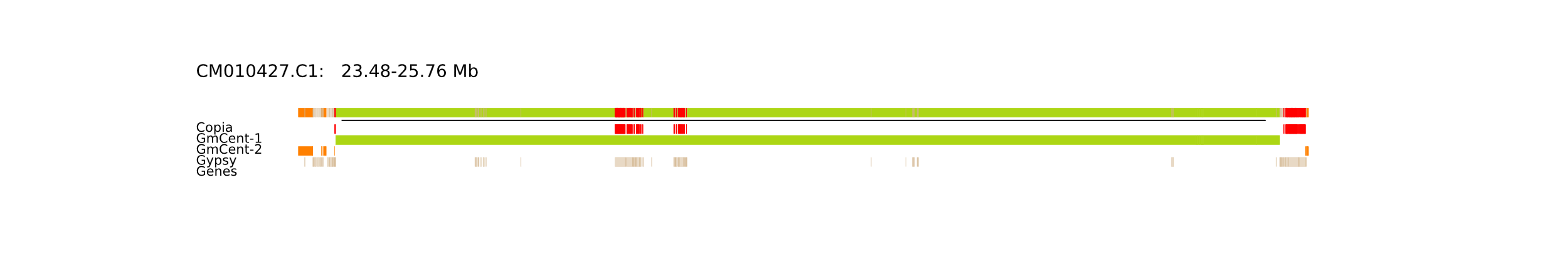

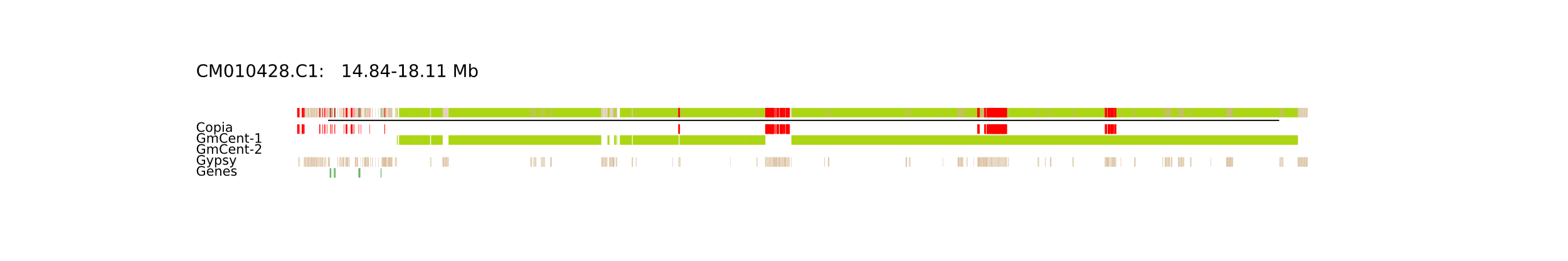

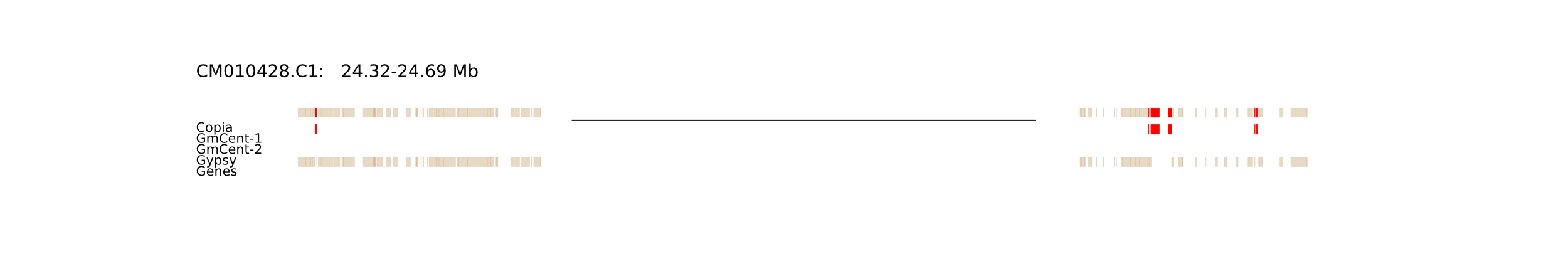

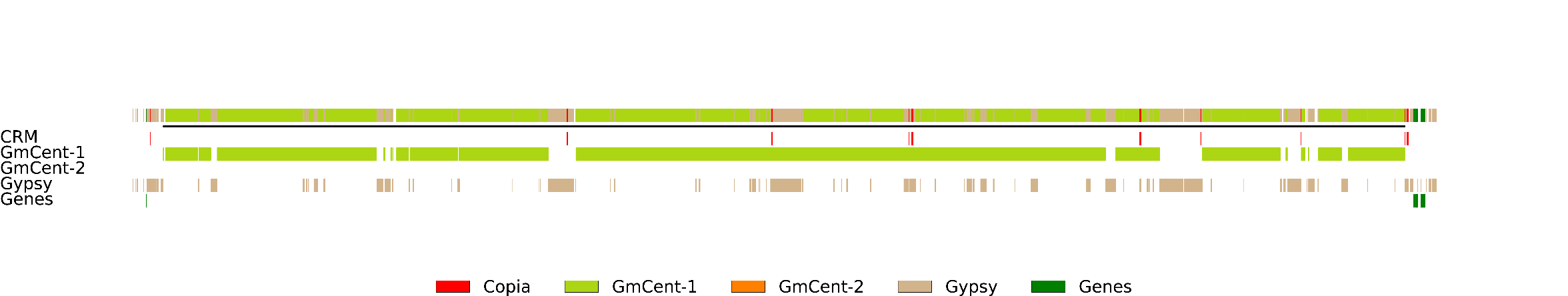

**Supplementary Figs. 5. The genomic structures of all Gaps.** **The gaps were represented by black lines.**

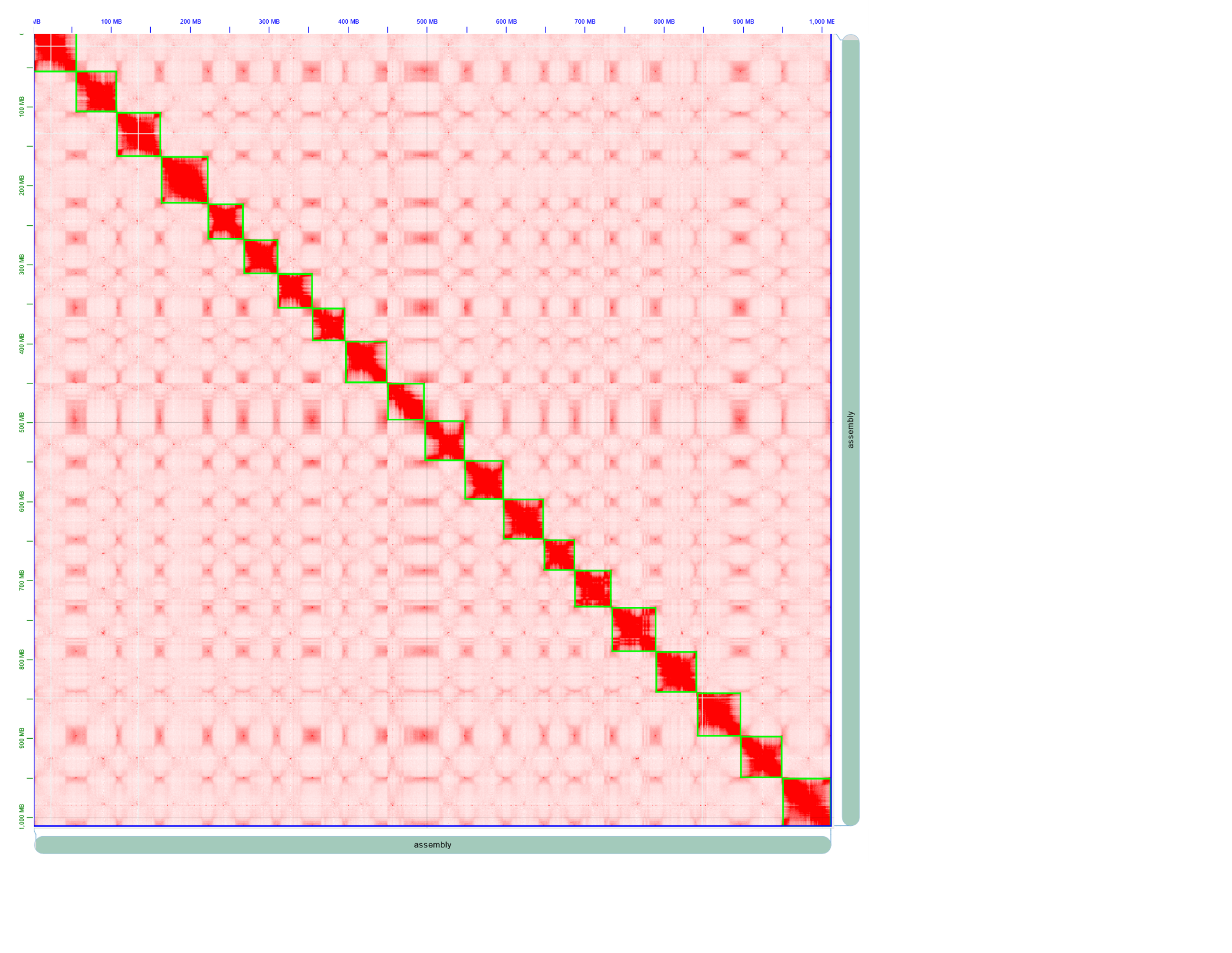

**Supplementary Fig. 6. Hi-C map of ZH13-T2T.**

The map was generated based on the alignments between the Illumina hi-C reads and ZH13-T2T with Juicerbox.

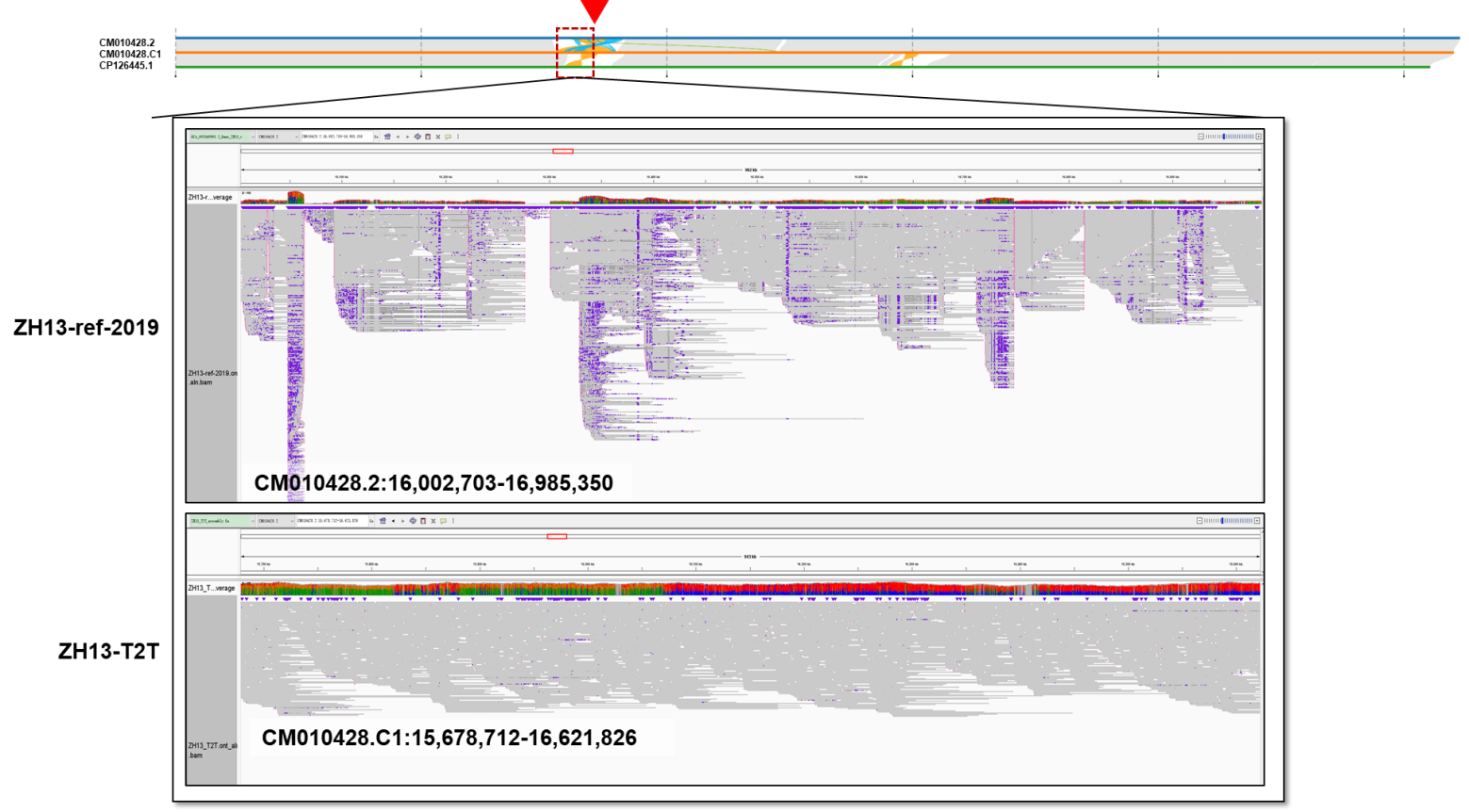

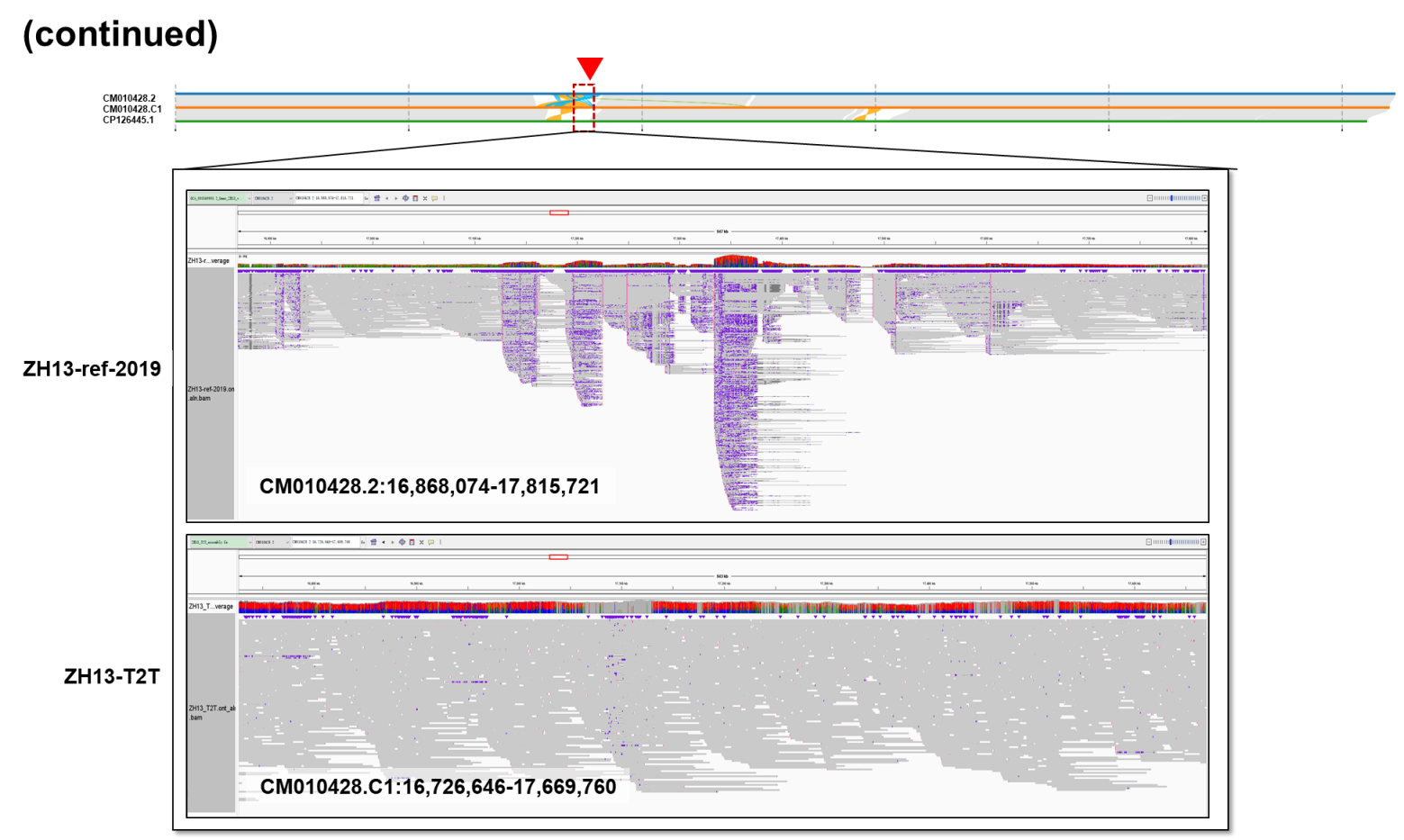

**Supplementary Fig. 7. Read alignment comparison between ZH13-ref-2019 and ZH13-T2T in an NAR with multiple SVs.**

The IGV snapshots of ONT ultralong read alignments in an NAR between ZH13-ref-2019 (CM010428.2:16,002,703-7,815,721) and ZH13-T2T (CM010428.C1:15,678,712-17,669,760) with complex SVs. There are plenty of read clippings, split alignments and high/low local coverages on ZH13-ref-2019, which suggest the disagreement between the local genomic sequences and reads. Meanwhile, the alignments on ZH13-T2T are consecutive with normal coverages along the region.

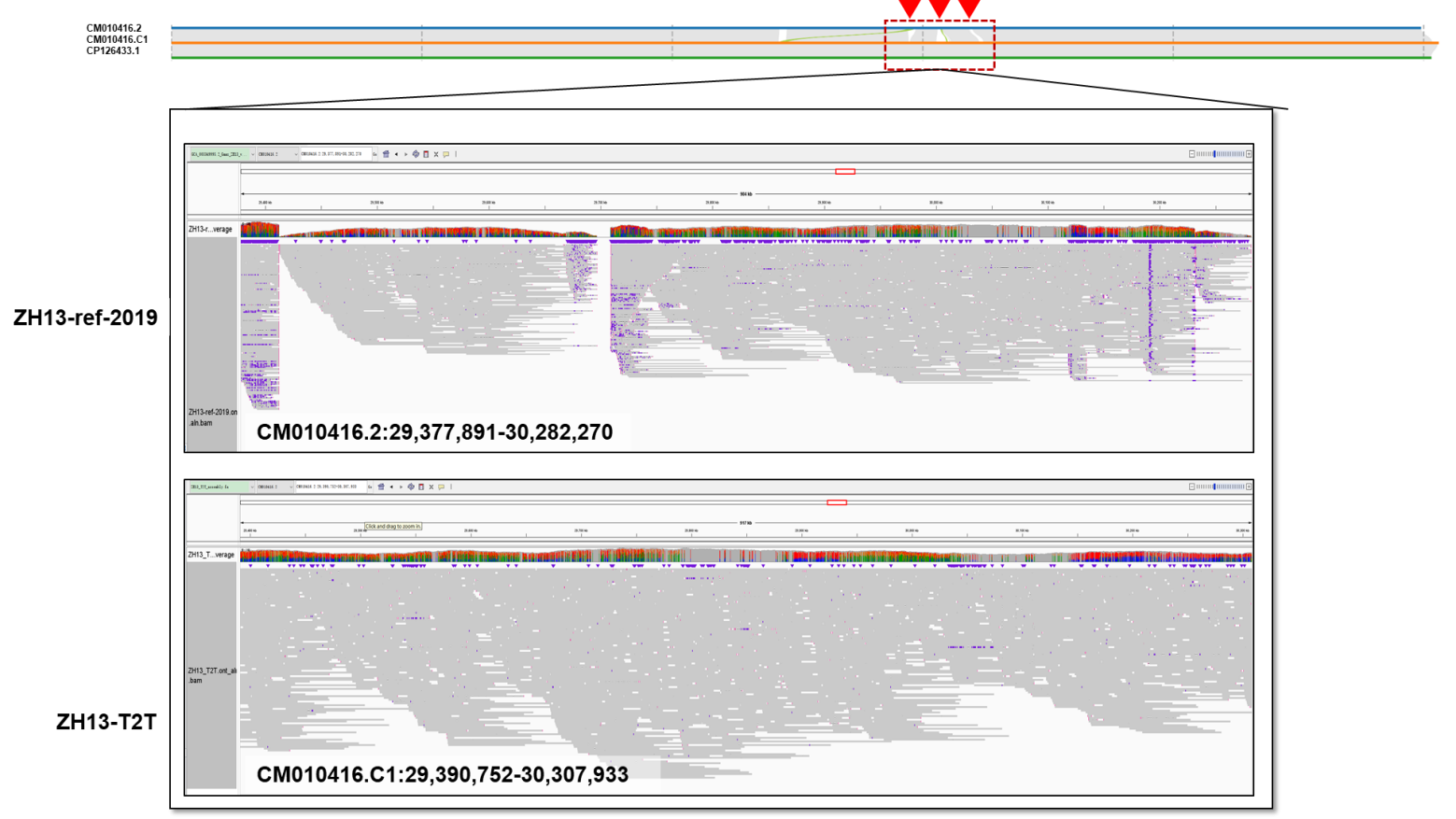

**Supplementary Fig. 8. Read alignment comparison between ZH13-ref-2019 and ZH13-T2T in an NAR without SV.**

The IGV snapshots of ONT ultralong read alignments in three NARs having no SV. The upper and lower parts are the alignments on ZH13-ref-2019 (CM010416.2:29,377,891-30,282,270) and ZH13-T2T (CM010416.C1:29,390,752-30,307,933). Concentrated read clippings are observed at the boundaries of the NARs from ZH13-ref-2019, but not for that of ZH13-T2T.

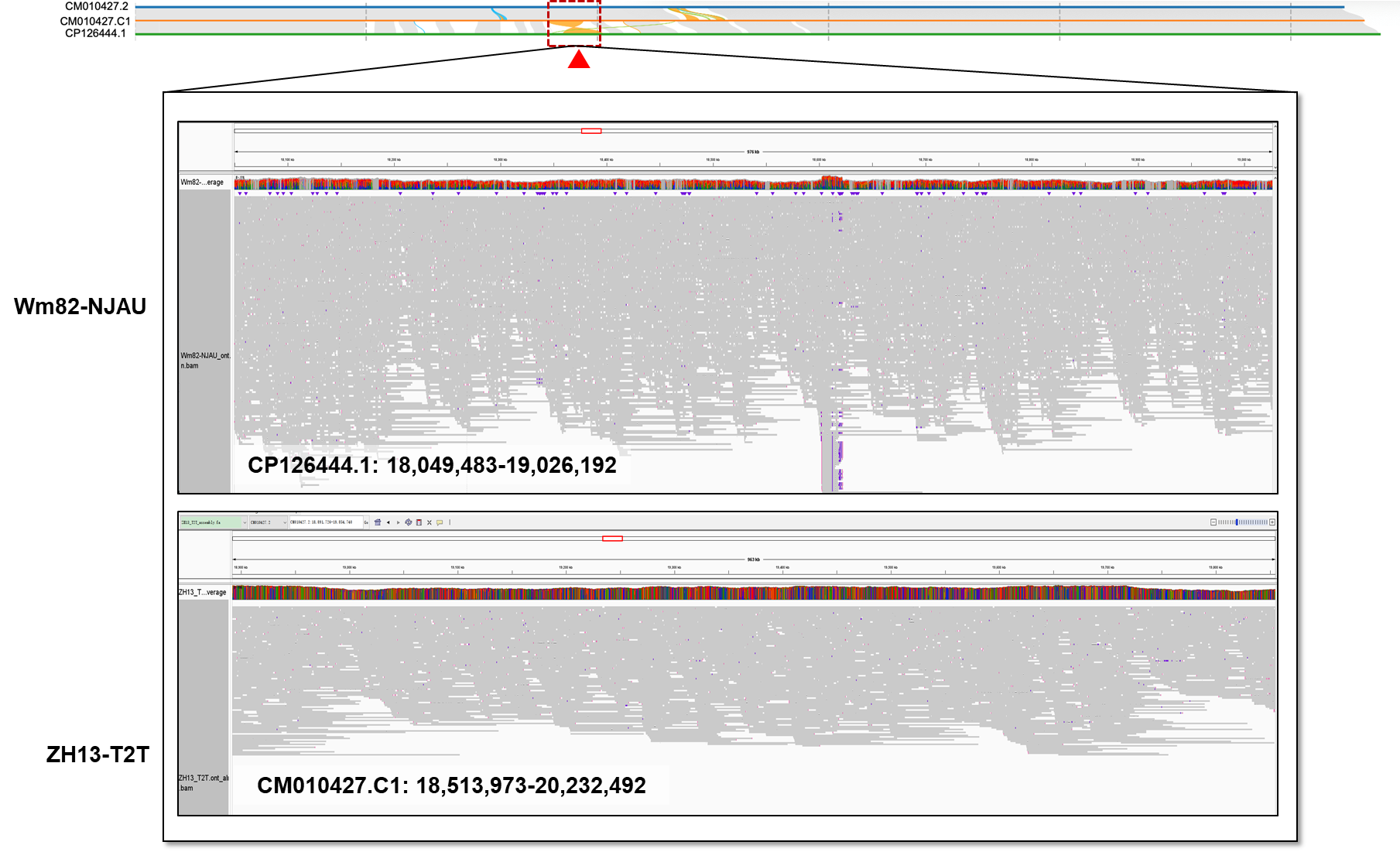

**Supplementary Fig. 9. Read alignment comparison between Wm82-NJAU and ZH13-T2T (case 1).**

The IGV snapshots of ONT ultralong read alignments in an NAR between Wm82-NJAU (CP126444.1:18,049,483-19,026,192) and ZH13-T2T (CM010427.C1: 18,513,973-20,232,492). There is no chimeric alignment concentrated in both of the two corresponding regions. Thus, the NAR is plausibly due to the SV between the two genomes.

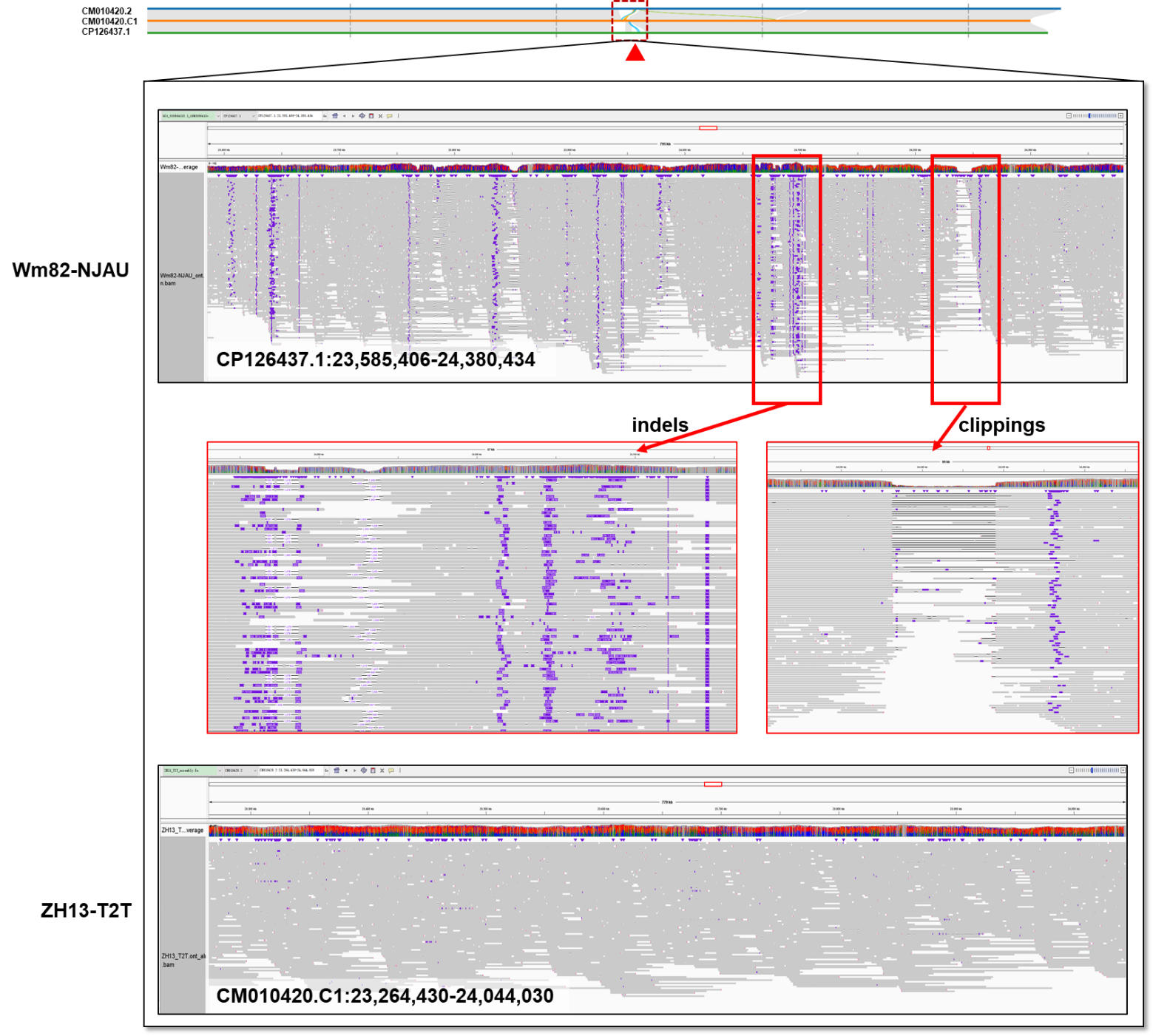

**Supplementary Fig. 10. Read alignment comparison between Wm82-NJAU and ZH13-T2T (case 2).**

The IGV snapshots of ONT ultralong read alignments in an NAR between Wm82-NJAU (CP126437.1:23,585,406-24,380,434) and ZH13-T2T (CM010420.C1:23,264,430-24,044,030). In the nearly 1 Mbp region, there are multiple sites having concentrated large indel and read clipping signatures in Wm82-NJAU. Meanwhile, no such signature is observed from ZH13-T2T.

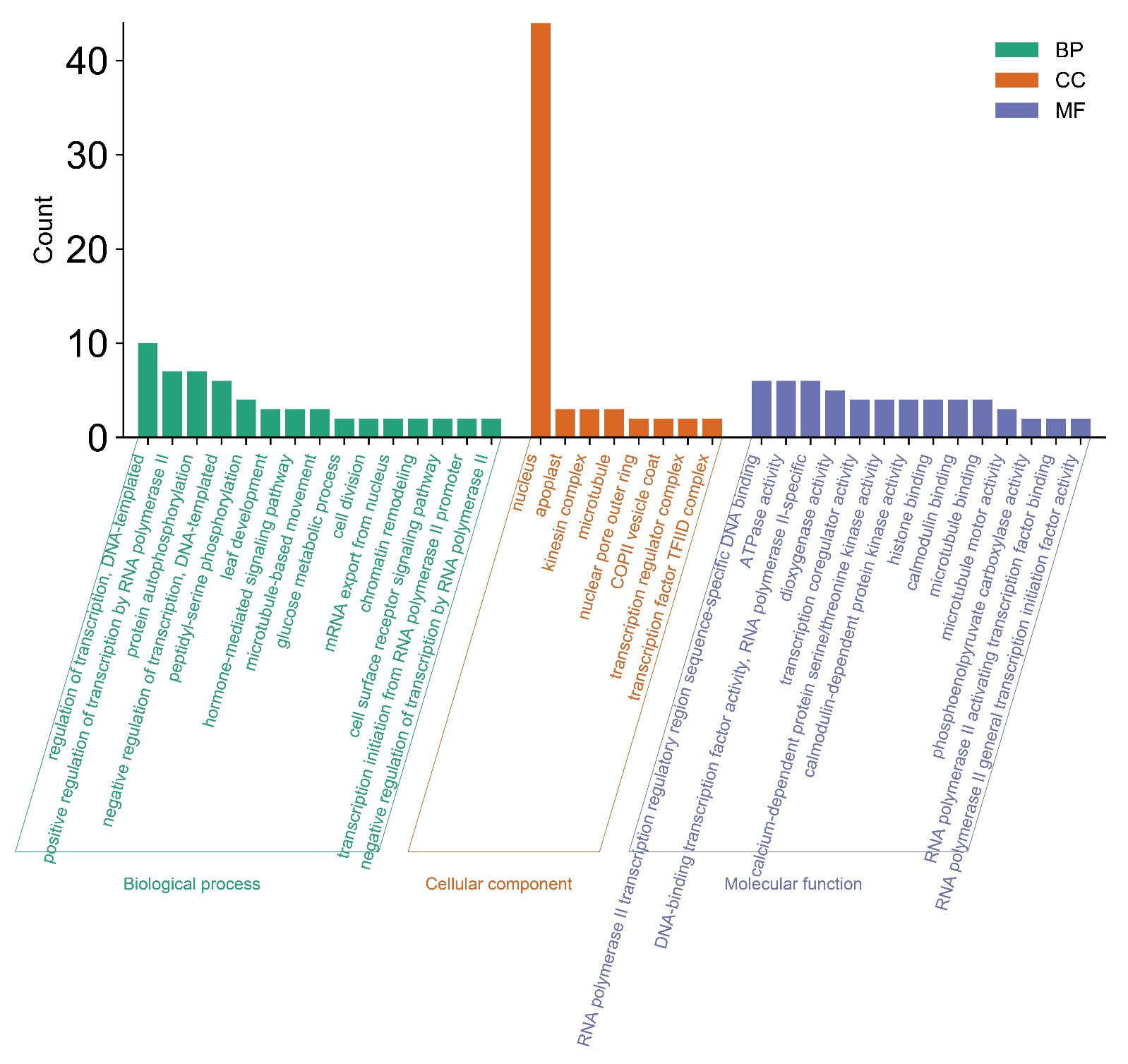

**Supplementary Fig. 11. GOterm enrichment analysis results of the newly identified genes in terms of biological processes, cellular components and molecular functions.**

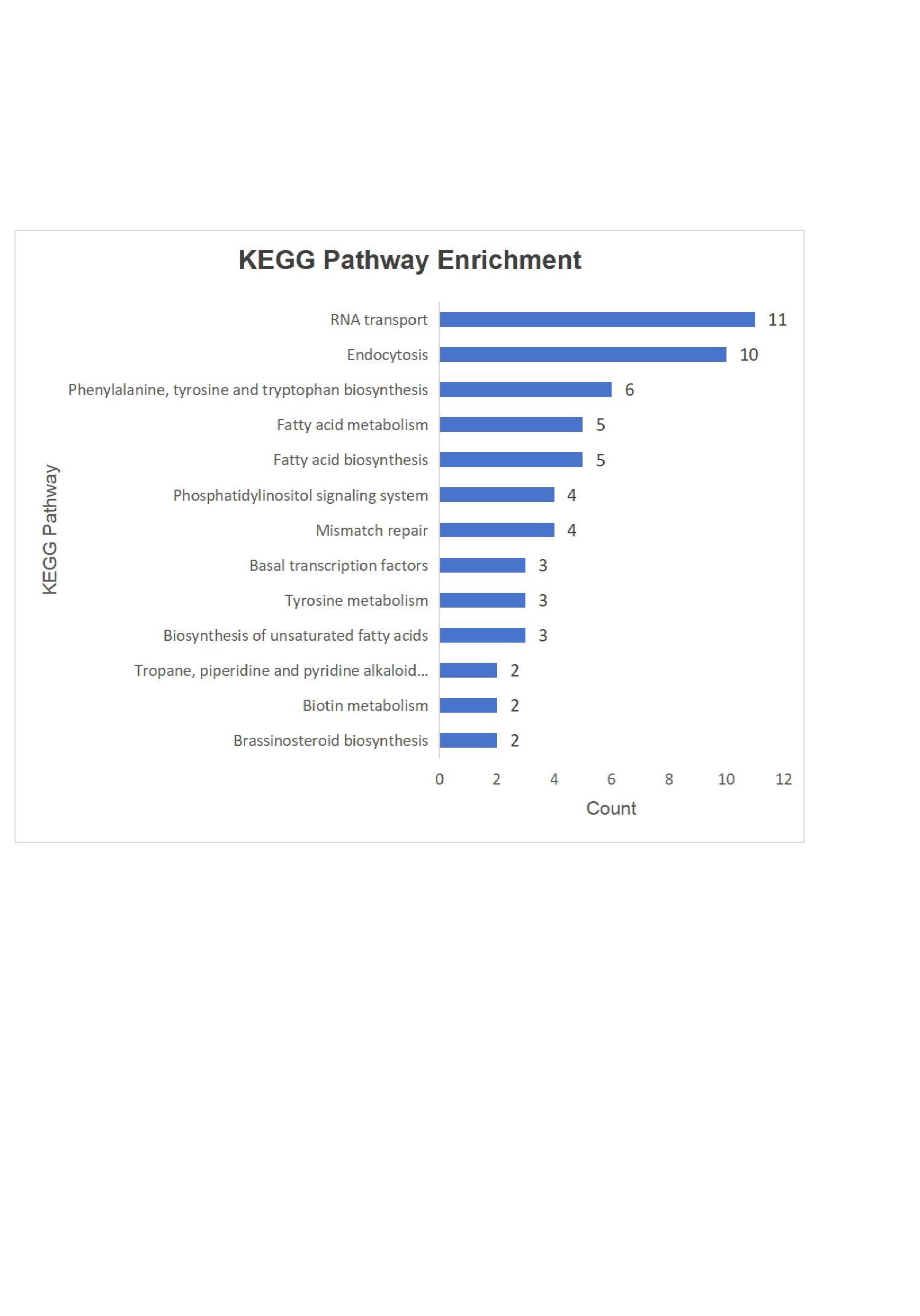

**Supplementary Fig. 12. Results of enrichment analysis of newly discovered genes on KEGG pathways.**

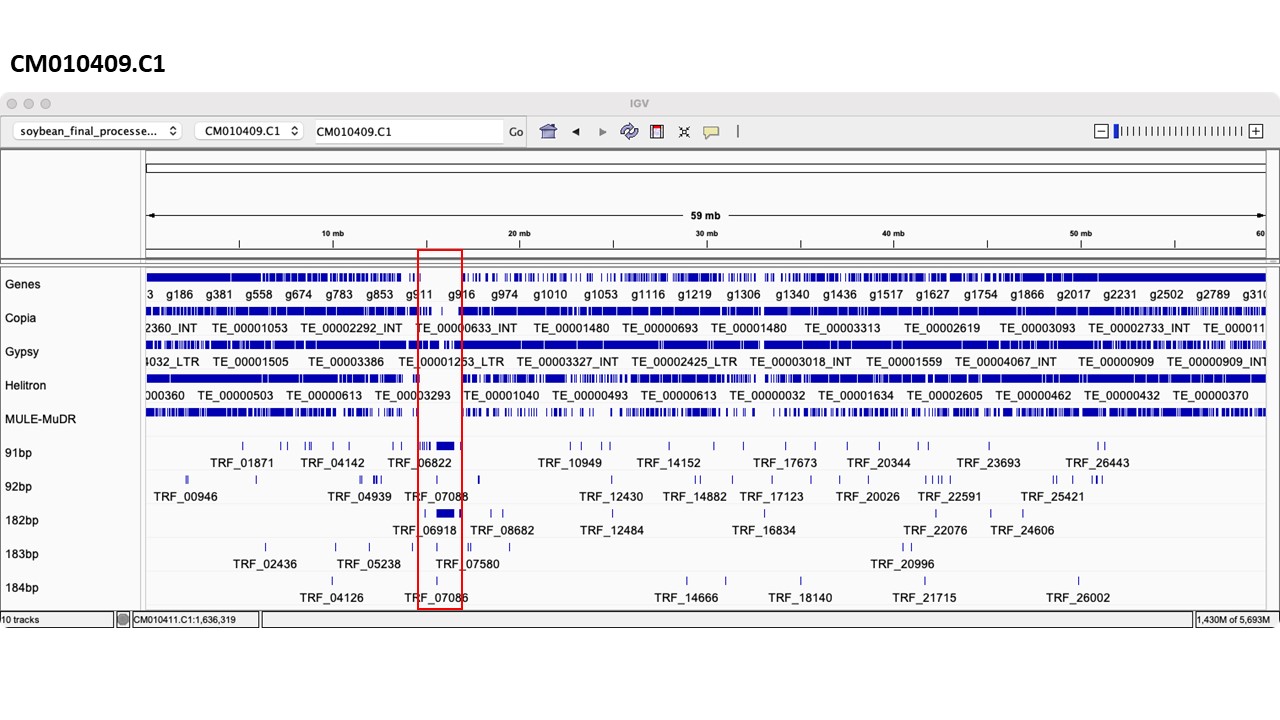

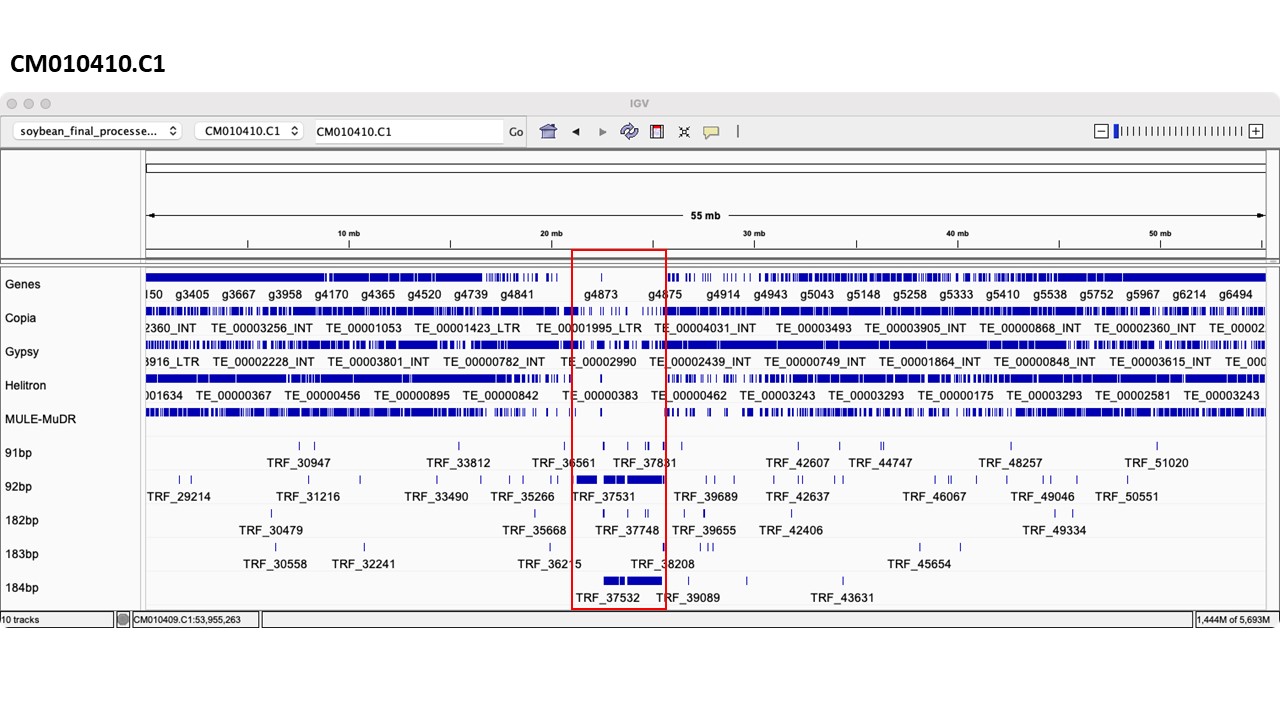

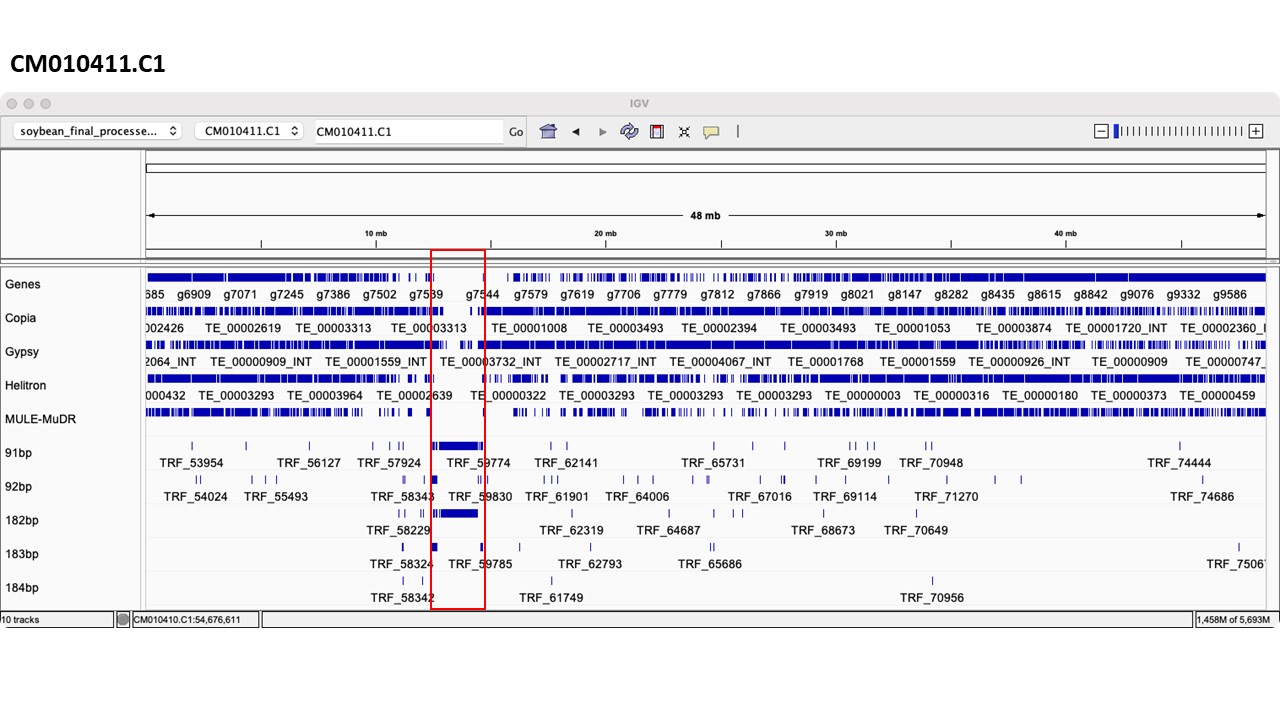

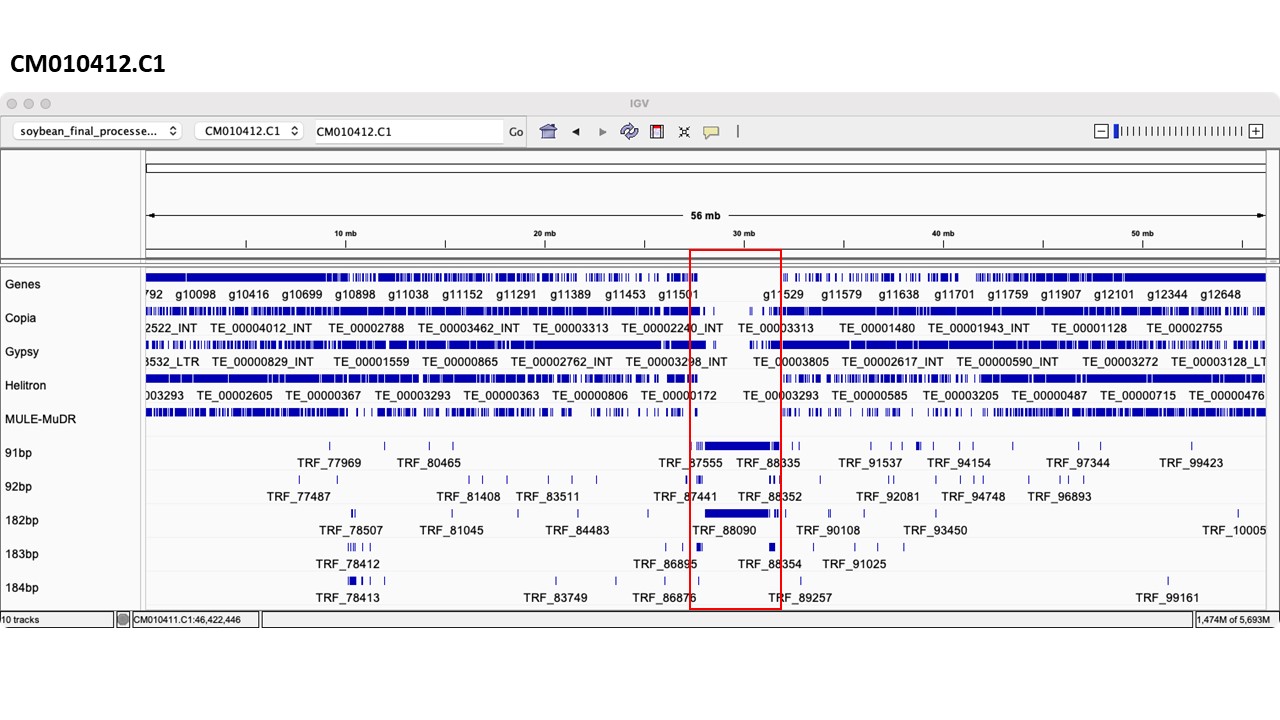

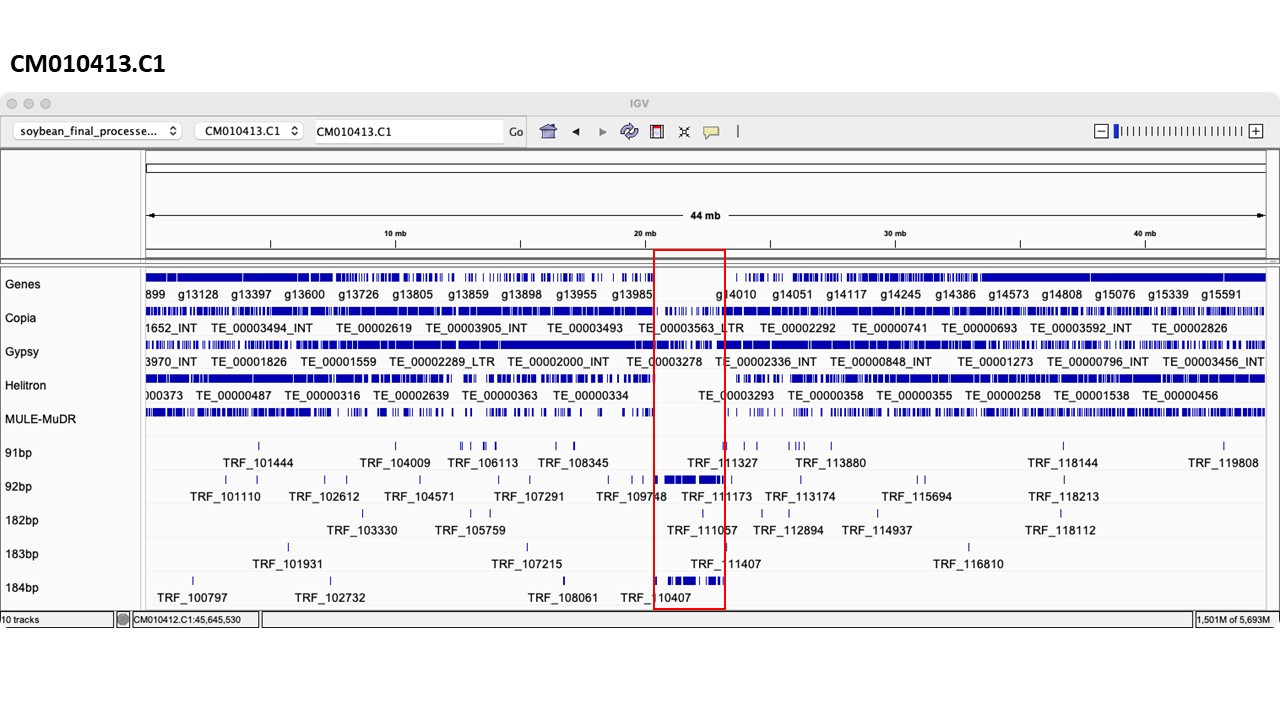

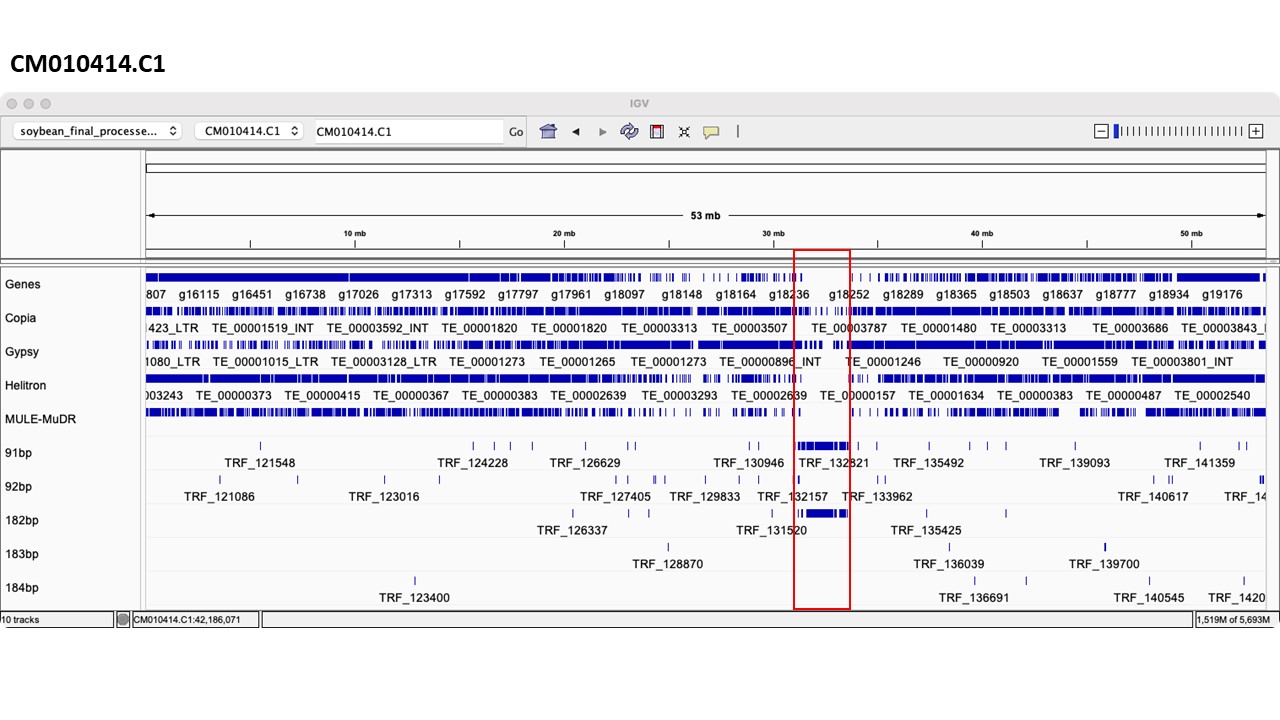

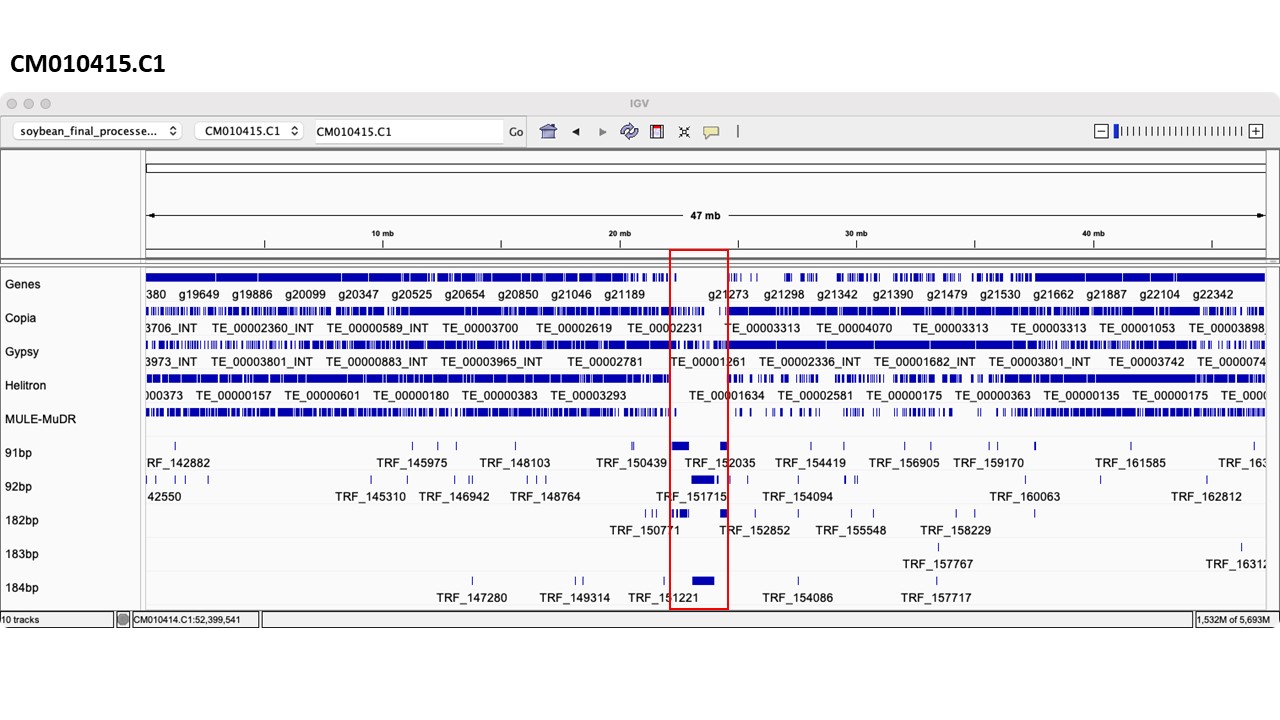

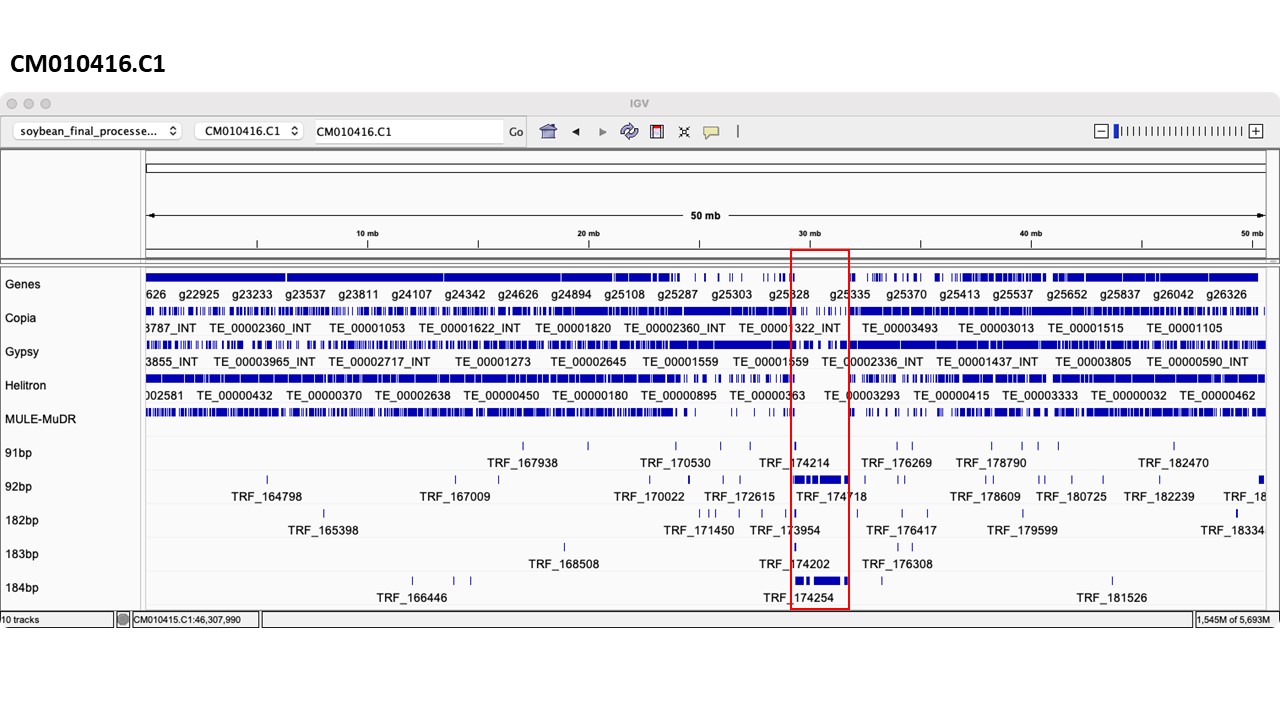

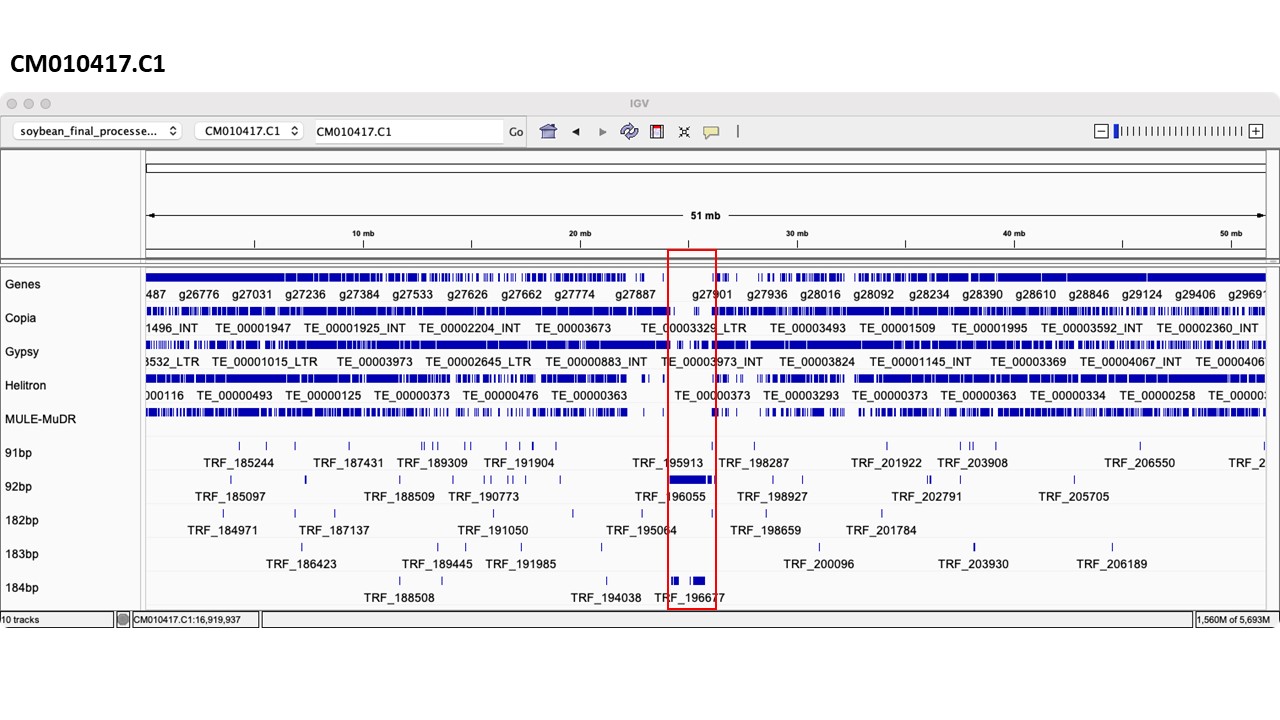

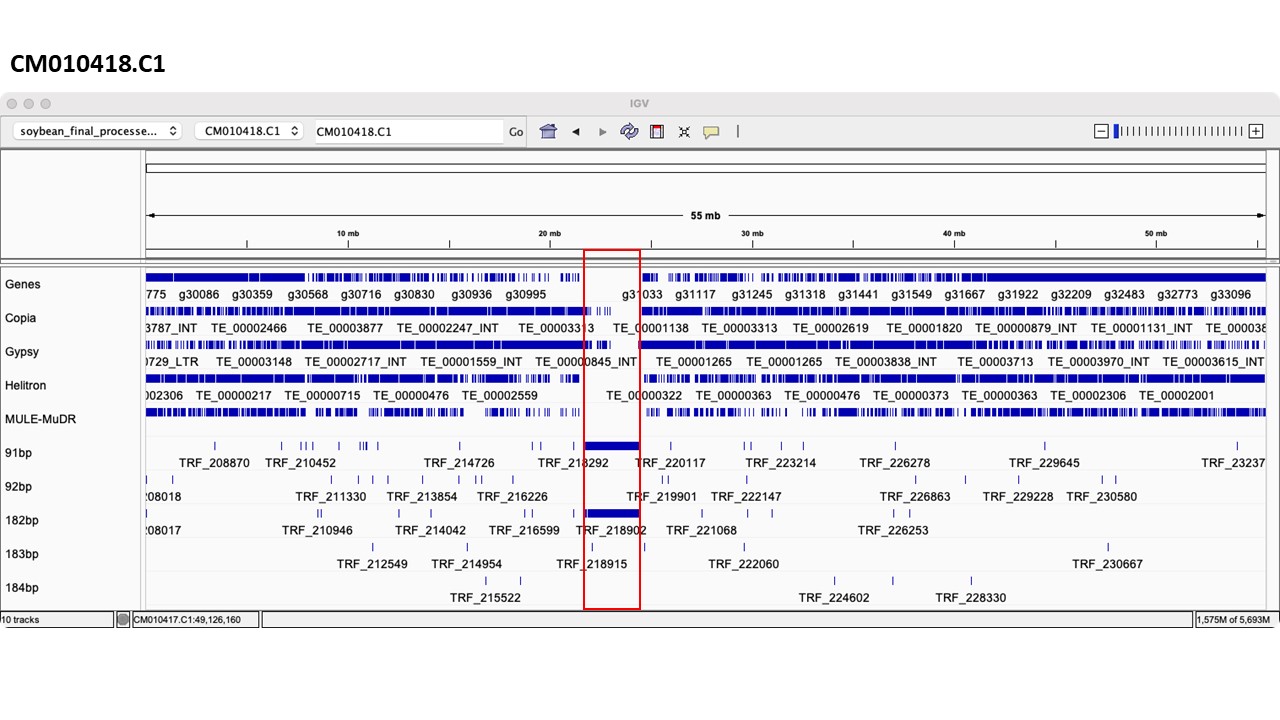

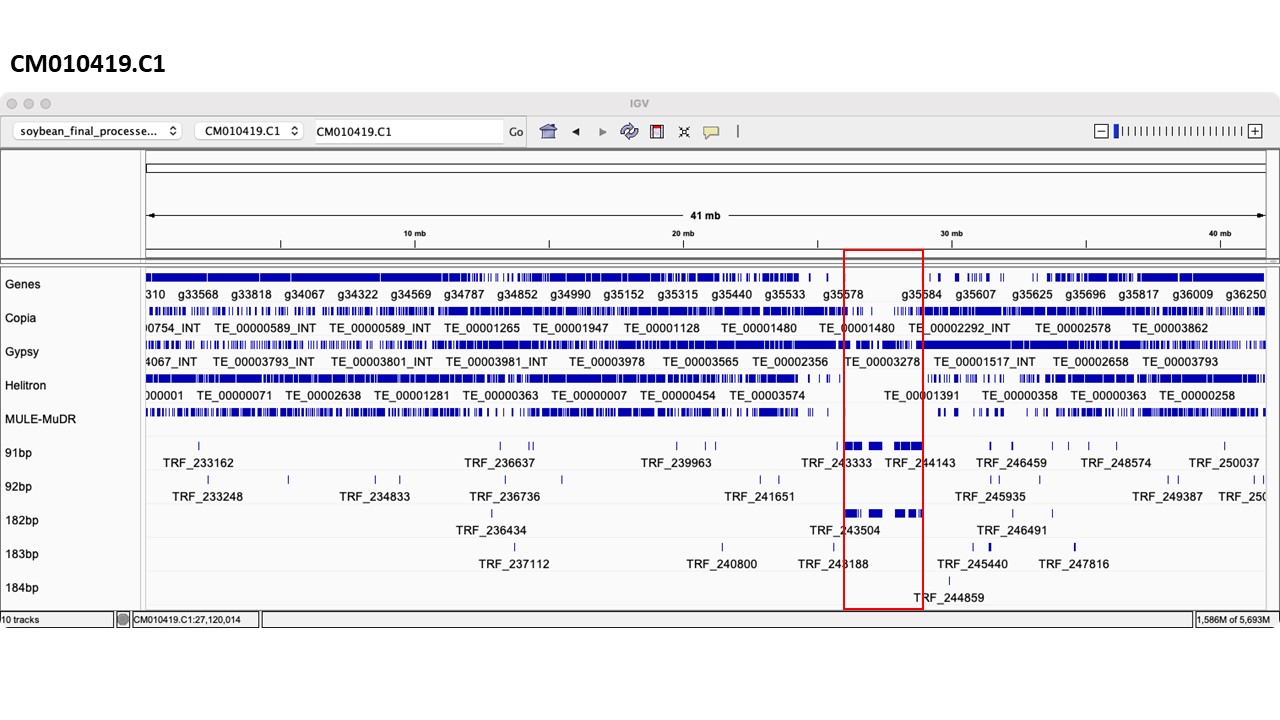

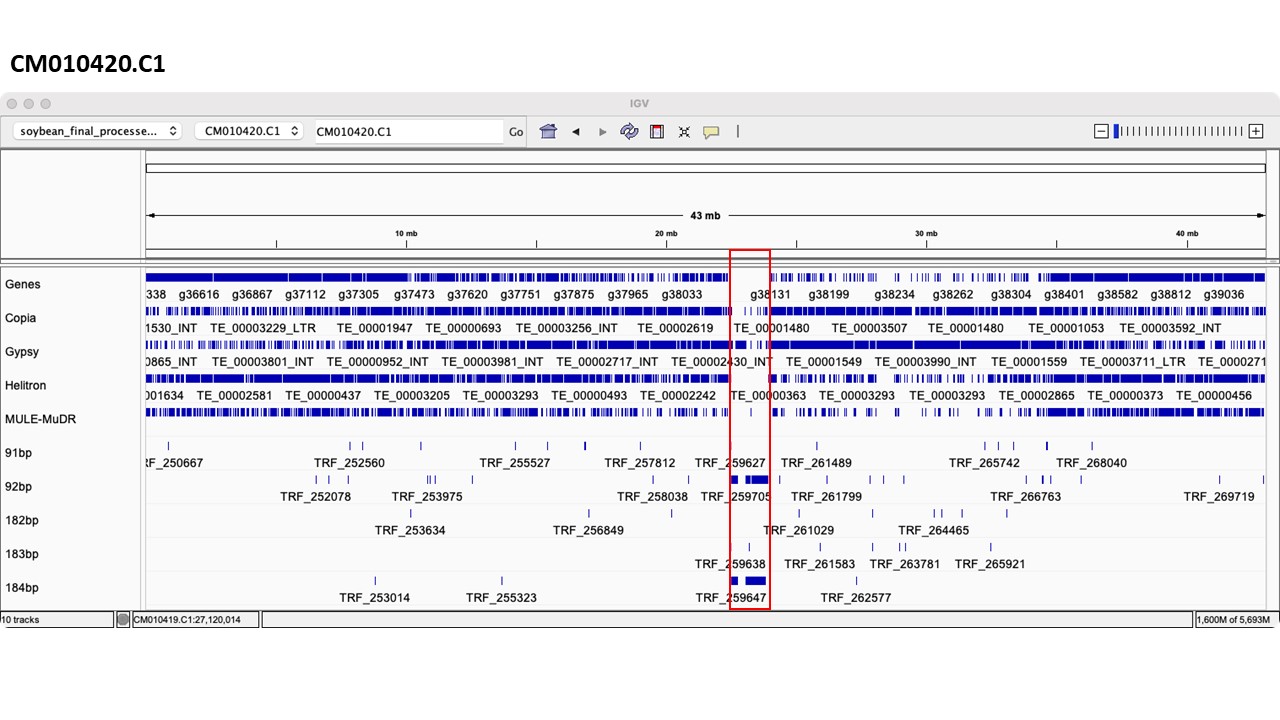

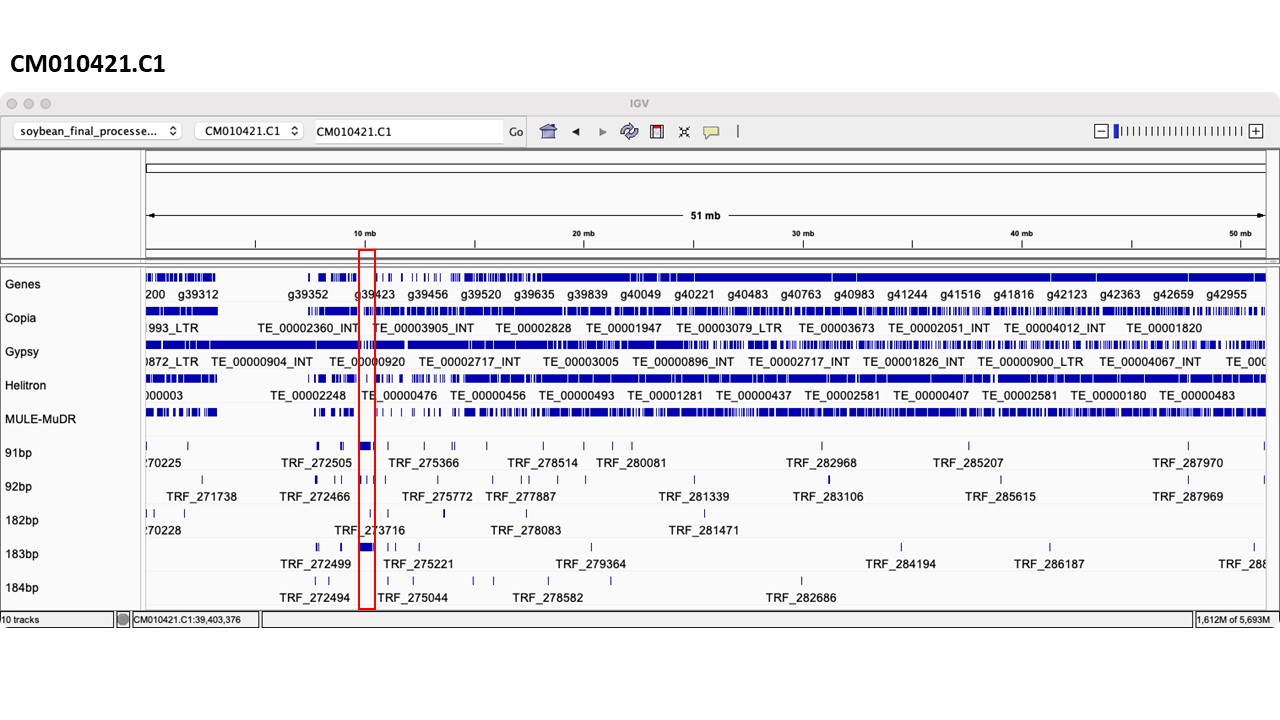

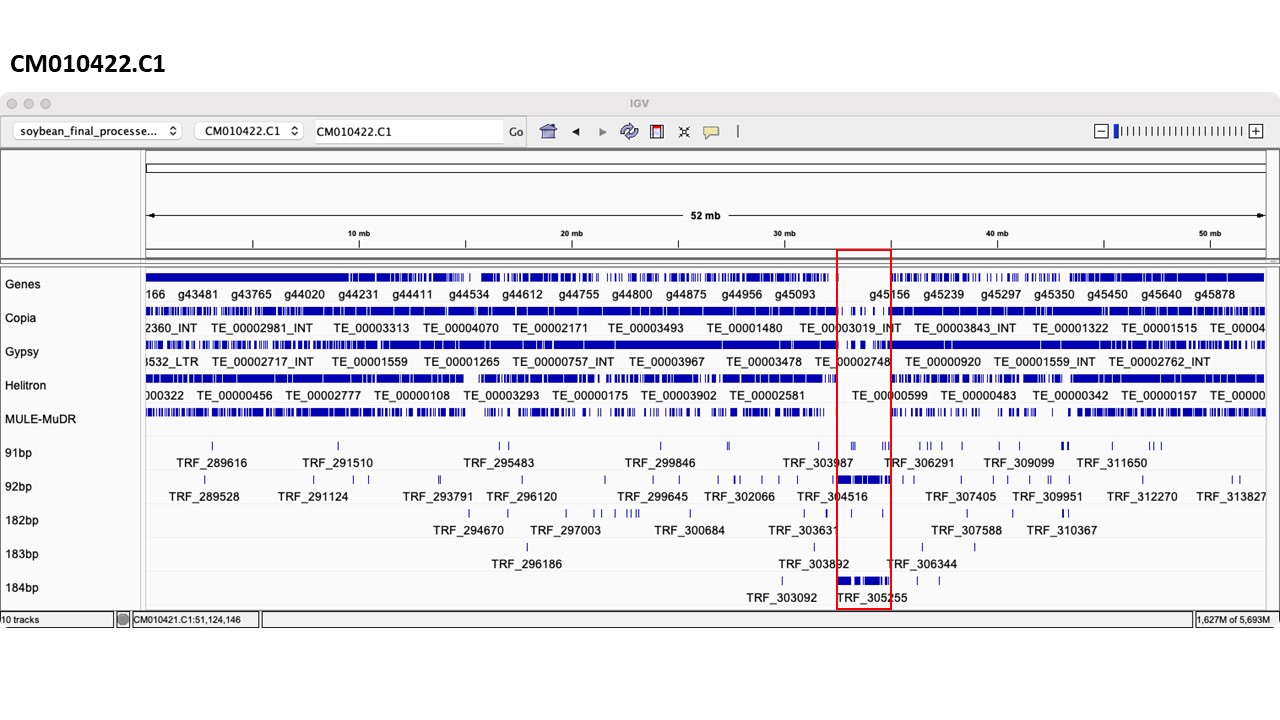

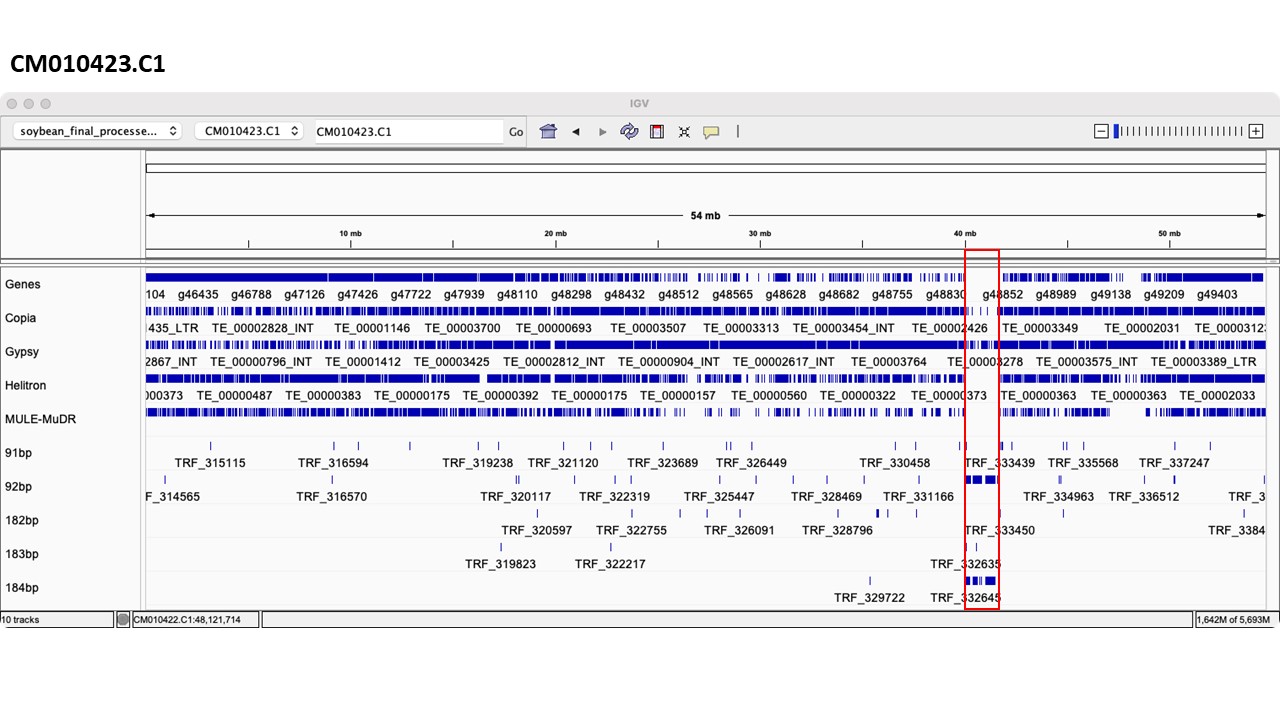

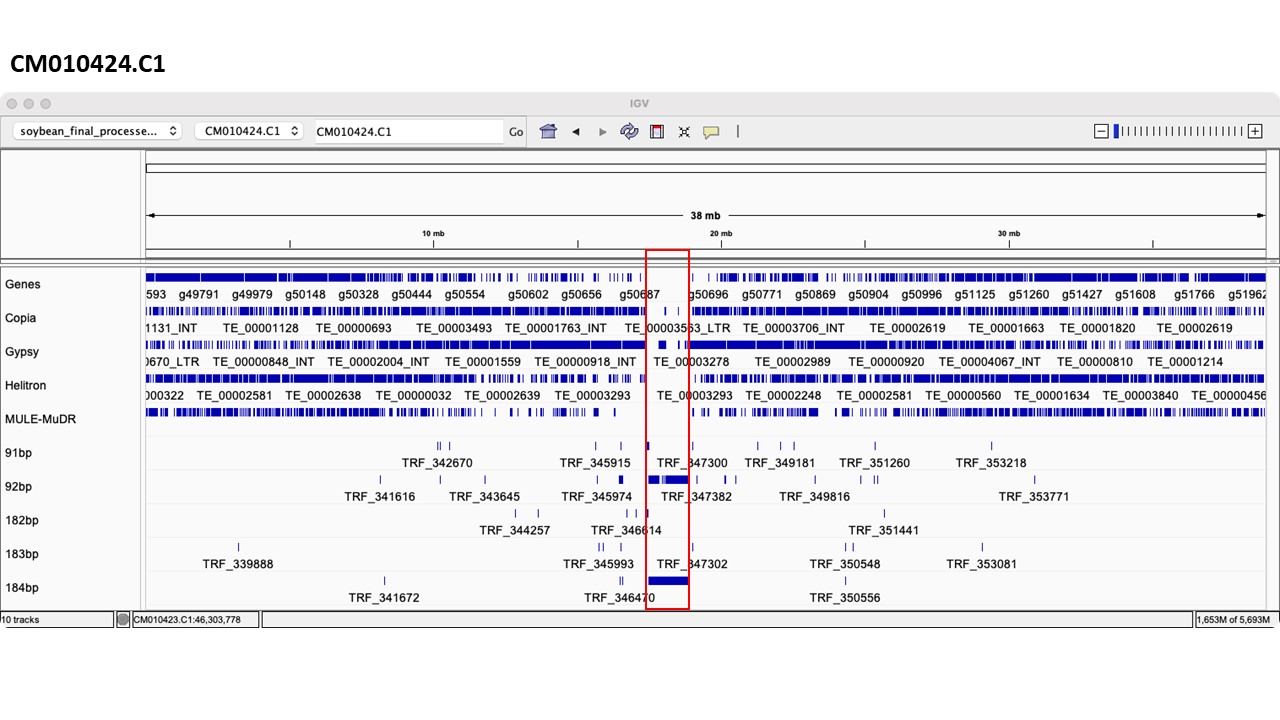

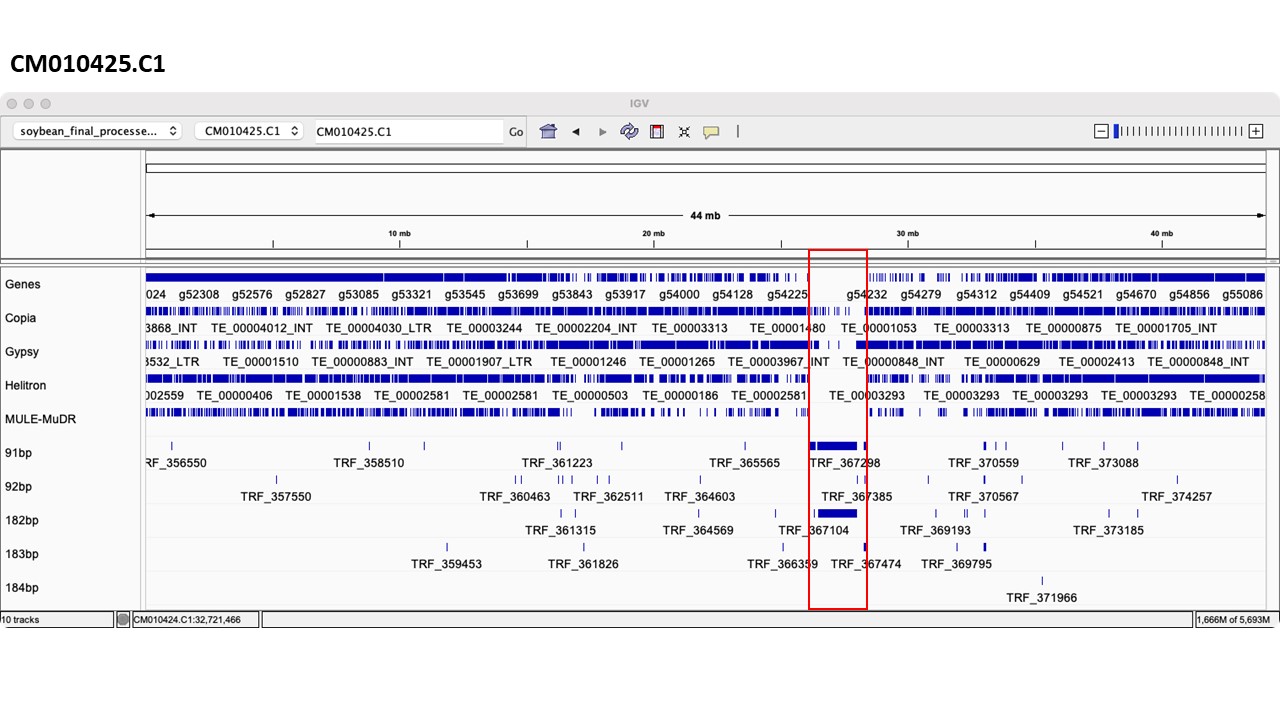

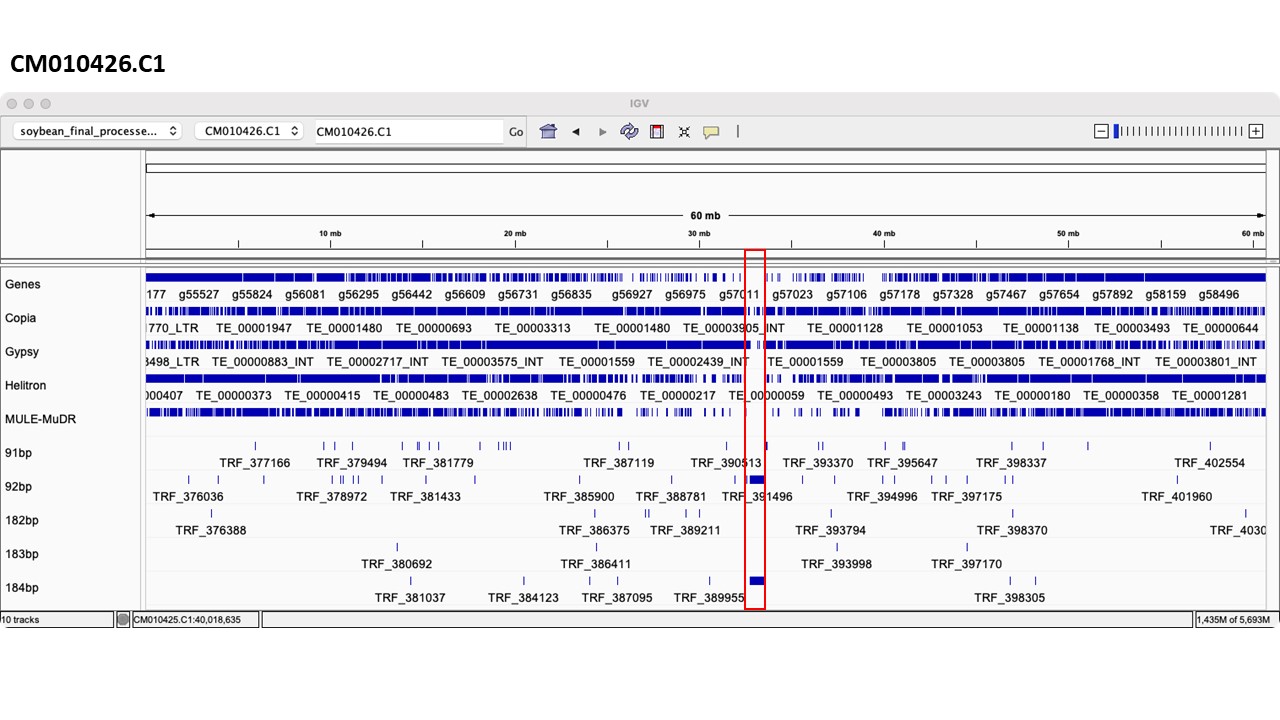

**Supplementary Fig. 13. Identification of centromere**

**Supplementary Fig. 14. Genome structure of centromere region.** The centromeres were represented by black lines.
